# *ATG* gene duplication in vertebrates: evolutionary divergence and its functional implications

**DOI:** 10.1101/2025.10.24.684263

**Authors:** Sidi Zhang, Ikuko Koyama-Honda, Daiki Hiratsuka, Noboru Mizushima

**Affiliations:** Department of Biochemistry and Molecular Biology, Graduate School of Medicine, The University of Tokyo, Tokyo, Japan; Department of Obstetrics and Gynecology, Graduate School of Medicine, The University of Tokyo, Tokyo, Japan

**Keywords:** *ATG* genes, evolutionary fate, functional difference, gene duplication, ohnolog, paralog, vertebrates, whole-genome duplication

## Abstract

Macroautophagy (hereafter referred to as autophagy) requires the coordinated action of approximately 20 autophagy-related (*ATG*) genes. Duplication of *ATG* genes has had a major impact on the evolution of the autophagy pathway among major lineages. One duplication hotspot is in vertebrates. However, the exact duplication timing, post-duplication evolutionary divergence patterns, and its relation to functional differences among paralogs have not been investigated in detail. Here, we demonstrate that most *ATG* genes were likely duplicated by whole-genome duplication events near the root of vertebrates. We compared the sequence and gene expression divergence between paralogs and categorized the evolutionary fates (i.e., how ancestral function is divided between paralogs). Within the paralog pairs that evolved most asymmetrically, namely *BECN, WIPI* (*WIPI1* and *WIPI2*), and *ATG16*, one paralog likely retained the ancestral function, allowing the other to evolve under less constraint. While no obvious asymmetry was observed between *ATG9A* and *ATG9B* in non-mammalian vertebrates, *ATG9B* experienced marked sequence divergence and expression level reduction in mammals, suggesting a shift in balance. Expression patterns among the *ULK-1* (*ULK1* and *ULK2*), *GABARAP* (*GABARAP* and *GABARAPL1*), and *LC3* (*LC3A* and *LC3B*) pairs were more consistent with hypofunctionalization/dosage sharing, such that ancestral function depends on both paralogs. We also demonstrate that both *ULK1* and *ULK2* can support autophagy, whereas only *BECN1*, but not *BECN2*, has autophagic function and discuss the relationship between autophagic function and evolutionary divergence between paralogs. The present detailed analysis of *ATG* gene duplication in vertebrates provides a critical time-line for interpreting functional differentiation between homologs.

## Introduction

Gene duplication, whether through whole-genome or smaller-scale duplications, is a major source of evolutionary innovation [1]. Following gene duplication, the fates of the duplicated pairs can generally be classified into one of the following categories [2-5]: (i) most commonly, perhaps ironically, the restoration of the singleton state by gene loss; (ii) the ancestral function being maintained in one paralog, while the other evolves under less constraint, sometimes leading to neofunctionalization; (iii) subfunctionalization, in which each paralog maintains a subset of ancestral functions; (iv) hypofunctionalization or dosage sharing, in which the expression level of each paralog is reduced such that expression from both paralogs is necessary for ancestral function; (v) outcomes related to other dosage-related mechanisms, such as positive dosage (increased gene expression from gene duplication being beneficial) and dosage balance (the need to maintain stoichiometry between complex-forming genes being a major determinant of this fate).

Macroautophagy (hereafter referred to as autophagy) is a process that captures and delivers cellular components to the lysosome/vacuole for degradation. Autophagy requires the coordinated action of the core autophagy-related (*ATG*) genes from approximately 20 gene families [6], which, through gene duplications, have expanded to around 37 genes in mammals. Apart from a few ancient events that likely date back to the eukaryotic or metazoan common ancestor, namely the formation of *ULK3* (unc-51 like kinase 3), *ULK4* (unc-51 like kinase 4), and *STK36*/*Fused* (serine/threonine kinase 36) [7,8], *WDR45B* and *WDR45* (WD repeat domain 45B and 45) [9,10], and *MAP1LC3C* (microtubule associated protein 1 light chain 3 gamma; shortened to *LC3C* hereafter) and *GABARAPL2* (GABA type A receptor associated protein like 2) [11,12], many of these duplication events presumably occurred in vertebrates [13]. Although many comparative genomic and transcriptomic studies have analyzed duplicated genes in vertebrates or mammals on the genome-wide scale [14-16], no study has focused specifically on the *ATG* genes (e.g., with more detailed orthology checks and tailored analyses). Consequently, neither the exact duplication timing of the *ATG* genes nor their evolutionary fates after duplication is clear.

Another topic closely related to the evolutionary fate of the *ATG* genes after duplication is their functional differences. Many (mostly experimental) studies have investigated the functional differences among *ATG* homologs, with the ATG8 and ATG4 families being the primary focus [17-23], but other families were also studied [7,24-28]. However, because these studies are often restricted to a single cell type or condition in humans, it remains unclear when the observed differences first evolved and whether any functional changes occurred during their evolution. Evolutionary analysis, while not directly gauging gene functions, offers another dimension that complements the published functional analyses.

Here, we identified the most likely timing of duplication of the core *ATG* genes in vertebrates, analyzed the post-duplication evolutionary divergence pattern, and discuss the relationship between evolutionary and functional divergence using experimental data from previous studies and newly generated in the present study. Our research reveals the post-duplication evolutionary dynamics of the *ATG* genes in vertebrates and provides a timeline for the emergence of functional differences between paralogs.

## Results

### Most ATG genes were likely duplicated during whole-genome duplication events near the root of vertebrates

Before analyzing the evolutionary dynamics of the *ATG* genes after duplication, we first determined the timing of their duplication. To this end, we selected 30 species in Chordata with highquality genome assemblies that are representative of the major lineages (including early-diverging vertebrate groups, the cyclostomes and cartilaginous fish) as well as five outgroup species (Table S1), and searched for homologous sequences in their proteomes using the human homolog sequences as queries. Based on the distribution of *ATG* homologs, the set of genes possessed by the common ancestor of Chordata and the most likely timings of the primary duplication events (i.e., the earliest events in vertebrates that gave rise to the major paralogs, e.g., duplication of *ULK-1* into *ULK1* and *ULK2* [unc51 like autophagy activating kinase 1 and 2]) were determined (Figure 1). Among the core *ATG* genes, *ULK-1, ATG9, BECN* (beclin), *ATG2, WIPI* (WD repeat domain, phosphoinositide interacting), *GABARAP* (GABA type A receptor-associated protein), *LC3, ATG4-1, ATG4-2*, and *ATG16* were inferred to have been duplicated in Chordata (i.e., there was only one copy in the chordate ancestor). Although most of these genes were duplicated in Gnathostomata, *GABARAP* and *BECN* were duplicated at later times (in the common ancestor of bony vertebrates and eutherians, respectively). We also identified more lineage-specific duplication and loss events (only those supported by more than one species are shown in Figure 1).

**Figure 1.**
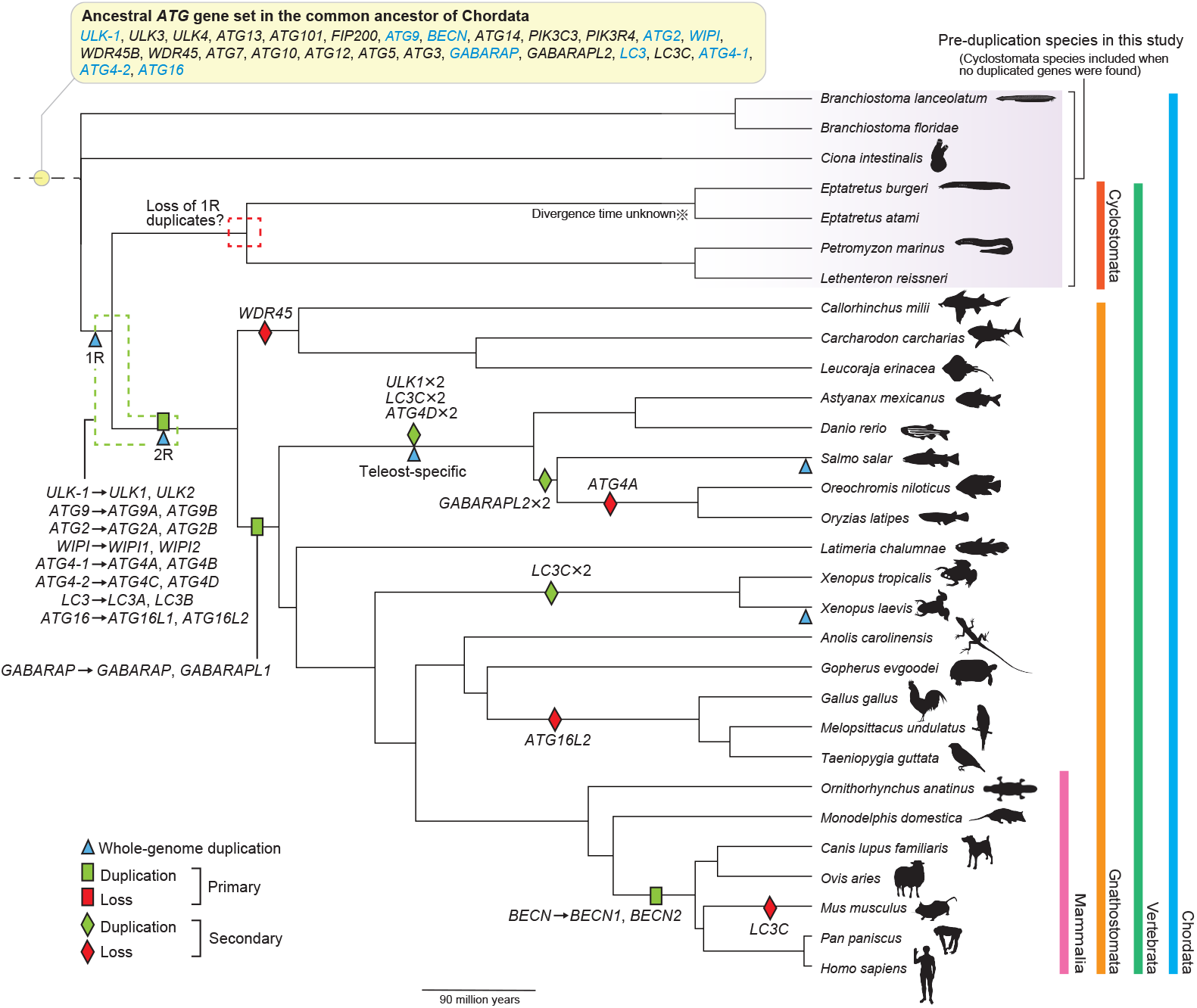
Most *ATG* genes were duplicated during whole-genome duplication events (1R and 2R) near the root of vertebrates. In the top panel, the set of *ATG* genes possessed by the common ancestor of Chordata is listed, and genes that duplicated in vertebrates are shown in cyan text. The species that diverged before duplication (i.e., pre-duplication species), whose sequences or gene expression levels were used as proxies for the ancestral levels, are shaded in purple. Inferred gene duplication (green) and loss events (red) in this study, as well as known whole-genome duplication events (blue), are labeled on a time-calibrated species tree from TimeTree (the divergence time between *Eptatretus burgeri* and *Eptatretus atami* is unknown; events found only in one species here are not shown; the relative positions of the labels on each branch are arbitrary). The events leading to the formation of the major *ATG* gene groups are denoted as “primary,” while more lineage-specific events are denoted as “secondary.” The silhouette images (all in the public domain) were downloaded from PhyloPic.

Two rounds of whole-genome duplications (1R and 2R, respectively) occurred in the ancestral vertebrate lineage [1,29]. Recent studies have suggested that 1R occurred in an ancestor of cyclostomes and gnathostomes, while 2R was specific to gnathostomes [30-33] (blue triangles in Figure 1). Three additional whole-genome duplication events in ancestors to the teleost fish, salmonids, and *Xenopus laevis* also occurred (Figure 1; blue triangles). The inferred timing of the primary duplication events of the core *ATG* genes (other than *GABARAP* and *BECN*) coincides with that of 2R, suggesting that they may be ohnologs (i.e., genes duplicated via the vertebrate whole-genome duplications, named in honor of Susumu Ohno). When we cross-checked three published lists of ohnologs, two of them consistently supported the ohnolog status of most *ATG* genes [34,35], while a third list supported none of them as being ohnologs [36] (Table S2). Because these lists were created by analyzing synteny blocks in the genomes, they provide strong evidence for gene duplications by whole-genome duplications. Note that one list also classified *GABARAP* and *GABARAPL1* (GABA type A receptor-associated protein like 1) as ohnologs [34], and we do not reject the possibility that they were first duplicated by whole-genome duplication but then lost in the cartilaginous fish lineage. Collectively, these findings indicate that, other than *GABARAP* and *BECN*, most *ATG* genes present as multiple copies in vertebrates were likely duplicated during whole-genome duplication events.

To confirm the homology and clarify their evolutionary history, we reconstructed phylogenetic trees for each *ATG* gene family that duplicated in vertebrates (Figure S1). Long branches in the phylogenetic trees were observed for *ATG9B* in mammals (Figure S1B) and *GABARAPL1* in the teleost fish (Figure S1Fi), suggesting accelerated evolution occurred in these lineages. In the *BECN* tree, the long branch represents the emergence of *BECN2* (beclin 2) in eutherian mammals (Figure S1C). For *ATG2A*, the branch leading to XP_033927741.1, XP_033927739.1, and XP_033927740.1 in *Melopsittacus undulatus* and XP_041567864.1 in *Taeniopygia guttata* is long because these sequences are short (incomplete) (Figure S1D). Because of their incomplete sequences, these two species were removed from the sequence divergence analysis below (see next section).

Thus, most of the *ATG* genes that duplicated in vertebrates (*ULK-1, ATG9, ATG2, WIPI, LC3, ATG4-1, ATG4-2*, and *ATG16*) likely did so as part of the whole-genome duplications occurring near the vertebrate common ancestor, whereas *GABARAP* and *BECN* duplicated at later times.

### Paralogs within the BECN, WIPI, GABARAP, and ATG16 pairs and ATG9 in mammals diverged highly asymmetrically at the sequence level

Many previous studies (mainly in mammals) have reported that sequence evolution is asymmetric following gene duplication (i.e., one copy, usually the derived copy [the copy at a new genome location], evolved faster [37,38]). To understand whether this phenomenon occurs among *ATG* genes, we compared the sequence divergence levels from the representatives of lineages that diverged before duplication (henceforth, pre-duplication species; shaded in purple in Figure 1) between paralogs. While the cyclostomes share one round of whole-genome duplication with other vertebrates, only one gene survived in most cases. Therefore, cyclostomes were still considered pre-duplication species in our analysis, unless duplicate genes were found (also reflected in the gene expression analysis below; see Materials and Methods for details). Loss of the *ATG* genes also occurred in Gnathostomata (e.g., only two paralogs survived after two rounds of wholegenome duplications), which likely occurred independently from the losses inferred in cyclostomes [31,32].

The non-synonymous substitution rate (number of non-synonymous substitutions per non-synonymous site; dN) was calculated between each preand post-duplication sequence pairs (Figure 2A). The overall distribution of dN values varied among genes (e.g., low in *BECN1* [beclin 1], *WIPI2* [WD repeat domain, phosphoinositide interacting 2], *GABARAP, LC3A*, and *LC3B* [microtubule associated protein 1 light chain 3 alpha and beta] and high in *ATG16L2*), a reflection of variability in both mutation rates and tolerance of non-synonymous mutations among different genes [39] (Figure 2B). No obvious difference in dN was observed between paralogs in the *ULK-1* (*ULK1* and *ULK2*), *ATG2, LC3* (*LC3A* and *LC3B*), *ATG4-1* (*ATG4A* and *ATG4B*), and *ATG4-2* (*ATG4C* and *ATG4D*) pairs. In contrast, clear differences were observed in the *BECN, WIPI* (*WIPI1* [WD repeat domain, phosphoinositide interacting 1] and *WIPI2*), *GABARAP* (*GABARAP* and *GABARAPL1*), and *ATG16* pairs (Cliff’s δ [40], a measure of the magnitude of differences ranging between 0 and 1; a value over 0.47 is generally interpreted as “large” [41]; Figure 2C), indicating strong asymmetry in sequence evolution between paralogs. Consistent with the long branch separating mammalian from nonmammalian *ATG9B* orthologs in the phylogenetic tree (Figure S1B), *ATG9B* had significantly higher dN values compared with *ATG9A* in mammals, but not when all species are considered (enlarged inset in Figure 2B).

**Figure 2.**
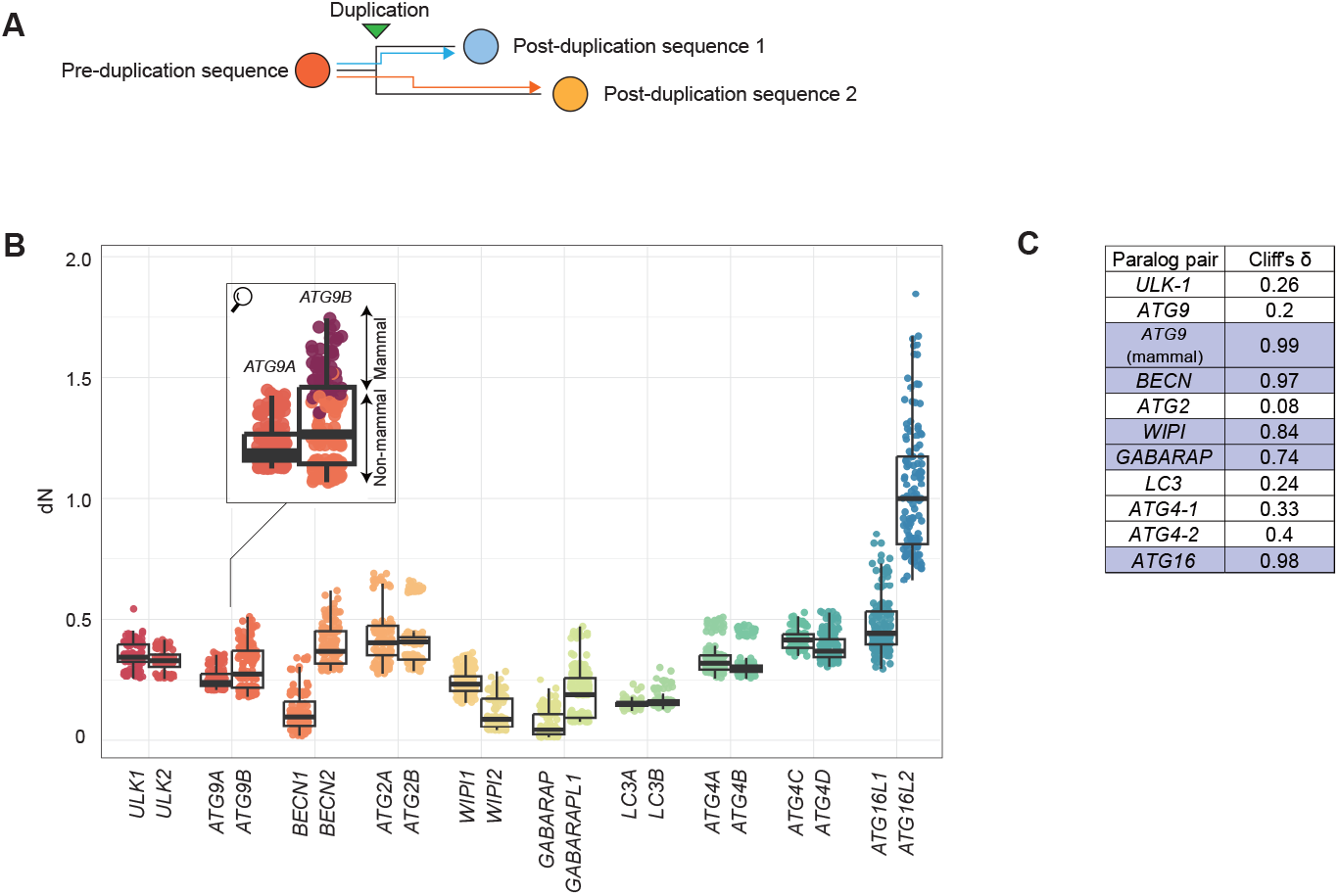
Paralogs within the *ATG9* (mammals only), *BECN, WIPI, GABARAP*, and *ATG16* pairs diverged highly asymmetrically at the sequence level. (**A**) Protein sequence divergence was quantified by dN, calculated for each pair of pre- and post-duplication sequences. (**B**) The dN values organized by paralog pairs. For display purposes, the *y*-axis is truncated at 2, and consequently, seven outliers are not displayed. The enlarged inset shows that sequence evolution of *ATG9B* differed between mammals (burgundy) and non-mammalian vertebrates (orange). (**C**) Cliff’s δ was used to compare the differences in dN between paralogs. Following convention, a value over 0.47 was interpreted as indicating a “large” difference (purple). A large difference in dN values was observed between *ATG9A* and *ATG9B* in mammals, but not when all species are considered.

Thus, paralogs in the *BECN, WIPI, GABARAP*, and *ATG16* pairs diverged highly asymmetrically from sequences of the pre-duplication species. *ATG9B* was comparable to *ATG9A* in non-mammalian vertebrates, but rapidly diverged in mammals.

### Paralogs within the BECN, WIPI, ATG4-1, and ATG16 pairs and ATG9 in mammals diverged asymmetrically in gene expression

We next examined the divergence in gene expression between paralogs. This analysis included 14 out of the 30 species shown in Figure 1; species were selected based on the availability of RNA-seq data from major tissues (brain, cerebellum, heart, intestine, kidney, liver, ovary, placenta, testis; Table S3). Among them, *Branchiostoma lanceolatum* and *Eptatretus burgeri* were considered pre-duplication species (Figure 1). The normalization procedure is described in the Materials and Methods section and Figure S2.

We calculated the gene expression and tissue specificity levels of each *ATG* gene in preand post-duplication species (Figure 3A, bottom and top panels). Tissue specificity was quantified by tau [42], which indicates how “even” expression levels across tissues are, with a lower value indicating broader expression. In general, tau is negatively correlated with gene expression level [14], which we also observed in our data. For example, *ULK4, LC3C, ATG9B*, and *BECN2* have low median expression levels across tissues and high tissue specificities.

**Figure 3.**
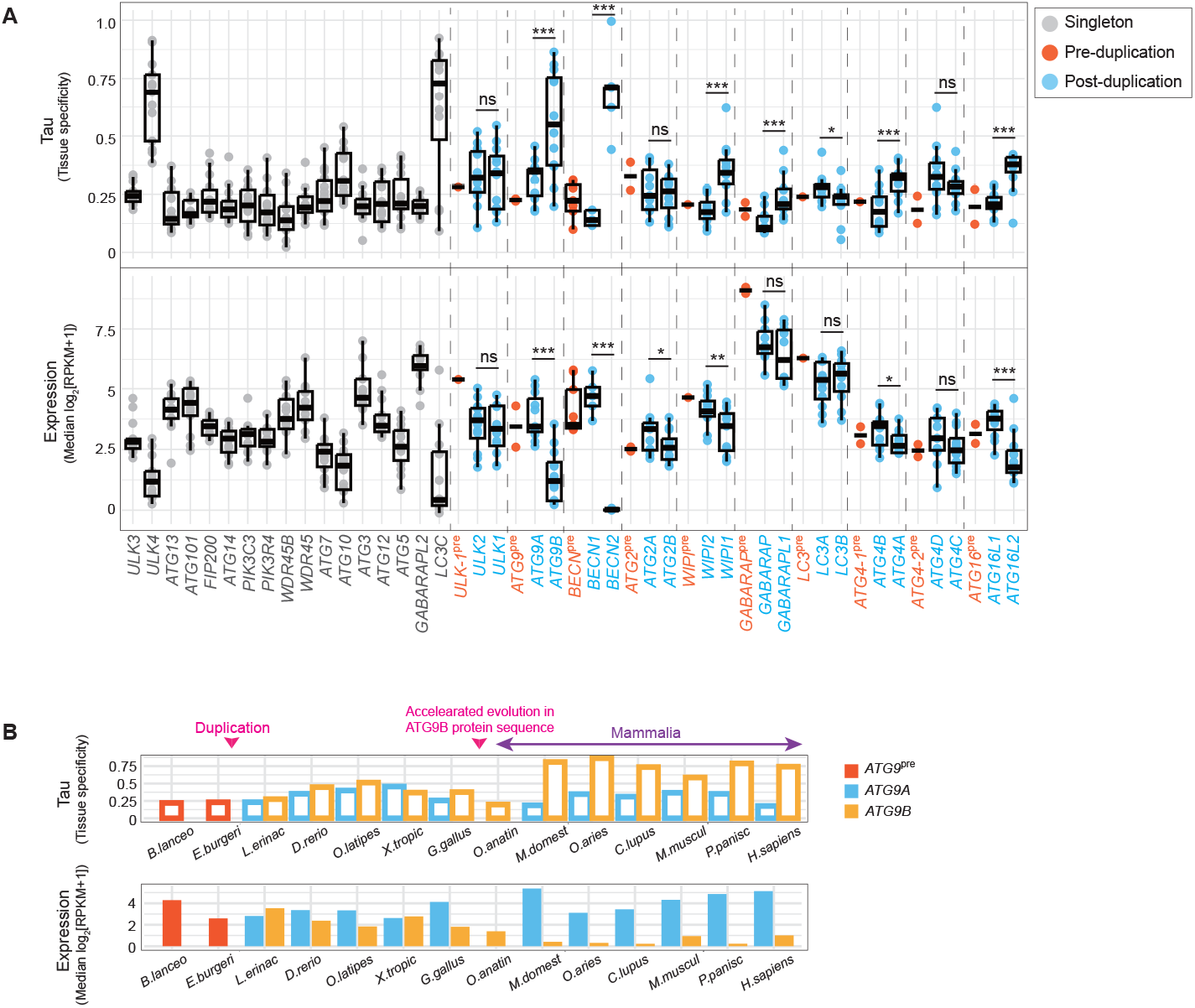
Paralogs within the *ATG9* (mammals only), *BECN, WIPI, ATG4-1*, and *ATG16* pairs diverged highly asymmetrically at the gene expression and tissue specificity levels. (**A**) The boxplot shows tissue specificity measured by tau (top) and the median gene expression levels across tissues (bottom); tau is a measure of the relative expression level in each tissue compared with the maximum, and a higher value indicated more tissue-specific expression. A generalized linear mixed model was used to compare the differences between paralogs. *, *p* < 0.05. **, *p* < 0.01. ***, *p* < 0.001. ns, non-significant (*p* > 0.05). (**B**) The tissue specificity and expression levels of *ATG9A* and *ATG9B* by species. The duplication timing and timing of accelerated protein evolution in *ATG9B* are labeled with pink arrows. *B. lanceo, Branchiostoma lanceolatum*; *E. burgeri, Eptatretus burgeri*; *L. erinac, Leucoraja erinacea*; *D. rerio, Danio rerio*; *O. latipes, Oryzias latipes*; *X. tropic, Xenopus tropicalis*; *G. gallus, Gallus gallus*; *O. anatin, Ornithorhynchus anatinus*; *M. domest, Monodelphis domestica*; *O. aries, Ovis aries*; *C. lupus, Canis lupus familiaris*; *M. muscul, Mus musculus*; *P. panisc, Pan paniscus*; *H. sapiens, Homo sapiens*.

Either no significant difference, or only a partial difference in gene expression and tissue specificity was detected between paralogs within the *ULK-1, ATG2, GABARAP, LC3*, and *ATG4-2* pairs; however, significant differences were observed within the *BECN, WIPI, ATG4-1*, and *ATG16* pairs (Figure 3A). Paralogs within the *BECN, WIPI*, and *ATG16* pairs also had the most asymmetric protein divergence (large Cliff’s δ value in Figure 2C), such that the paralog that diverged less from the pre-duplication species in terms of sequence (*BECN1, WIPI2*, and *ATG16L1*) also had higher and broader expression, suggesting a positive association between protein and gene expression evolution [14,43].

Mirroring its sequence divergence pattern, *ATG9B* had expression levels comparable to those of *ATG9A* in non-mammalian vertebrates, but reduced expression and elevated tissue specificity in mammals or placental mammals (Figure 3B; only *ATG9B* was found in the platypus *Ornithorhynchus anatinus*, the only non-placental mammal included in our analysis, and it had mostly ubiquitous, albeit lower, expression). Because mammals were enriched among the 14 species included in the RNA-seq analysis, the overall divergence pattern of *ATG9* was dominated by the mammalian pattern (Figure 3A). *ATG9B* is known to have placentaand pituitary gland-enriched expression in humans [44]. Across a broader range of mammals, *ATG9B* had either placenta- or testis-enriched expression (Figure S3).

Some other differences between mammals and non-mammalian vertebrates (not accompanied by obvious sequence-level change) were also observed (e.g., increased expression of *ULK1, ATG2B, GABARAPL1*, and *ATG4D* in mammals relative to their paralogs). Here, because the total number of species is limited, we did not conduct statistical tests for the subgroups of species. In future studies, changes between subgroups should be investigated among more species, samples, and tissue types.

Collectively, these findings indicate that paralogs in the *BECN, WIPI, ATG4-1*, and *ATG16* pairs differed significantly in both gene expression levels and tissue specificity. Similar to its sequence divergence pattern, *ATG9B* had reduced gene expression and elevated tissue specificity in mammals.

### ATG9B, BECN2, WIPI1, *and* ATG16L2 *evolved under relaxed negative selection in mammals*

Because paralogs within the *BECN, WIPI*, and *ATG16* pairs and *ATG9* pairs in mammals diverged highly asymmetrically at both the sequence and gene expression levels, we wondered whether they have also been under natural selection of differing strengths (i.e., negative selection, which purges deleterious mutations from the population, may be stronger in one paralog). To test this hypothesis, we calculated the ratio of the rates of non-synonymous to synonymous substitutions (the dN/dS ratio), a molecular signature of natural selection, of these paralogs in mammals. Assuming that only the non-synonymous mutations can affect function, the dN/dS ratio compares the rate at which non-synonymous substitutions are permitted relative to a mutation rate baseline (i.e., the synonymous mutation rate). When dN/dS < 1, lower values indicate stronger negative selection (deleterious mutations are removed more quickly from the population).

The dN/dS ratio was calculated using Phylogenetic Analysis by Maximum Likelihood (PAML) [45]. Given a phylogenetic tree, PAML calculates the dN/dS ratio along each branch by comparing the extant sequences with the inferred ancestral sequences. For our purpose, we calculated one dN/dS ratio per gene. Gene expression level is known to be negatively correlated with the dN/dS ratio (i.e., highly expressed genes have lower dN/dS ratios) [46], which we also observed among the *ATG* genes, including *BECN2, ATG9B*, and *ATG16L2* (Figure 4A). This negative correlation is hypothesized to occur because a mutation (especially a non-synonymous mutation) in a highly expressed gene tends to incur a greater cost (translational cost, risk of misinteraction given its higher concentration, or a cost incurred by its reduced physiological function) [46].

**Figure 4.**
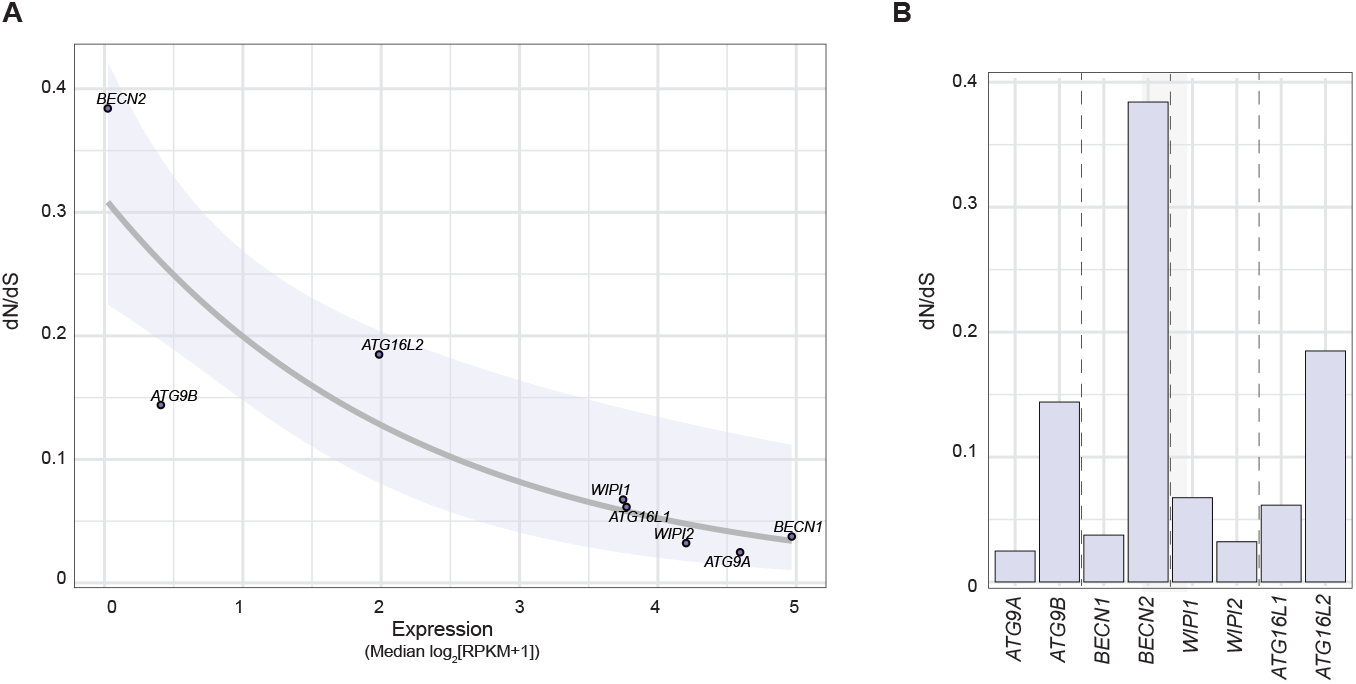
Within the *ATG9, BECN, WIPI*, and *ATG16* pairs, *ATG9B, BECN2, WIPI1* and *ATG16L2* were under more relaxed negative selection in mammals. (**A**) The dN/dS ratio (*y*-axis) is negatively correlated with the median gene expression level across tissues (*x*-axis). The best-fit line (generalized linear model with log link) and its 95% confidence interval (shaded region) are shown. (**B**) The dN/dS ratio of each *ATG* gene in the group of around 120 mammals was calculated using PAML. Lower dN/dS ratios indicate stronger negative selection.

As expected, *ATG9B, BECN2, WIPI1*, and *ATG16L2* all had substantially higher dN/dS ratios than their paralogs, suggesting that deleterious mutations have been less efficiently removed from these genes (i.e., they are under more relaxed negative selection) in mammals (Figure 4B). Because strong negative selection is an indication of conserved function [47,48], the ancestral function may have been better preserved in the other paralog.

### Evolutionary fate classification based on RNA-seq data

To connect the evolutionary divergence patterns to functional differences, we categorized the evolutionary fates, i.e., how much each paralog contributes to the ancestral function(s), of the *ATG* paralogs. Common evolutionary fate categories include ancestral function being preserved by one paralog (allowing the other paralog to evolve under less constraint), subfunctionalization, or hypofunctionalization/dosage sharing [2-5]. Assuming that a similar expression pattern implies the conservation of function, we categorized the *ATG* genes based on whether a single paralog or the total expression of two paralogs had the expression pattern most similar to that found in the pre-duplication species as a proxy for the pre-duplication pattern [49,50] (Figure 5Ai).

**Figure 5.**
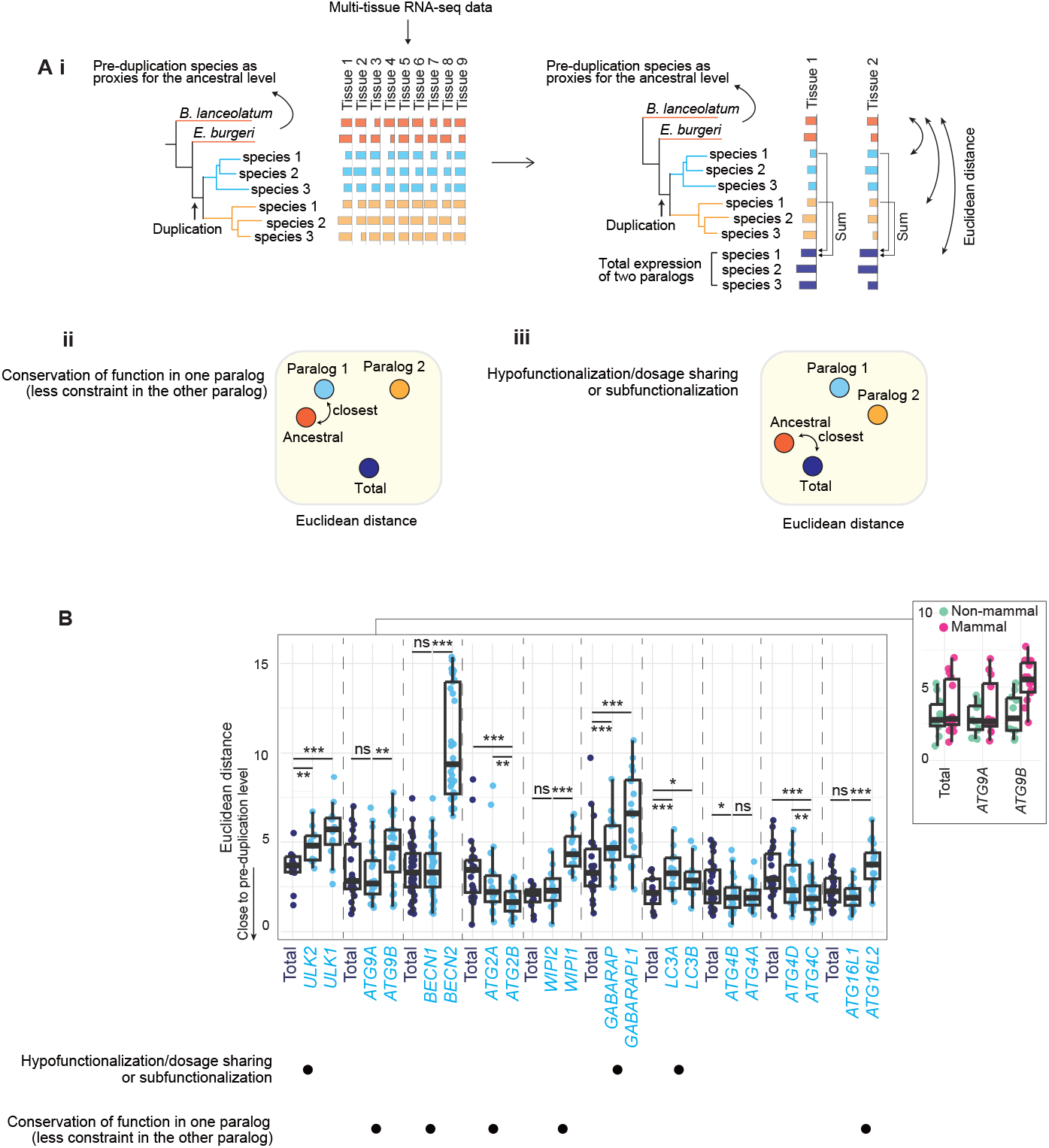
Evolutionary fate classification of the *ATG* paralogs according to their absolute gene expression levels (log_2_[RPKM+1]). (**A**) **i**, The evolutionary fate was classified by comparing the multi-tissue gene expression of either individual paralogs or the total expression of two paralogs to that of the preduplication level. Differences in gene expression were quantified by Euclidean distance. **ii**, If one of the paralogs had the lowest Euclidean distance from the pre-duplication level (i.e., was most similar), this paralog was inferred to likely preserve the ancestral function, allowing the other paralog to evolve under less constraint. **iii**, If the total expression of two paralogs had the lowest Euclidean distance from the pre-duplication level, the pair was classified as hypofunctionalization/dosage sharing or subfunctionalization. (**B**) Euclidean distances calculated from the absolute gene expression levels (log_2_[RPKM+1]), with associated evolutionary fate categories indicated below. A panel showing the differences between mammals and non-mammalian vertebrates within the *ATG9* pair is shown on the right. A GLMM was used to compare the differences in the Euclidean distances. *, *p* < 0.05. **, *p* < 0.01. ***, *p* < 0.001. ns, non-significant (*p* > 0.05).

Similarity to the pre-duplication expression level was measured as the Euclidean distance, calculated as Euclidean distance 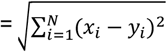, where *N* is the number of tissues and *x*_*i*_ and *y*_*i*_ are the expression levels of the pre- and post-duplication copy (or the total expression of the post-duplication copies). To better differentiate between different evolutionary fate categories (see below), we calculated the Euclidean distances using both the absolute and relative gene expression levels (ratios normalized to the total expression summed across all tissues). For simplicity, these are referred to as the absolute and relative Euclidean distances.

The classification rule that was applied is as follows: if one paralog has the lowest Euclidean distance (i.e., the most similar expression pattern) from the pre-duplication level, this paralog likely preserved the ancestral function, allowing the other paralog to evolve under less constraint (Figure 5Aii). Sometimes this other paralog would have been able to develop new functions, as predicted by Ohno’s neofunctionalization model [1]. If the total expression of two paralogs had the lowest *absolute* Euclidean distance from the pre-duplication level, this paralog pair was deemed consistent with either subfunctionalization or hypofunctionalization/dosage sharing (Figure 5Aiii), and the *relative* Euclidean distance was then employed to differentiate between these cases. At the gene expression level, complementary relative expression across tissues (i.e., one paralog is dominant in tissue A, while the other paralog is dominant in tissue B) is often interpreted as evidence of subfunctionalization. In hypofunctionalization or dosage sharing (also known as quantitative subfunctionalization), the absolute expression level is reduced in each paralog post-duplication, but there can be no change in relative expression among tissues [51-53]. Therefore, in this case, the paralog pair can be categorized as demonstrating subfunctionalization if the total expression has the lowest *relative* Euclidean distance and can be categorized as hypofunctionalization otherwise.

Overall, depending on the expression level of the pre-duplication species and the level of asymmetry between paralogs, two types of patterns emerged. Within the *BECN, WIPI*, and *ATG16* pairs, consistent with the strong asymmetry observed, one paralog (referred to as the major paralog, i.e., *BECN1, WIPI2*, and *ATG16L1*, respectively) or the total expression had the lowest Euclidean distance from the preduplication level (i.e., no significant difference between the two), while the other (minor) paralog had a significantly higher Euclidean distance (absolute and relative Euclidean distances are shown in Figure 5B and Figure S4, respectively), suggesting that the major paralog alone is likely sufficient to recapitulate the pre-duplication expression level, which is consistent with the ancestral function being preserved by the major paralog. The *ATG9* pair also had a similar pattern, but in this case, *ATG9A* became the major paralog only in mammals (and the overall pattern was driven by an enrichment of mammals in the RNA-seq data; Figure 3B and Figure 5B [right panel]). Within the *ATG2* pair, *ATG2B* had the lowest Euclidean distance from the pre-duplication level, suggesting that the ancestral function might have been preserved in *ATG2B*.

Within the *ULK-1, GABARAP*, and *LC3* pairs, the total expression of two paralogs exhibited the lowest absolute Euclidean distance from the pre-duplication level (Figure 5B), suggesting either subfunctionalization or hypofunctionalization. However, because there was no statistically significant difference between the relative Euclidean distances (Figure S4), hypofunctionalization/dosage sharing is the more likely fate among these three paralog pairs. We note that, for the *ULK-1* pair, there exists some difference between mammals and non-mammalian vertebrates (*ULK2* has a lower relative Euclidean distance in mammals), but the total expression always exhibited the lowest absolute Euclidean distance.

The *ATG4-1* pair could not be categorized according to these criteria because there was no statistically significant difference in either absolute or relative Euclidean distances between *ATG4A* and *ATG4B* (Figure 5B and Figure S4). The *ATG4-2* pair could not be categorized because of the differences both between results based on the absolute and relative Euclidean distances and between mammals and non-mammalian vertebrates (*ATG4D* has higher expression and increased relative Euclidean distance in mammals).

Overall, the *BECN, WIPI, ATG16*, and *ATG2* pairs were most consistent with the ancestral function being preserved in the major paralog, while the *ULK-1, GABARAP*, and *LC3* pairs were most consistent with hypofunctionalization/dosage sharing, in which expression of both paralogs is needed for function. For the *ATG9* pair, *ATG9B* showed a change in its expression patterns, and *ATG9A* is likely the paralog that has preserved the ancestral function in mammals.

### The relationship between the evolutionary fate category and autophagic function

*ATG* genes can have both autophagic and non-autophagic functions [6]. Because the autophagy pathway is an ancient pathway broadly conserved among eukaryotes, it is likely the major, if not the only, ancestral function of the *ATG* genes. Accordingly, it is expected that among pairs in which one paralog likely has preserved the ancestral function, this paralog would be most important for autophagy, while the other paralog may be dispensable; in contrast, among pairs classified as exhibiting hypofunctionalization or dosage sharing, both paralogs would be important for autophagy.

Indeed, among the *GABARAP* and *LC3* genes, *LC3A, LC3B, GABARAP*, and *GABARAPL1* can all rescue autophagy (mitophagy) activity in mammalian cells, although *LC3B* is sometimes found to have reduced activity [19,54], consistent with the hypofunctionalization/dosage sharing classification (although dosage constraints cannot be concluded from singlegene rescue experiments).

Within the *WIPI* and *ATG16* pairs, *WIPI2* is nearly essential for autophagy while *WIPI1* alone cannot rescue autophagy flux [26]; additionally, *ATG16L1* is required for autophagy, while *ATG16L2* is not [55,56]. These findings are consistent with the evolutionary fate category in which the major paralog has preserved the ancestral function.

Within the *ATG9* pair, evolutionary fate classification suggests that *ATG9A* likely preserves the ancestral function in mammals. However, since exogenous expression of *ATG9B* can still rescue autophagic flux in HEK293A cells [25], the protein sequence of *ATG9B* clearly has not diverged enough to abolish its scramblase function. One possible explanation for this finding may be that *ATG9B* divergence occurred more recently compared with, for example, the divergence of *WIPI1* and *ATG16L2*, which can no longer rescue autophagy in mammalian cells.

For the *ATG2* pair, even though *ATG2B* is the one preserving the ancestral function according to our classification, both *ATG2A* and *ATG2B* have redundant functions in mammalian cells [57,58], suggesting that the pre-duplication gene expression level may not be an adequate measure of function in this case.

Although *ULK1* and *ULK2* are generally considered redundant, some tissueand celltype-specific differences have been reported [24,59]. Between *BECN1* and *BECN2*, although *BECN1* is likely the major paralog, some studies suggest that *BECN2* may also be able to function in autophagy [27]. To further clarify the functions of the *ULK-1* pair and *BECN2* in autophagy, their ability to rescue autophagy flux was tested using the HaloTag-based processing assay [60]. Autophagy delivers HaloTag-SNAP to the lysosome, where HaloTag is efficiently degraded. However, HaloTag is stabilized upon ligand binding, enabling the accumulation of free Halo bands, the amount of which reflects autophagic flux. HaloTag-SNAP processing was accelerated in wildtype (WT) mouse embryonic fibroblasts (MEFs) after starvation but not in *RB1CC1*/*FIP200* (RB1 inducible coiled-coil 1) knockout (KO) MEFs or in *ULK1/2* double knockout (DKO) MEFs. When green fluorescent protein (GFP)-ULK1 or GFPULK2 was expressed in *ULK1/2* DKO MEFs, either could individually restore HaloTag-SNAP processing, although to a degree somewhat lower than that observed in WT cells (Figure 6A, B), suggesting that both *ULK1* and *ULK2* can rescue autophagic flux. In HeLa cells, deletion of only *BECN1* was sufficient to block autophagy (Figure 6C, D). When GFP-Beclin 1 or GFP-Beclin 2 was expressed in *BECN1* KO HeLa cells, GFP-Beclin 1 robustly restored HaloTag-SNAP processing, while GFP-Beclin 2 did not (Figure 6C, D). Furthermore, HaloTag-SNAP processing was normal in *BECN2* KO cells, suggesting that only *BECN1* is essential for autophagy. These experimental results are consistent with their respective evolutionary fate categories (i.e., hypofunctionalization/dosage sharing for *ULK-1* and ancestral function being preserved in the major paralog for *BECN*).

**Figure 6.**
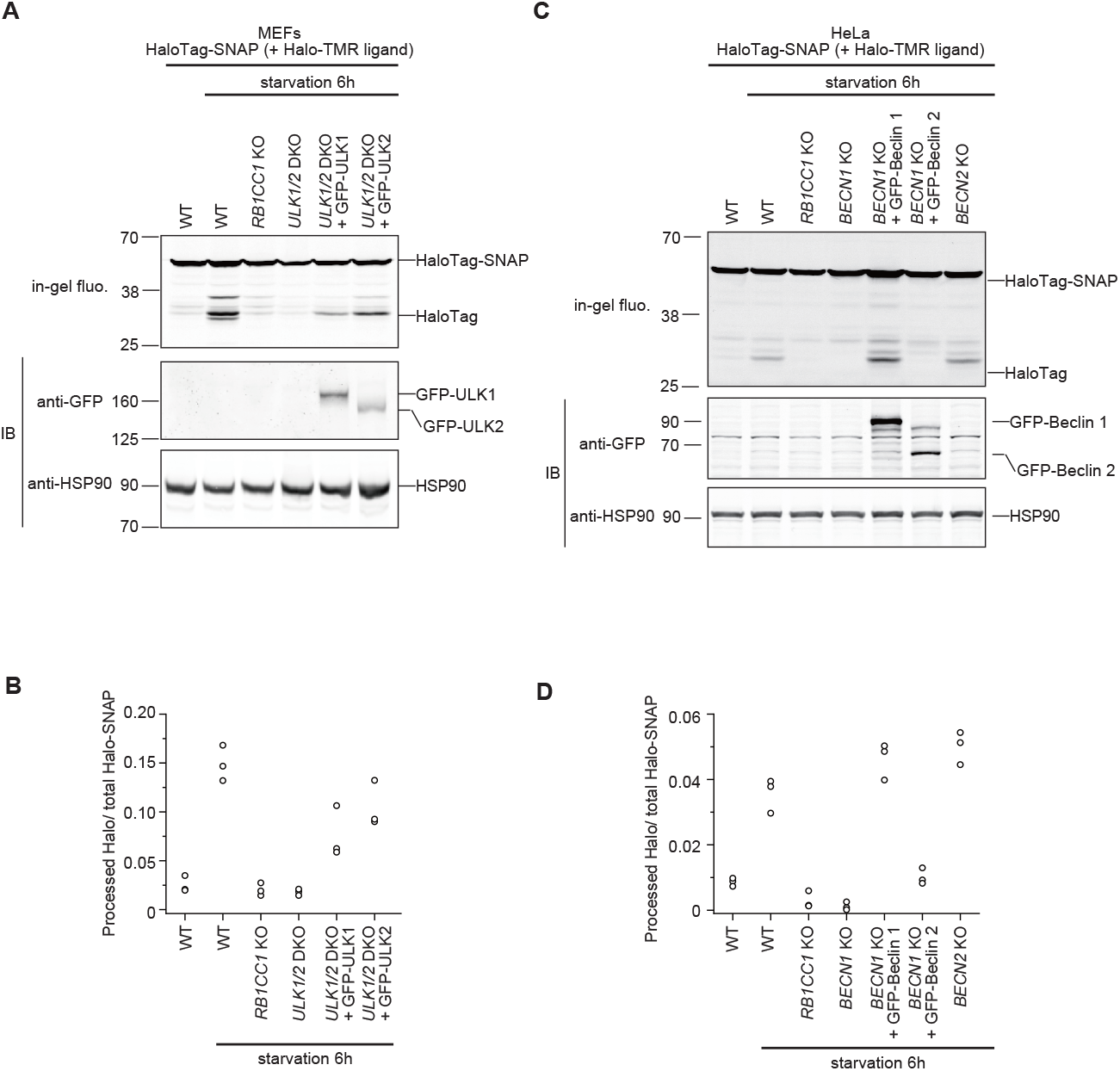
Analysis of autophagic flux in the *ULK-1* and *BECN* paralogs. (**A and C**) HaloTag-SNAP processing assay. In this assay, HaloTag-SNAP is incorporated into autophagosomes non-selectively. After TMR-conjugated HaloTag ligand (Halo-TMR ligand) binding, HaloTag protein becomes stable and accumulates in autolysosomes, the amount of which reflects autophagic flux. Cells stably expressing HaloTag-SNAP were labeled for 30 min with 100 nM Halo-TMR ligand and then were incubated in starvation medium for 6 h. Total cell lysates were subjected to immunoblotting with the indicated antibodies or in-gel fluorescence detection. (**B and D**) HaloTag-SNAP processing assays across three independent experiments. The HaloTag-SNAP processing rate was calculated as the band intensity of processed HaloTag over the sum of the band intensities of the processed HaloTag and unprocessed HaloTag-SNAP.

## Discussion

Gene duplication is a key driver of evolutionary diversification and novelty. Here, with a focus on the *ATG* gene duplications in vertebrates, we present an analysis of the evolutionary diversification patterns between paralogs and a discussion of their functional implications in light of previous experimental studies in mammals. We also provide new experimental data comparing the ability of both *ULK1* and *ULK2* and both *BECN1* and *BECN2* to rescue autophagic flux using the HaloTag-SNAP assays.

Overall, following gene duplication, ancestral gene expression may have been partitioned between two paralogs in the *ULK-1, GABARAP*, and *LC3* pairs, such that both paralogs are important for autophagy, and only weak or partial asymmetry in expression and sequence evolution exists between paralogs (Figure 7). In comparison, strong asymmetry has been observed between the paralogs in the *BECN, WIPI*, and *ATG16* pairs, and only one paralog (*BECN1, WIPI2*, and *ATG16L1*) is required for autophagic function (the other paralog lost autophagic function at least in mammals, but possibly earlier, considering the asymmetry). *ATG9B*, while comparable to *ATG9A* in non-mammalian vertebrates, diverged in both sequence and expression level in mammals. *ATG9B* can still function if exogenously expressed [25], but is less likely to play a major role given its low expression level in most tissues examined here (Figure 3B).

**Figure 7.**
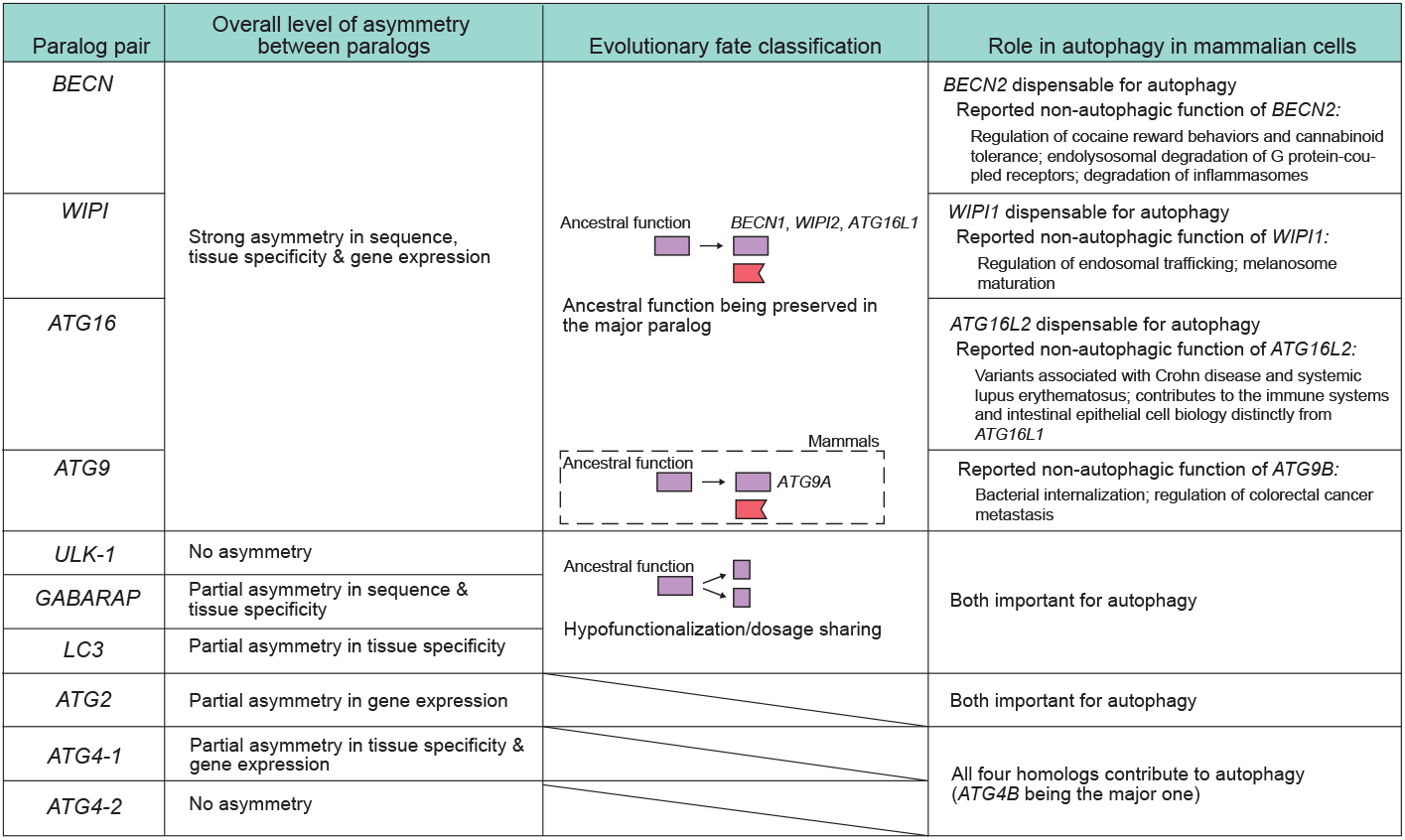
A summary of the level of asymmetry, evolutionary fate classification, and functions in mammalian cells (based on previous reports and experimental data from this study). The *ULK-1, GABARAP*, and *LC3* pairs exhibit weak or partial asymmetry between paralogs; among these pairs, both paralogs contribute to the overall gene expression level (hypofunctionalization/dosage sharing), and both are important for autophagy, which is likely a key ancestral function of the *ATG* genes. In contrast, the *BECN, WIPI*, and *ATG16* pairs display strong asymmetry between paralogs, such that *BECN1, WIPI2*, and *ATG16L1* likely preserve the ancestral function, allowing the other paralog to evolve under less constraint. *BECN2, WIPI1*, and *ATG16L2* are dispensable for autophagy, and the reported non-autophagic functions are listed (see text for references). *ATG9B* diverged from *ATG9A* in terms of both sequence and gene expression in mammals. The *ATG2, ATG4-1*, and *ATG4-2* pairs exhibit weak or partial asymmetry. Both *ATG2A* and *ATG2B* and all of the *ATG4* genes contribute to autophagy.

One exception may be the placenta (Figure S3), although knocking in a C-terminal-truncated *ATG9B* in mice resulted in no obvious phenotype, including in the placenta [61]. The *ATG2, ATG4-1*, and *ATG4-2* pairs exhibited weak or partial asymmetry in expression and sequence evolution, but their evolutionary fates could not be classified or were inconsistent with existing studies.

The evolutionary divergence of *BECN2, WIPI1*, and *ATG16L2* from their paralogs and their loss of autophagic function raise the question of why these genes were not pseudogenized and lost. While it has been suggested that within an asymmetric paralog pair, the less-expressed paralog can slowly approach pseudogenization [62], at least some non-autophagic functions have been reported for these genes, including the regulation of cocaine reward behaviors and cannabinoid tolerance [63,64], endolysosomal degradation of G protein coupled-receptors [27], and degradation of inflammasomes for *BECN2* [65,66], as well as the regulation of endosomal trafficking and melanosome maturation for *WIPI1* [67,68]. Variants in *ATG16L2* are associated with Crohn disease and systemic lupus erythematosus [69-71], and this gene contributes to the immune system and intestinal epithelial cell biology distinctly from *ATG16L1* [56]. *ATG9B*, which diverged from *ATG9A* in mammals, is reported to regulate bacteria internalization and colorectal cancer metastasis [72,73]. Many of these reported functions are related to the central nervous system or the immune system, which are later additions in evolution (compared with the autophagic function), suggesting the acquisition of new functions may be a mechanism contributing to the preservation of these genes.

One limitation of this study is the limited number of species in the RNA-seq analysis; specifically, a limited number of pre-duplication species prevented the reconstruction of the ancestral expression (as opposed to using the expression levels of the pre-duplication species as proxies), and a limited number of post-duplication species complicates the examination of subgroup-specific changes. In the future, more species, samples, tissues, and cell types should be used for such analyses (should they become available).

## Materials and Methods

### Selection of species

For the homology search and protein sequence evolution analysis, 30 species in Chordata were selected based on their evolutionary positions and the quality of their genome assemblies, and five outgroup species were selected either because they are commonly used model organisms (*Drosophila melanogaster* and *Caenorhabditis elegans*), evolutionarily close to Chordata (*Strongylocentrotus purpuratus*), or commonly used as representative invertebrate species in vertebrate genome evolution studies (*Mizuhopecten yessoensis* and *Trichoplax adhaerens* [30,31]). Of these 30 species, 14 with publicly available multi-tissue RNA-seq data were included in the gene expression evolution analysis. The species and RNA-seq sample information are summarized in Table S1 and Table S3, respectively. The time-calibrated species tree in Figure 1 was downloaded from TimeTree [74], and the silhouette images, all in the public domain, were obtained from PhyloPic (https://www.phylopic.org).

### Homology search and determination of duplication timing

The genome, gene annotation, and protein fasta files were downloaded from either the National Center for Biotechnology Information Assembly or Ensembl databases (Table S1). Protein Basic Local Alignment Search Tool (BLASTP) search (v2.13.0+) was conducted using the human sequences as queries and the downloaded protein fasta files (based on only the longest sequence per gene according to the gene annotation) as the database [75], and domain enhanced lookup time accelerated BLAST search results were checked when no homologs were found by BLASTP [76]. For *ULK1–4, PIK3C3* (phosphatidylinositol 3-kinase catalytic subunit type 3), *PIK3R4* (phosphoinositide-3-kinase regulatory subunit 4), and *ATG16L1–2*, because BLASTP search also identified non-homologs with similar domains, conserved domain search and reciprocal BLASTP (only reciprocal BLASTP for *PIK3C3*) were used to filter out non-homologs [77]. The candidate sequences were further checked and annotated by reconstructing the phylogenetic trees (some short sequences were removed, and some incomplete sequences were counted as a single sequence). For the phylogenetic analysis, protein sequences were aligned using MUSCLE (v3.8.1551) [78], and columns with >50% gaps were removed using trimAl (v1.4.rev15) [79]. Phylogenetic trees were reconstructed using IQ-TREE (v2.1.4) with ModelFinder and options “-alrt 1000 -B 1000 -T 4 -nmax 5000 --bnni” [80], and then visualized using the ggtree package (v3.8.2) [81].

### Calculation of protein sequence divergence

The pairwise dN values between sequences of the preand post-duplication species were calculated using PAML [82]. Sequences deemed too short (less than approximately half the length of the human protein) were removed from the analysis. While cyclostomes were treated as pre-duplication species because most *ATG* genes exist as singletons in this clade, the following sequences were removed because duplicate genes were found: *ULK-1* from *Eptatretus burgeri, WIPI* from *Eptatretus burgeri* and *Eptatretus atami, LC3* from all four cyclostomes, and *ATG4-2* from *Petromyzon marinus* and *Lethenteron reissneri* (and similar removals were made for the RNA-seq data described below). *Taeniopygia guttata* was removed from the *ATG2* analysis because its *ATG2A* sequence (XP_041567864.1) is on a long branch in the phylogenetic tree (Figure S1D). Sequences corresponding to other noticeable long branches in the post-duplication species were either already excluded owing to their short length or confirmed to have no effect on the statistical significance, whether excluded or included. Cliff’s δ was used to compare the distribution of dN values between paralogs [40,41].

### Quality control and normalization of the RNAseq data

The overall quality control and normalization pipeline is summarized in Figure S2A. Briefly, the RNA-seq samples were downloaded (Table S3), trimmed using fastp with the option “-5 -3” (v0.23.2) [83], and mapped to their respective genomes using STAR with the options “-runThreadN 4 --twopassMode Basic --outSAMstrandField intronMotif --readFilesCommand zcat --outSAMtype BAM SortedByCoordinate --outSAMunmapped Within” (v2.7.10b) [84]. Then, reads mapped to each gene were counted using featureCounts (v2.0.6) [85]. Results counting only the uniquely mapped reads (default) versus counting both uniquely mapped and multi-mapping reads (as fractions using “-M --fraction”) were compared, and genes for which read counts differed by more than 20% were removed from further analysis.

Three steps were applied in the normalization pipeline (roughly a combination of methods used in two previous publications [14,15], using code obtained from one of them [15]). First, single-copy orthologs (orthologs with exactly one copy in each species) were identified using OrthoFinder (v2.5.4) [86], and the trimmed mean of M values (TMM) normalization was applied based on the read counts of the single-copy orthologs [87]. Here, single-copy orthologs were used, as gene duplication is expected to have an effect on gene expression levels. Second, the expression levels were calculated as log_2_(reads per kilobase per million mapped reads [RPKM]+1), and surrogate variable analysis (SVA) was applied to remove noise in gene expression caused by factors such as technical artefacts [88,89]. Here, SVA was conducted for each species separately instead of all species together, because in the latter case, genes not present in all species would have to be excluded. Tissue of origin was used as the primary variable, and as expected, SVA increased the Pearson’s *r* coefficient (calculated using all available genes) between samples from the same tissue (Figure S2C). SVA was not performed for two species in which there was only one sample per tissue in the dataset (*Danio rerio* and *Oryzias latipes*) and for another two species in which SVA resulted in numerical errors (*Eptatretus burgeri* and *Pan paniscus*; the correlation between samples from the same tissues was generally high for these species). Third, to remove any potential (if any) global differences in gene expression among species, a group of genes with relatively stable expression across species and samples (*N* = 375) was identified, and the median expression levels of these genes were used to calculate a normalizing factor (relative to sample SRR6246036; Figure S2B). The genes with relatively stable expression were determined from the single-copy orthologs as follows: the relative expression levels of these genes in each sample (percent rank) were calculated, and after removing genes with median percent ranks across samples within the top or bottom 25% of all genes, the remaining genes with the lowest standard deviations in percent ranks were chosen.

### Analysis of gene expression evolution

The tissue specificity measure that was used is tau [42], defined as 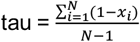, where *N* is the number of tissues and *x*_*i*_ is the expression level in tissue *i* relative to the maximum expression level among all tissues. Occasionally, the expression levels (log_2_[RPKM+1]) of very low-expression genes became negative after multiple rounds of normalizations. In such cases, 0 was used instead of a negative value when calculating tau (to avoid tau > 1). Using expression levels from the pre-duplication species (*Branchiostoma lanceolatum* and *Eptatretus burgeri* for most *ATG* genes and more species according to the inferred duplication timing for the *GABARAP* and *BECN* pair) as proxies of the ancestral expression levels, the Euclidean distances between the expression levels of the pre-duplication species and either individual post-duplication paralog or the total expression of both paralogs were calculated. Euclidean distances were calculated using both the absolute expression levels (log_2_[RPKM+1]; Figure 5B) and relative gene expression levels (absolute expression in each tissue divided by the total expression across all tissues; Figure S4). A generalized linear mixed model (GLMM) treating the phylogenetic relationships as random effects and *ATG* genes (and the total expression in the case of Euclidean distance) as fixed effects was used to compare the post-duplication divergence in tissue specificity and gene expression between paralogs (Figure 3A) as well as the Euclidean distances (Figure 5B and Figure S4). In the latter case, identity of the pre-duplication species was also included as a random effect variable. The GLMM was fitted using the phyr package in R (v1.1.0) [90], and the phylogenetic tree used in the GLMM was the time-calibrated species tree downloaded from TimeTree [74]. In all gene expression analyses, if more than one sequence was identified for a paralog within a post-duplication species (i.e., from secondary duplications or technical issues related to the genome assembly), the max expression level of all such sequences was used in the analyses (to avoid including odd sequences with low expression levels).

### Calculation of the per-gene dN/dS ratio

Gene annotations and pairwise alignments against the human sequences were downloaded from the Zoonomia project [91,92]. Sequences that contained at least one segment without a premature stop codon that would cover >50% of the human sequence and with the classification “I” or “PI” (intact or partially intact) were included.

The OMM-MACSE pipeline was used to obtain codon-level alignments and remove both falsepositive sequences and poorly aligned regions [93]. The sequences were further manually checked based on the phylogenetic analysis. Next, Treemmer was used to downsample the number of species to around 120 while still retaining the major mammalian lineages [94]. The species-level phylogenetic tree used for downsampling was downloaded from VertLife [95]. The number of sequences remaining differed slightly among *ATG* genes. The M0 model in PAML was used to calculate the dN/dS ratios for each gene (v4.10.6) [82].

### Cell lines and cell culture conditions

MEFs and HeLa cells were cultured in a 5% CO_2_ incubator at 37°C, using Dulbecco’s Modified Eagle Medium (Wako Pure Chemical Corp., 043-30085) supplemented with 10% fetal bovine serum (Sigma-Aldrich, 173012). *RB1CC1* KO and *ULK1/2* DKO MEFs were provided by Prof. JunLin Guan and Prof. Thompson [96,97], respectively. *RB1CC1* KO HeLa cells were previously generated [98]. *BECN1* and *BECN2* KO HeLa cells were established using the CRISPR-Cas9 system as previously described [26], with the following gRNAs: human *BECN1*, 5′-GCCTGGATGGTGACACGGTCC-3′, human *BECN2*, 5′GTCGGTGCATTCTTCACACAG-3′.

### Preparation of retrovirus for stable expression

DNA fragments encoding HaloTag7 (Promega, N2701), SNAP-tag (New England BioLabs, N9181S), human *BECN1* (NM_001313998.2), and *BECN2* (NM_001290693.1) were inserted into pMRX-IP retroviral plasmids [99,100]. Retroviral plasmids containing GFP-ULK1 and GFP-ULK2 were previously generated [101]. For the preparation of retrovirus, HEK293T cells were transfected with a retroviral vector together with pCG-VSV-G and pCG-gag-pol (gifts from Dr T. Yasui) using Lipofectamine 2000 (Thermo Fisher Scientific, 11668019). Two days after transfection, the supernatant was passed through a 0.45-μm syringe filter unit (Merck Millipore, SLHV033RB) and collected. Then, the retrovirus was applied to cells, and stable transformants were selected by application of 2 μg/ml puromycin (Sigma-Aldrich, P8833). If required, cells expressing multiple tagged proteins were sorted using CytoFLEX SRT (Beckman Coulter).

### Protein extraction and immunoblotting

Cells were lysed with 50 mm Tris–HCl, pH 7.5, 150 mm NaCl, 1 mm MgCl_2_, 1% Triton-X100, and protease inhibitor (Nacalai Tesque, 03969-34) on ice. The cell lysates were mixed with sample buffer and heated to 95°C for 5 min, after which they were subjected to sodium dodecyl sulfate polyacrylamide gel electrophoresis and then transferred to a polyvinylidene difluoride membrane (Merk Millipore, IPFL00010) using the Trans-Blot Turbo Transfer System (BioRad). Immunoblotting analysis was performed with the following antibodies: rabbit anti-GFP antibody (Invitrogen, A6455), mouse anti-HSP90 antibody (BD Transduction Lab, 610419,), donkey anti-rabbit IgG antibody-IRDye 800CW (LI-COR, 926-32216), and donkey anti-mouse IgG antibody-IRDye 680LT (LI-COR, 926-68072). The fluorescent signals were visualized with the Odyssey M imager (LI-COR). Contrast and brightness were adjusted and quantified using the ImageJ (v1.54f) -Fiji image processing package [102].

### HaloTag-SNAP processing assay

Cells were incubated with 100 nM tetramethylrhodamine (TMR)-conjugated HaloTag ligand (Promega, G8251) for 30 min. After being washed twice with phosphate-buffered saline, cells were cultured in starvation medium for 6 h. Then, cells were lysed, and proteins were obtained as described in the previous section. Proteins were separated by sodium dodecyl sulfate polyacrylamide gel electrophoresis, and the gel was visualized for in-gel fluorescence from TMR with the Odyssey M imager. The signal intensities of the bands of HaloTag-SNAP and free HaloTag were measured with ImageJ-Fiji. The HaloTag-SNAP processing rate was calculated as the intensity of the free HaloTag band divided by the sum of the intensities of the free HaloTag and unprocessed HaloTag-SNAP bands.

## Acknowledgments

We thank Toshio Kitamura and Shoji Yamaoka for providing pMRXIP, and Teruhito Yasui for providing pCG-VSV-G and pCG-gag-pol. We would like to express our sincere gratitude to Nozomi Sato for technical assistance, and to all of the Mizushima lab members for helpful discussions. This work was supported by the Exploratory Research for Advanced Technology (ER-ATO) research funding program of the Japan Science and Technology Agency (JST) (JPMJER1702 to N.M.), by a Grant-in-Aid for Specially Promoted Research (22H04919 to N.M.), and by the Chugai Foundation for Innovative Drug Discovery Science (C-FINDs) Postdoctoral Fellowship for Foreign Researchers (to S.Z.).

## Data availability statement

The sources of the genomic and RNA-seq data used in this study (all publicly available) are listed in Table S1 and Table S3. The programming code and intermediate result files associated with this study will be openly available in figshare.

## Abbreviations

ATG: autophagy-related
BLAST: Basic Local Alignment Search Tool
DKO: double knockout
GFP: green fluorescent protein
GLMM: generalized linear mixed model
KO: knockout
LC3: MAP1LC3
MEF: mouse embryonic fibroblast
ns: non-significant
PAML: Phylogenetic Analysis by Maximum Likelihood
RPKM: reads per kilobase per million mapped reads
SVA: surrogate variable analysis
TMM: trimmed mean of M values
TMR: tetramethylrhodamine
WT: wild type

