## Supplementary Figures and Tables for "*ATG* gene duplication in vertebrates: evolutionary divergence and its functional implications"

Figure S1

**A**

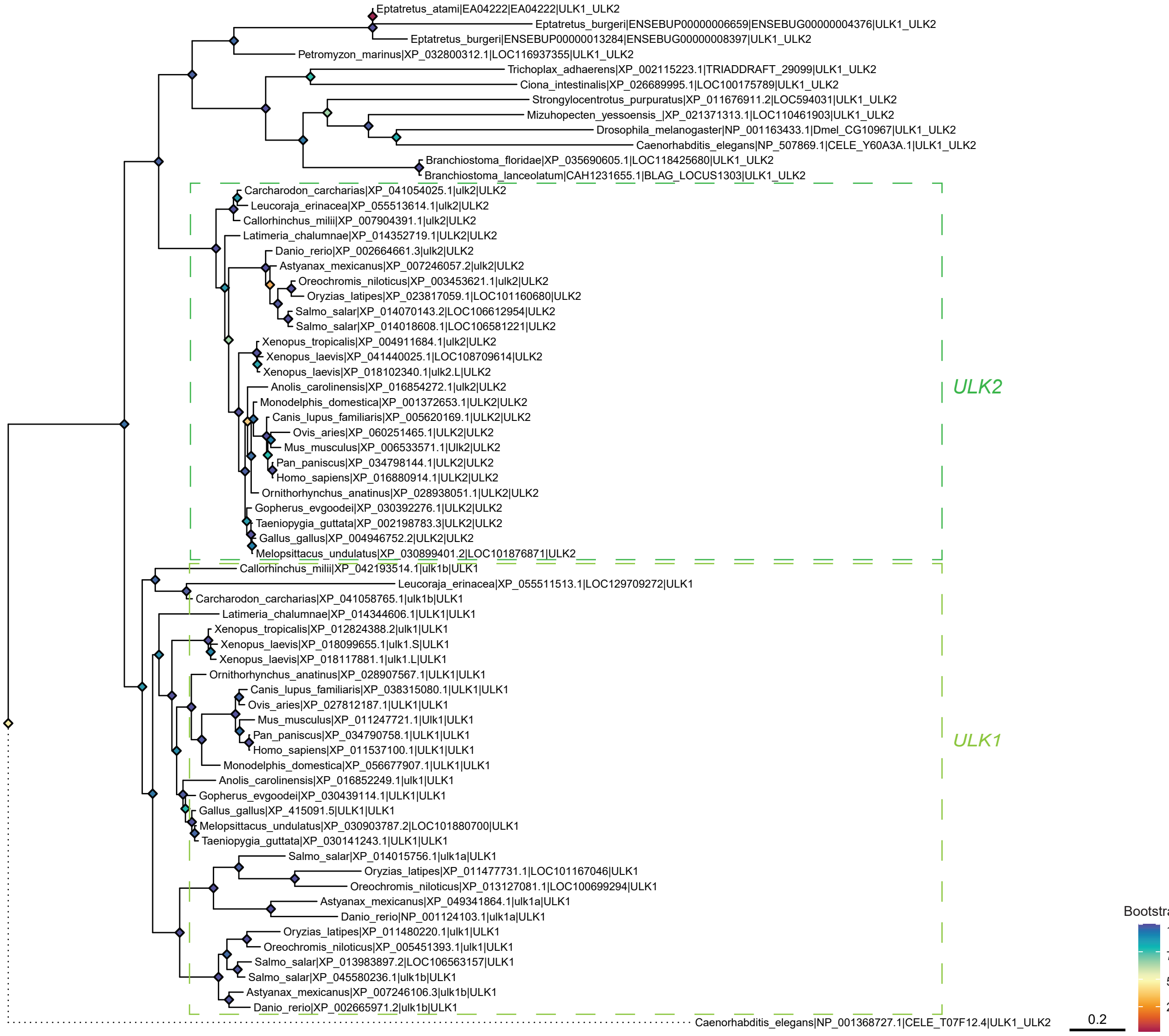

**Figure S1**. The phylogenetic trees of (**A**) *ULK-1* (**B**) *ATG9* (**C**) *BECN* (**D**) *ATG2* (**E**) *WIPI* (**F**) *ATG8* (including **i**, *GABARAP*s and **ii**, *LC3*s) (**G**) *ATG4*, and (**H**) *ATG16* homologs before and after gene duplications, manually rooted with five outgroup species, namely *Drosophila melanogaster*, *Caenorhabditis elegans*, *Strongylocentrotus purpuratus* (purple sea urchin), *Mizuhopecten yessoensis* (scallop), and *Trichoplax adhaerens*. Bootstrap values of the nodes are indicated by their color. Dotted lines indicate long branches that were manually shrunk out of proportion for display purposes. Mammalian *ATG9B* sequences and teleost fish *GABARAPL1* sequences on long branches are highlighted in purple. *BECN2* sequences (orange rectangle) have only been found in eutherians.

B

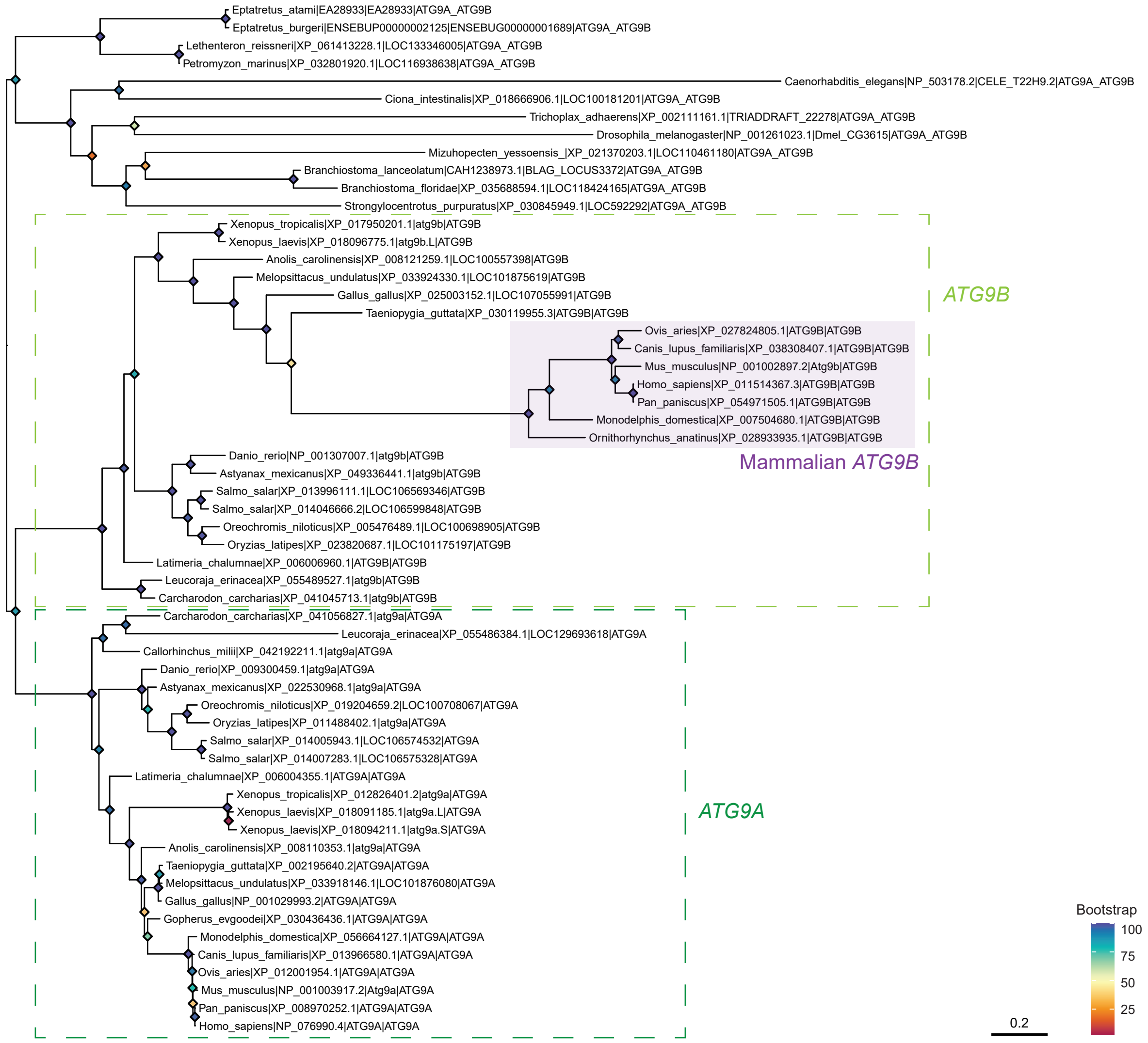

Eptatretus_atami|EA28933|EA28933|ATG9A_ATG9B
Eptatretus_burgeri|ENSEBUP00000002125|ENSEBUG00000001689|ATG9A_ATG9B
Lethenteron_reissneri|XP_061413228.1|LOC133346005|ATG9A_ATG9B
Petromyzon_marinus|XP_032801920.1|LOC116938638|ATG9A_ATG9B
Caenorhabditis_elegans|NP_503178.2|CELE_T22H9.2|ATG9A_ATG9B
Ciona_intestinalis|XP_018666906.1|LOC100181201|ATG9A_ATG9B
Trichoplax_adhaerens|XP_002111161.1|TRIADDRAFT_22278|ATG9A_ATG9B
Drosophila_melanogaster|NP_001261023.1|Dmel_CG3615|ATG9A_ATG9B
Mizuhopecten_yessoensis_|XP_021370203.1|LOC110461180|ATG9A_ATG9B
Branchiostoma_lanceolatum|CAH1238973.1|BLAG_LOCUS3372|ATG9A_ATG9B
Branchiostoma_floridae|XP_035688594.1|LOC118424165|ATG9A_ATG9B
Strongylocentrotus_purpuratus|XP_030845949.1|LOC592292|ATG9A_ATG9B
Xenopus_tropicalis|XP_017950201.1|atg9b|ATG9B
Xenopus_laevis|XP_018096775.1|atg9b.L|ATG9B
Anolis_carolinensis|XP_008121259.1|LOC100557398|ATG9B
Melopsittacus_undulatus|XP_033924330.1|LOC101875619|ATG9B
Gallus_gallus|XP_025003152.1|LOC107055991|ATG9B
Taeniopygia_guttata|XP_030119955.3|ATG9B|ATG9B
ATG9B
Ovis_aries|XP_027824805.1|ATG9B|ATG9B
Canis_lupus_familiaris|XP_038308407.1|ATG9B|ATG9B
Mus_musculus|NP_001002897.2|Atg9b|ATG9B
Homo_sapiens|XP_011514367.3|ATG9B|ATG9B
Pan_paniscus|XP_054971505.1|ATG9B|ATG9B
Monodelphis_domestica|XP_007504680.1|ATG9B|ATG9B
Ornithorhynchus_anatinus|XP_028933935.1|ATG9B|ATG9B
Mammalian ATG9B
Danio_rerio|NP_001307007.1|atg9b|ATG9B
Astyanax_mexicanus|XP_049336441.1|atg9b|ATG9B
Salmo_salar|XP_013996111.1|LOC106569346|ATG9B
Salmo_salar|XP_014046666.2|LOC106599848|ATG9B
Oreochromis_niloticus|XP_005476489.1|LOC100698905|ATG9B
Oryzias_latipes|XP_023820687.1|LOC101175197|ATG9B
Latimeria_chalumnae|XP_006006960.1|ATG9B|ATG9B
Leucoraja_erinacea|XP_055489527.1|atg9b|ATG9B
Carcharodon_carcharias|XP_041045713.1|atg9b|ATG9B
Carcharodon_carcharias|XP_041056827.1|atg9a|ATG9A
Leucoraja_erinacea|XP_055486384.1|LOC129693618|ATG9A
Callorhinchus_milii|XP_042192211.1|atg9a|ATG9A
Danio_rerio|XP_009300459.1|atg9a|ATG9A
Astyanax_mexicanus|XP_022530968.1|atg9a|ATG9A
Oreochromis_niloticus|XP_019204659.2|LOC100708067|ATG9A
Oryzias_latipes|XP_011488402.1|atg9a|ATG9A
Salmo_salar|XP_014005943.1|LOC106574532|ATG9A
Salmo_salar|XP_014007283.1|LOC106575328|ATG9A
Latimeria_chalumnae|XP_006004355.1|ATG9A|ATG9A
Xenopus_tropicalis|XP_012826401.2|atg9a|ATG9A
Xenopus_laevis|XP_018091185.1|atg9a.L|ATG9A
Xenopus_laevis|XP_018094211.1|atg9a.S|ATG9A
ATG9A
Anolis_carolinensis|XP_008110353.1|atg9a|ATG9A
Taeniopygia_guttata|XP_002195640.2|ATG9A|ATG9A
Melopsittacus_undulatus|XP_033918146.1|LOC101876080|ATG9A
Gallus_gallus|NP_001029993.2|ATG9A|ATG9A
Gopherus_evgoodei|XP_030436436.1|ATG9A|ATG9A
Monodelphis_domestica|XP_056664127.1|ATG9A|ATG9A
Canis_lupus_familiaris|XP_013966580.1|ATG9A|ATG9A
Ovis_aries|XP_012001954.1|ATG9A|ATG9A
Mus_musculus|NP_001003917.2|Atg9a|ATG9A
Pan_paniscus|XP_008970252.1|ATG9A|ATG9A
Homo_sapiens|NP_076990.4|ATG9A|ATG9A
Bootstrap
100
75
50
25
0.2

**C**

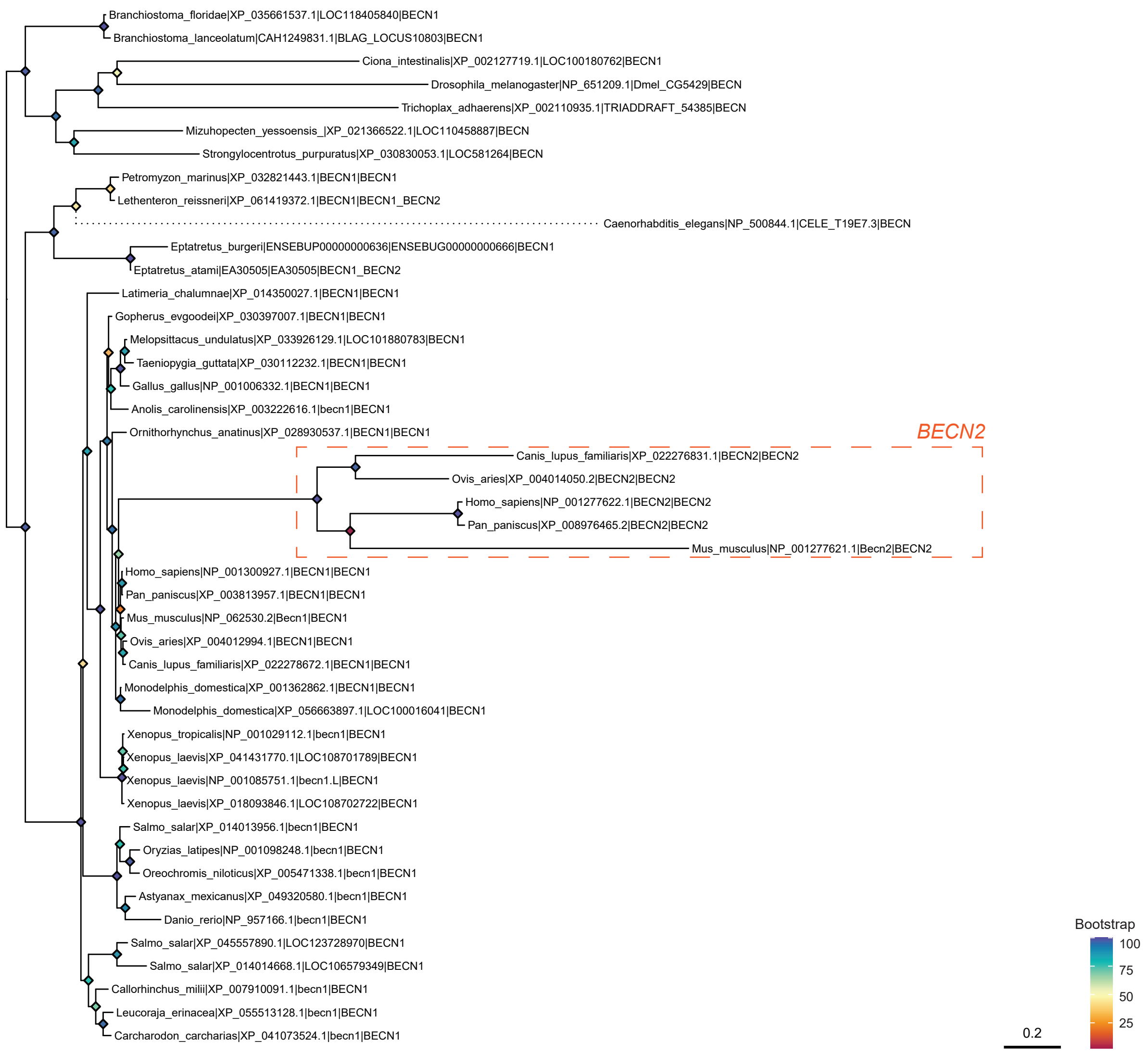

Branchiostoma_floridae|XP_035661537.1|LOC118405840|BECN1
Branchiostoma_lanceolatum|CAH1249831.1|BLAG_LOCUS10803|BECN1
Ciona_intestinalis|XP_002127719.1|LOC100180762|BECN1
Drosophila_melanogaster|NP_651209.1|Dmel_CG5429|BECN
Trichoplax_adhaerens|XP_002110935.1|TRIADDRAFT_54385|BECN
Mizuhopecten_yessoensis_|XP_021366522.1|LOC110458887|BECN
Strongylocentrotus_purpuratus|XP_030830053.1|LOC581264|BECN
Petromyzon_marinus|XP_032821443.1|BECN1|BECN1
Lethenteron_reissneri|XP_061419372.1|BECN1|BECN1_BECN2
Caenorhabditis_elegans|NP_500844.1|CELE_T19E7.3|BECN
Eptatretus_burgeri|ENSEBUP00000000636|ENSEBUG00000000666|BECN1
Eptatretus_atami|EA30505|EA30505|BECN1_BECN2
Latimeria_chalumnae|XP_014350027.1|BECN1|BECN1
Gopherus_evgoodei|XP_030397007.1|BECN1|BECN1
Melopsittacus_undulatus|XP_033926129.1|LOC101880783|BECN1
Taeniopygia_guttata|XP_030112232.1|BECN1|BECN1
Gallus_gallus|NP_001006332.1|BECN1|BECN1
Anolis_carolinensis|XP_003222616.1|becn1|BECN1
Ornithorhynchus_anatinus|XP_028930537.1|BECN1|BECN1
BECN2
Canis_lupus_familiaris|XP_022276831.1|BECN2|BECN2
Ovis_aries|XP_004014050.2|BECN2|BECN2
Homo_sapiens|NP_001277622.1|BECN2|BECN2
Pan_paniscus|XP_008976465.2|BECN2|BECN2
Mus_musculus|NP_001277621.1|Becn2|BECN2
Homo_sapiens|NP_001300927.1|BECN1|BECN1
Pan_paniscus|XP_003813957.1|BECN1|BECN1
Mus_musculus|NP_062530.2|Becn1|BECN1
Ovis_aries|XP_004012994.1|BECN1|BECN1
Canis_lupus_familiaris|XP_022278672.1|BECN1|BECN1
Monodelphis_domestica|XP_001362862.1|BECN1|BECN1
Monodelphis_domestica|XP_056663897.1|LOC100016041|BECN1
Xenopus_tropicalis|NP_001029112.1|becn1|BECN1
Xenopus_laevis|XP_041431770.1|LOC108701789|BECN1
Xenopus_laevis|NP_001085751.1|becn1.L|BECN1
Xenopus_laevis|XP_018093846.1|LOC108702722|BECN1
Salmo_salar|XP_014013956.1|becn1|BECN1
Oryzias_latipes|NP_001098248.1|becn1|BECN1
Oreochromis_niloticus|XP_005471338.1|becn1|BECN1
Astyanax_mexicanus|XP_049320580.1|becn1|BECN1
Danio_rerio|NP_957166.1|becn1|BECN1
Salmo_salar|XP_045557890.1|LOC123728970|BECN1
Salmo_salar|XP_014014668.1|LOC106579349|BECN1
Callorhinchus_milii|XP_007910091.1|becn1|BECN1
Leucoraja_erinacea|XP_055513128.1|becn1|BECN1
Carcharodon_carcharias|XP_041073524.1|becn1|BECN1
Bootstrap
100
75
50
25
0.2

D

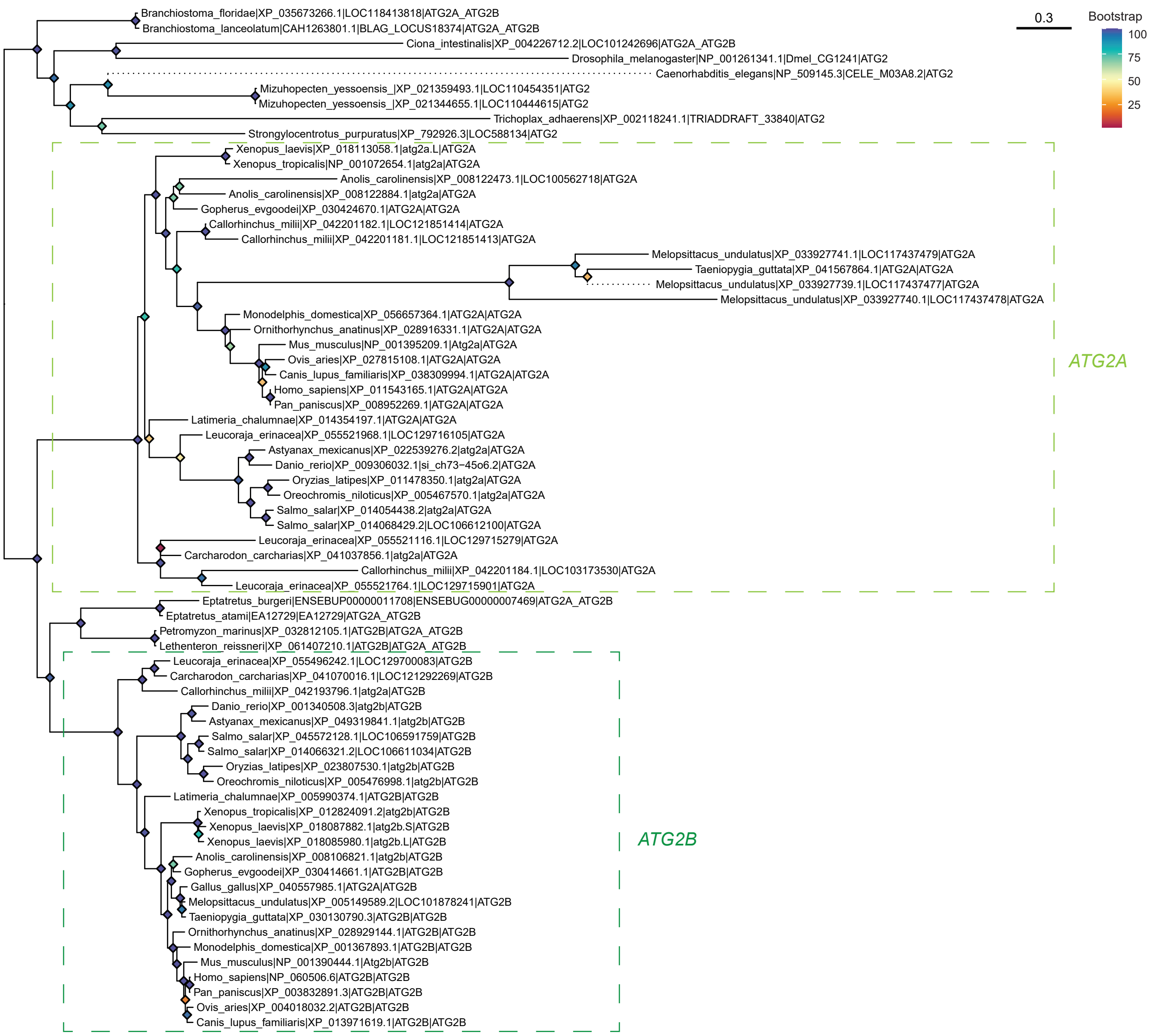
0.3
Bootstrap
100
75
50
25
Branchiostoma_floridae|XP_035673266.1|LOC118413818|ATG2A_ATG2B
Branchiostoma_lanceolatum|CAH1263801.1|BLAG_LOCUS18374|ATG2A_ATG2B
Ciona_intestinalis|XP_004226712.2|LOC101242696|ATG2A_ATG2B
Drosophila_melanogaster|NP_001261341.1|Dmel_CG1241|ATG2
Caenorhabditis_elegans|NP_509145.3|CELE_M03A8.2|ATG2
Mizuhopecten_yessoensis_|XP_021359493.1|LOC110454351|ATG2
Mizuhopecten_yessoensis_|XP_021344655.1|LOC110444615|ATG2
Trichoplax_adhaerens|XP_002118241.1|TRIADDRAFT_33840|ATG2
Strongylocentrotus_purpuratus|XP_792926.3|LOC588134|ATG2
Xenopus_laevis|XP_018113058.1|atg2a.L|ATG2A
Xenopus_tropicalis|NP_001072654.1|atg2a|ATG2A
Anolis_carolinensis|XP_008122473.1|LOC100562718|ATG2A
Anolis_carolinensis|XP_008122884.1|atg2a|ATG2A
Gopherus_evgoodei|XP_030424670.1|ATG2A|ATG2A
Callorhinchus_milii|XP_042201182.1|LOC121851414|ATG2A
Callorhinchus_milii|XP_042201181.1|LOC121851413|ATG2A
Melopsittacus_undulatus|XP_033927741.1|LOC117437479|ATG2A
Taeniopygia_guttata|XP_041567864.1|ATG2A|ATG2A
Melopsittacus_undulatus|XP_033927739.1|LOC117437477|ATG2A
Melopsittacus_undulatus|XP_033927740.1|LOC117437478|ATG2A
Monodelphis_domestica|XP_056657364.1|ATG2A|ATG2A
Ornithorhynchus_anatinus|XP_028916331.1|ATG2A|ATG2A
Mus_musculus|NP_001395209.1|Atg2a|ATG2A
Ovis_aries|XP_027815108.1|ATG2A|ATG2A
Canis_lupus_familiaris|XP_038309994.1|ATG2A|ATG2A
Homo_sapiens|XP_011543165.1|ATG2A|ATG2A
Pan_paniscus|XP_008952269.1|ATG2A|ATG2A
Latimeria_chalumnae|XP_014354197.1|ATG2A|ATG2A
Leucoraja_erinacea|XP_055521968.1|LOC129716105|ATG2A
Astyanax_mexicanus|XP_022539276.2|atg2a|ATG2A
Danio_rerio|XP_009306032.1|si_ch73-45o6.2|ATG2A
Oryzias_latipes|XP_011478350.1|atg2a|ATG2A
Oreochromis_niloticus|XP_005467570.1|atg2a|ATG2A
Salmo_salar|XP_014054438.2|atg2a|ATG2A
Salmo_salar|XP_014068429.2|LOC106612100|ATG2A
Leucoraja_erinacea|XP_055521116.1|LOC129715279|ATG2A
Carcharodon_carcharias|XP_041037856.1|atg2a|ATG2A
Callorhinchus_milii|XP_042201184.1|LOC103173530|ATG2A
Leucoraja_erinacea|XP_055521764.1|LOC129715901|ATG2A
ATG2A
Eptatretus_burgeri|ENSEBUP00000011708|ENSEBUG00000007469|ATG2A_ATG2B
Eptatretus_atami|EA12729|EA12729|ATG2A_ATG2B
Petromyzon_marinus|XP_032812105.1|ATG2B|ATG2A_ATG2B
Lethenteron_reissneri|XP_061407210.1|ATG2B|ATG2A_ATG2B
Leucoraja_erinacea|XP_055496242.1|LOC129700083|ATG2B
Carcharodon_carcharias|XP_041070016.1|LOC121292269|ATG2B
Callorhinchus_milii|XP_042193796.1|atg2a|ATG2B
Danio_rerio|XP_001340508.3|atg2b|ATG2B
Astyanax_mexicanus|XP_049319841.1|atg2b|ATG2B
Salmo_salar|XP_045572128.1|LOC106591759|ATG2B
Salmo_salar|XP_014066321.2|LOC106611034|ATG2B
Oryzias_latipes|XP_023807530.1|atg2b|ATG2B
Oreochromis_niloticus|XP_005476998.1|atg2b|ATG2B
Latimeria_chalumnae|XP_005990374.1|ATG2B|ATG2B
Xenopus_tropicalis|XP_012824091.2|atg2b|ATG2B
Xenopus_laevis|XP_018087882.1|atg2b.S|ATG2B
Xenopus_laevis|XP_018085980.1|atg2b.L|ATG2B
Anolis_carolinensis|XP_008106821.1|atg2b|ATG2B
Gopherus_evgoodei|XP_030414661.1|ATG2B|ATG2B
Gallus_gallus|XP_040557985.1|ATG2A|ATG2B
Melopsittacus_undulatus|XP_005149589.2|LOC101878241|ATG2B
Taeniopygia_guttata|XP_030130790.3|ATG2B|ATG2B
Ornithorhynchus_anatinus|XP_028929144.1|ATG2B|ATG2B
Monodelphis_domestica|XP_001367893.1|ATG2B|ATG2B
Mus_musculus|NP_001390444.1|Atg2b|ATG2B
Homo_sapiens|NP_060506.6|ATG2B|ATG2B
Pan_paniscus|XP_003832891.3|ATG2B|ATG2B
Ovis_aries|XP_004018032.2|ATG2B|ATG2B
Canis_lupus_familiaris|XP_013971619.1|ATG2B|ATG2B
ATG2B

**E**

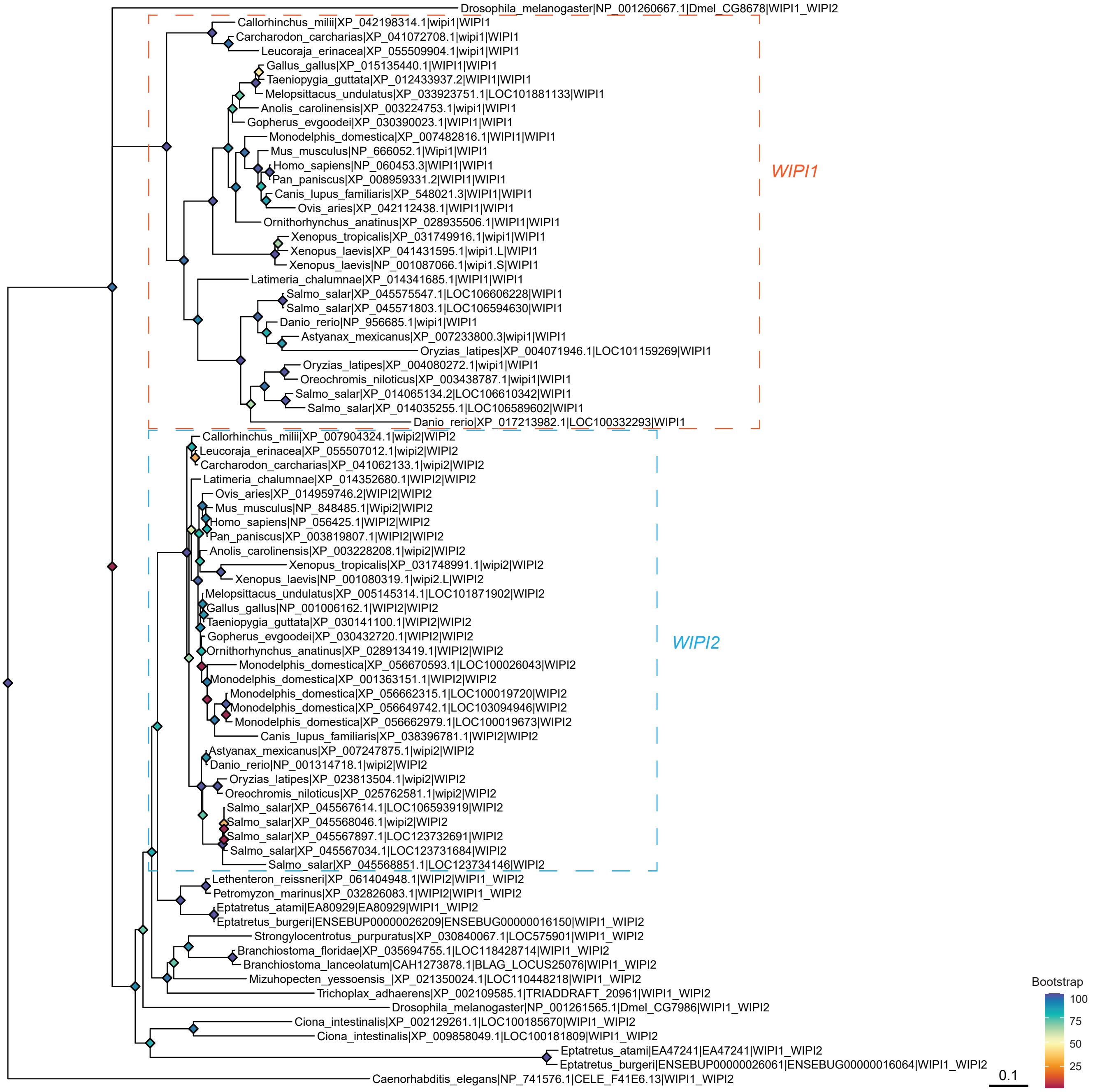
Drosophila_melanogaster|NP_001260667.1|Dmel_CG8678|WIPI1_WIPI2
Callorhinchus_milii|XP_042198314.1|wipi1|WIPI1
Carcharodon_carcharias|XP_041072708.1|wipi1|WIPI1
Leucoraja_erinacea|XP_055509904.1|wipi1|WIPI1
Gallus_gallus|XP_015135440.1|WIPI1|WIPI1
Taeniopygia_guttata|XP_012433937.2|WIPI1|WIPI1
Melopsittacus_undulatus|XP_033923751.1|LOC101881133|WIPI1
Anolis_carolinensis|XP_003224753.1|wipi1|WIPI1
Gopherus_evgoodei|XP_030390023.1|WIPI1|WIPI1
Monodelphis_domestica|XP_007482816.1|WIPI1|WIPI1
Mus_musculus|NP_666052.1|Wipi1|WIPI1
Homo_sapiens|NP_060453.3|WIPI1|WIPI1
Pan_paniscus|XP_008959331.2|WIPI1|WIPI1
Canis_lupus_familiaris|XP_548021.3|WIPI1|WIPI1
Ovis_aries|XP_042112438.1|WIPI1|WIPI1
Ornithorhynchus_anatinus|XP_028935506.1|WIPI1|WIPI1
Xenopus_tropicalis|XP_031749916.1|wipi1|WIPI1
Xenopus_laevis|XP_041431595.1|wipi1.L|WIPI1
Xenopus_laevis|NP_001087066.1|wipi1.S|WIPI1
Latimeria_chalumnae|XP_014341685.1|WIPI1|WIPI1
Salmo_salar|XP_045575547.1|LOC106606228|WIPI1
Salmo_salar|XP_045571803.1|LOC106594630|WIPI1
Danio_rerio|NP_956685.1|wipi1|WIPI1
Astyanax_mexicanus|XP_007233800.3|wipi1|WIPI1
Oryzias_latipes|XP_004071946.1|LOC101159269|WIPI1
Oryzias_latipes|XP_004080272.1|wipi1|WIPI1
Oreochromis_niloticus|XP_003438787.1|wipi1|WIPI1
Salmo_salar|XP_014065134.2|LOC106610342|WIPI1
Salmo_salar|XP_014035255.1|LOC106589602|WIPI1
Danio_rerio|XP_017213982.1|LOC100332293|WIPI1
WIPI1
Callorhinchus_milii|XP_007904324.1|wipi2|WIPI2
Leucoraja_erinacea|XP_055507012.1|wipi2|WIPI2
Carcharodon_carcharias|XP_041062133.1|wipi2|WIPI2
Latimeria_chalumnae|XP_014352680.1|WIPI2|WIPI2
Ovis_aries|XP_014959746.2|WIPI2|WIPI2
Mus_musculus|NP_848485.1|Wipi2|WIPI2
Homo_sapiens|NP_056425.1|WIPI2|WIPI2
Pan_paniscus|XP_003819807.1|WIPI2|WIPI2
Anolis_carolinensis|XP_003228208.1|wipi2|WIPI2
Xenopus_tropicalis|XP_031748991.1|wipi2|WIPI2
Xenopus_laevis|NP_001080319.1|wipi2.L|WIPI2
Melopsittacus_undulatus|XP_005145314.1|LOC101871902|WIPI2
Gallus_gallus|NP_001006162.1|WIPI2|WIPI2
Taeniopygia_guttata|XP_030141100.1|WIPI2|WIPI2
Gopherus_evgoodei|XP_030432720.1|WIPI2|WIPI2
Ornithorhynchus_anatinus|XP_028913419.1|WIPI2|WIPI2
Monodelphis_domestica|XP_056670593.1|LOC100026043|WIPI2
Monodelphis_domestica|XP_001363151.1|WIPI2|WIPI2
Monodelphis_domestica|XP_056662315.1|LOC100019720|WIPI2
Monodelphis_domestica|XP_056649742.1|LOC103094946|WIPI2
Monodelphis_domestica|XP_056662979.1|LOC100019673|WIPI2
Canis_lupus_familiaris|XP_038396781.1|WIPI2|WIPI2
Astyanax_mexicanus|XP_007247875.1|wipi2|WIPI2
Danio_rerio|NP_001314718.1|wipi2|WIPI2
Oryzias_latipes|XP_023813504.1|wipi2|WIPI2
Oreochromis_niloticus|XP_025762581.1|wipi2|WIPI2
Salmo_salar|XP_045567614.1|LOC106593919|WIPI2
Salmo_salar|XP_045568046.1|wipi2|WIPI2
Salmo_salar|XP_045567897.1|LOC123732691|WIPI2
Salmo_salar|XP_045567034.1|LOC123731684|WIPI2
Salmo_salar|XP_045568851.1|LOC123734146|WIPI2
WIPI2
Lethenteron_reissneri|XP_061404948.1|WIPI2|WIPI1_WIPI2
Petromyzon_marinus|XP_032826083.1|WIPI2|WIPI1_WIPI2
Eptatretus_atami|EA80929|EA80929|WIPI1_WIPI2
Eptatretus_burgeri|ENSEBUP00000026209|ENSEBUG00000016150|WIPI1_WIPI2
Strongylocentrotus_purpuratus|XP_030840067.1|LOC575901|WIPI1_WIPI2
Branchiostoma_floridae|XP_035694755.1|LOC118428714|WIPI1_WIPI2
Branchiostoma_lanceolatum|CAH1273878.1|BLAG_LOCUS25076|WIPI1_WIPI2
Mizuhopecten_yessoensis_|XP_021350024.1|LOC110448218|WIPI1_WIPI2
Trichoplax_adhaerens|XP_002109585.1|TRIADDRAFT_20961|WIPI1_WIPI2
Drosophila_melanogaster|NP_001261565.1|Dmel_CG7986|WIPI1_WIPI2
Ciona_intestinalis|XP_002129261.1|LOC100185670|WIPI1_WIPI2
Ciona_intestinalis|XP_009858049.1|LOC100181809|WIPI1_WIPI2
Eptatretus_atami|EA47241|EA47241|WIPI1_WIPI2
Eptatretus_burgeri|ENSEBUP00000026061|ENSEBUG00000016064|WIPI1_WIPI2
Caenorhabditis_elegans|NP_741576.1|CELE_F41E6.13|WIPI1_WIPI2
0.1
Bootstrap
100
75
50
25

**F i**

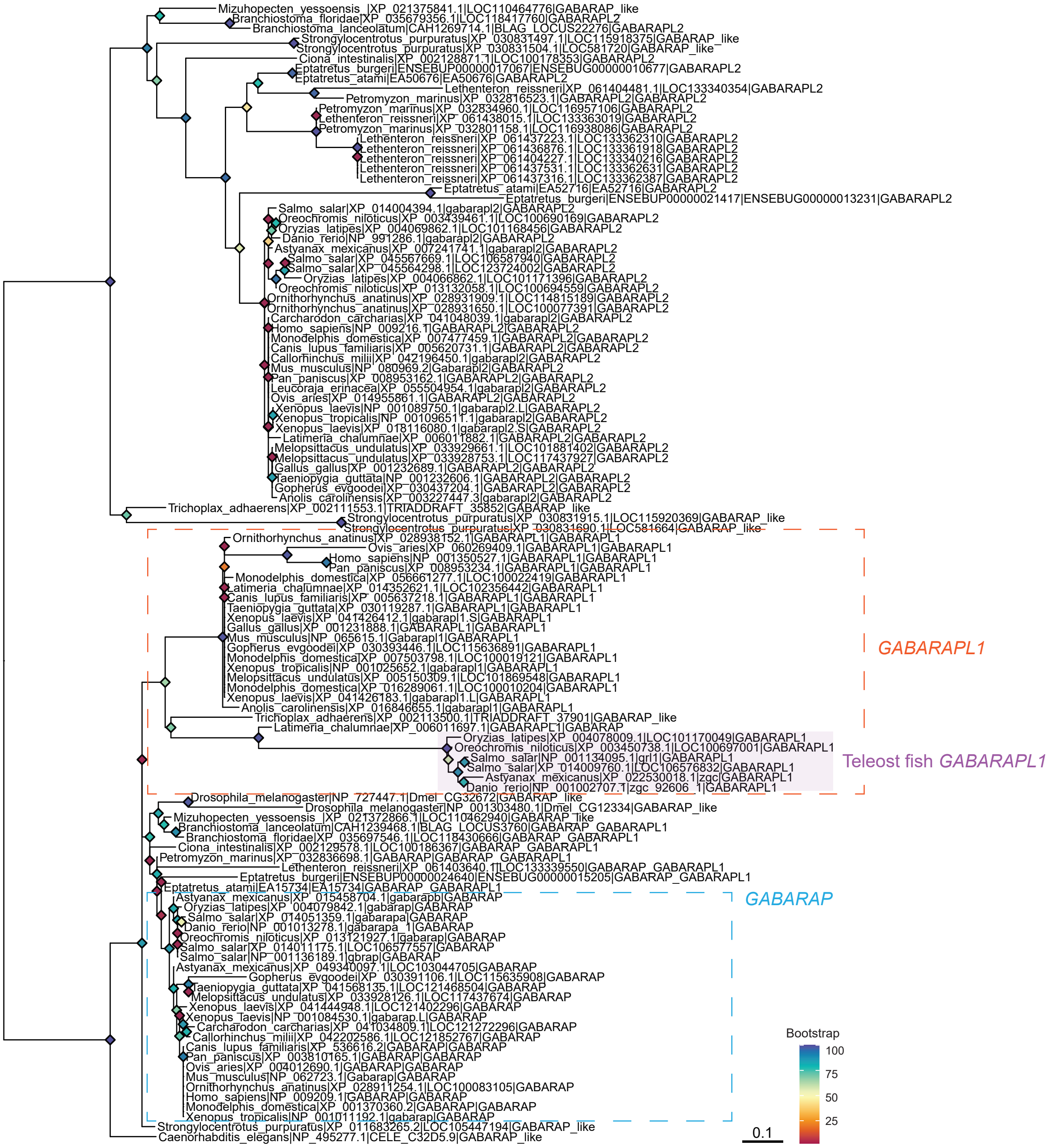

Mizuhopecten_yessoensis_|XP_021375841.1|LOC110464776|GABARAP_like
Branchiostoma_floridae|XP_035679356.1|LOC118417760|GABARAPL2
Branchiostoma_lanceolatum|CAH1269714.1|BLAG_LOCUS22276|GABARAPL2
Strongylocentrotus_purpuratus|XP_030831497.1|LOC115918375|GABARAP_like
Strongylocentrotus_purpuratus|XP_030831504.1|LOC581720|GABARAP_like
Ciona_intestinalis|XP_002128871.1|LOC100178353|GABARAPL2
Eptatretus_burgeri|ENSEBUP00000017067|ENSEBUG00000010677|GABARAPL2
Eptatretus_atami|EA50676|EA50676|GABARAPL2
Lethenteron_reissneri|XP_061404481.1|LOC133340354|GABARAPL2
Petromyzon_marinus|XP_032816523.1|GABARAPL2|GABARAPL2
Petromyzon_marinus|XP_032834960.1|LOC116957106|GABARAPL2
Lethenteron_reissneri|XP_061438015.1|LOC133363019|GABARAPL2
Petromyzon_marinus|XP_032801158.1|LOC116938086|GABARAPL2
Lethenteron_reissneri|XP_061437223.1|LOC133362310|GABARAPL2
Lethenteron_reissneri|XP_061436876.1|LOC133361918|GABARAPL2
Lethenteron_reissneri|XP_061404227.1|LOC133340216|GABARAPL2
Lethenteron_reissneri|XP_061437531.1|LOC133362631|GABARAPL2
Lethenteron_reissneri|XP_061437316.1|LOC133362387|GABARAPL2
Eptatretus_atami|EA52716|EA52716|GABARAPL2
Eptatretus_burgeri|ENSEBUP00000021417|ENSEBUG00000013231|GABARAPL2
Salmo_salar|XP_014004394.1|gabarapl2|GABARAPL2
Oreochromis_niloticus|XP_003439461.1|LOC100690169|GABARAPL2
Oryzias_latipes|XP_004069862.1|LOC101168456|GABARAPL2
Danio_rerio|NP_991286.1|gabarapl2|GABARAPL2
Astyanax_mexicanus|XP_007241741.1|gabarapl2|GABARAPL2
Salmo_salar|XP_045567669.1|LOC106587940|GABARAPL2
Salmo_salar|XP_045564298.1|LOC123724002|GABARAPL2
Oryzias_latipes|XP_004066862.1|LOC101171396|GABARAPL2
Oreochromis_niloticus|XP_013132058.1|LOC100694559|GABARAPL2
Ornithorhynchus_anatinus|XP_028931909.1|LOC114815189|GABARAPL2
Ornithorhynchus_anatinus|XP_028931650.1|LOC100077391|GABARAPL2
Carcharodon_carcharias|XP_041048039.1|gabarapl2|GABARAPL2
Homo_sapiens|NP_009216.1|GABARAPL2|GABARAPL2
Monodelphis_domestica|XP_007477459.1|GABARAPL2|GABARAPL2
Canis_lupus_familiaris|XP_005620731.1|GABARAPL2|GABARAPL2
Callorhinchus_milii|XP_042196450.1|gabarapl2|GABARAPL2
Mus_musculus|NP_080969.2|Gabarapl2|GABARAPL2
Pan_paniscus|XP_008953162.1|GABARAPL2|GABARAPL2
Leucoraja_erinacea|XP_055504954.1|gabarapl2|GABARAPL2
Ovis_aries|XP_014955861.1|GABARAPL2|GABARAPL2
Xenopus_laevis|NP_001089750.1|gabarapl2.L|GABARAPL2
Xenopus_tropicalis|NP_001096511.1|gabarapl2|GABARAPL2
Xenopus_laevis|XP_018116080.1|gabarapl2.S|GABARAPL2
Latimeria_chalumnae|XP_006011882.1|GABARAPL2|GABARAPL2
Melopsittacus_undulatus|XP_033929661.1|LOC101881402|GABARAPL2
Melopsittacus_undulatus|XP_033928753.1|LOC117437927|GABARAPL2
Gallus_gallus|XP_001232689.1|GABARAPL2|GABARAPL2
Taeniopygia_guttata|NP_001232606.1|GABARAPL2|GABARAPL2
Gopherus_evgoodei|XP_030437204.1|GABARAPL2|GABARAPL2
Anolis_carolinensis|XP_003227447.3|gabarapl2|GABARAPL2
Trichoplax_adhaerens|XP_002111553.1|TRIADDRAFT_35852|GABARAP_like
Strongylocentrotus_purpuratus|XP_030831915.1|LOC115920369|GABARAP_like
Strongylocentrotus_purpuratus|XP_030831690.1|LOC581664|GABARAP_like
Ornithorhynchus_anatinus|XP_028938152.1|GABARAPL1|GABARAPL1
Ovis_aries|XP_060269409.1|GABARAPL1|GABARAPL1
Homo_sapiens|NP_001350527.1|GABARAPL1|GABARAPL1
Pan_paniscus|XP_008953234.1|GABARAPL1|GABARAPL1
Monodelphis_domestica|XP_056661277.1|LOC100022419|GABARAPL1
Latimeria_chalumnae|XP_014352621.1|LOC102356442|GABARAPL1
Canis_lupus_familiaris|XP_005637218.1|GABARAPL1|GABARAPL1
Taeniopygia_guttata|XP_030119287.1|GABARAPL1|GABARAPL1
Xenopus_laevis|XP_041426412.1|gabarapl1.S|GABARAPL1
Gallus_gallus|XP_001231888.1|GABARAPL1|GABARAPL1
Mus_musculus|NP_065615.1|Gabarapl1|GABARAPL1
Gopherus_evgoodei|XP_030393446.1|LOC115636891|GABARAPL1
Monodelphis_domestica|XP_007503798.1|LOC100019121|GABARAPL1
Xenopus_tropicalis|NP_001025652.1|gabarapl1|GABARAPL1
Melopsittacus_undulatus|XP_005150309.1|LOC101869548|GABARAPL1
Monodelphis_domestica|XP_016289061.1|LOC100010204|GABARAPL1
Xenopus_laevis|XP_041426183.1|gabarapl1.L|GABARAPL1
Anolis_carolinensis|XP_016846655.1|gabarapl1|GABARAPL1
Trichoplax_adhaerens|XP_002113500.1|TRIADDRAFT_37901|GABARAP_like
Latimeria_chalumnae|XP_006011697.1|GABARAPL1|GABARAP
Oryzias_latipes|XP_004078009.1|LOC101170049|GABARAPL1
Oreochromis_niloticus|XP_003450738.1|LOC100697001|GABARAPL1
Salmo_salar|NP_001134095.1|gri1|GABARAPL1
Salmo_salar|XP_014009760.1|LOC106576832|GABARAPL1
Astyanax_mexicanus|XP_022530018.1|zgc|GABARAPL1
Danio_rerio|NP_001002707.1|zgc_92606_1|GABARAPL1
GABARAPL1
Teleost fish GABARAPL1
Drosophila_melanogaster|NP_727447.1|Dmel_CG32672|GABARAP_like
Drosophila_melanogaster|NP_001303480.1|Dmel_CG12334|GABARAP_like
Mizuhopecten_yessoensis_|XP_021372866.1|LOC110462940|GABARAP_like
Branchiostoma_lanceolatum|CAH1239468.1|BLAG_LOCUS3760|GABARAP_GABARAPL1
Branchiostoma_floridae|XP_035697546.1|LOC118430666|GABARAP_GABARAPL1
Ciona_intestinalis|XP_002129578.1|LOC100186367|GABARAP_GABARAPL1
Petromyzon_marinus|XP_032836698.1|GABARAP|GABARAP_GABARAPL1
Lethenteron_reissneri|XP_061403640.1|LOC133339550|GABARAP_GABARAPL1
Eptatretus_burgeri|ENSEBUP00000024640|ENSEBUG00000015205|GABARAP_GABARAPL1
Eptatretus_atami|EA15734|EA15734|GABARAP_GABARAPL1
GABARAP
Astyanax_mexicanus|XP_015458704.1|gabarapb|GABARAP
Oryzias_latipes|XP_004079842.1|gabarap|GABARAP
Salmo_salar|XP_014051359.1|gabarapa|GABARAP
Danio_rerio|NP_001013278.1|gabarapa_1|GABARAP
Oreochromis_niloticus|XP_013121927.1|gabarap|GABARAP
Salmo_salar|XP_014011175.1|LOC106577557|GABARAP
Salmo_salar|NP_001136189.1|gbrap|GABARAP
Astyanax_mexicanus|XP_049340097.1|LOC103044705|GABARAP
Gopherus_evgoodei|XP_030391106.1|LOC115635908|GABARAP
Taeniopygia_guttata|XP_041568135.1|LOC121468504|GABARAP
Melopsittacus_undulatus|XP_033928126.1|LOC117437674|GABARAP
Xenopus_laevis|XP_041444948.1|LOC121402296|GABARAP
Xenopus_laevis|NP_001084530.1|gabarap.L|GABARAP
Carcharodon_carcharias|XP_041034809.1|LOC121272296|GABARAP
Callorhinchus_milii|XP_042202586.1|LOC121852767|GABARAP
Canis_lupus_familiaris|XP_536616.2|GABARAP|GABARAP
Pan_paniscus|XP_003810165.1|GABARAP|GABARAP
Ovis_aries|XP_004012690.1|GABARAP|GABARAP
Mus_musculus|NP_062723.1|Gabarap|GABARAP
Ornithorhynchus_anatinus|XP_028911254.1|LOC100083105|GABARAP
Homo_sapiens|NP_009209.1|GABARAP|GABARAP
Monodelphis_domestica|XP_001370360.2|GABARAP|GABARAP
Xenopus_tropicalis|NP_001011192.1|gabarap|GABARAP
Strongylocentrotus_purpuratus|XP_011683265.2|LOC105447194|GABARAP_like
Caenorhabditis_elegans|NP_495277.1|CELE_C32D5.9|GABARAP_like
Bootstrap
100
75
50
25
0.1

**F ii**

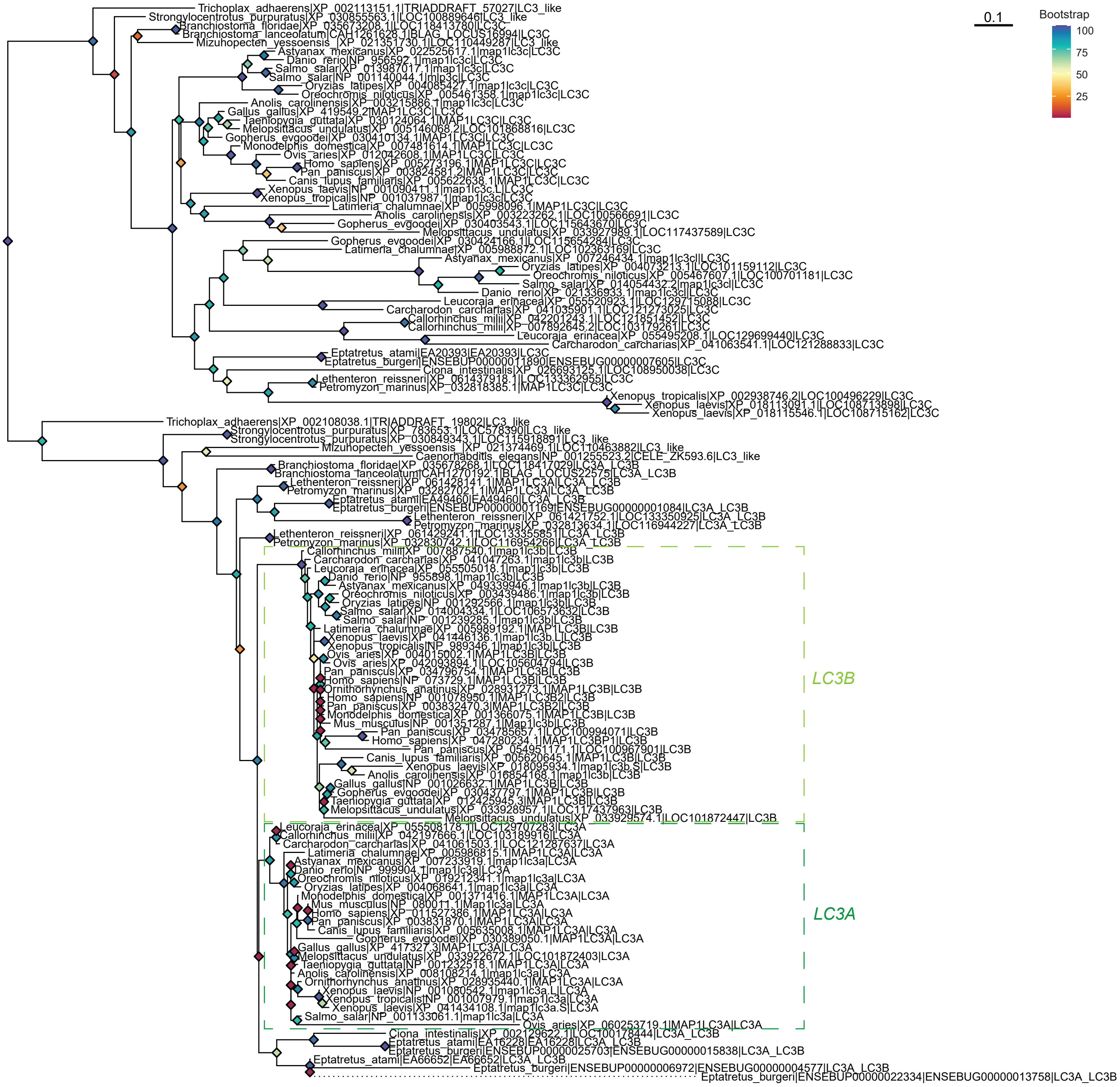

G

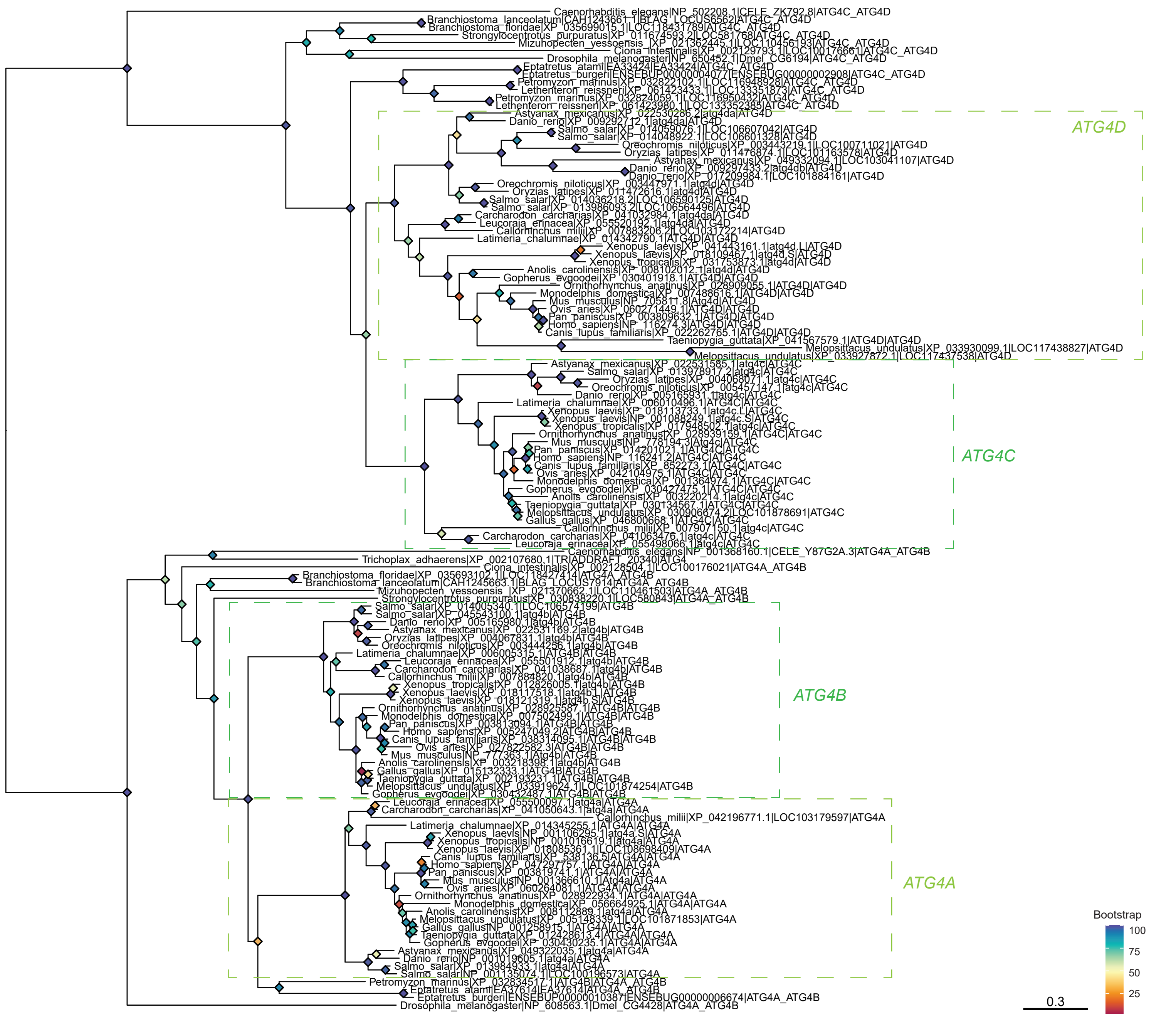

H

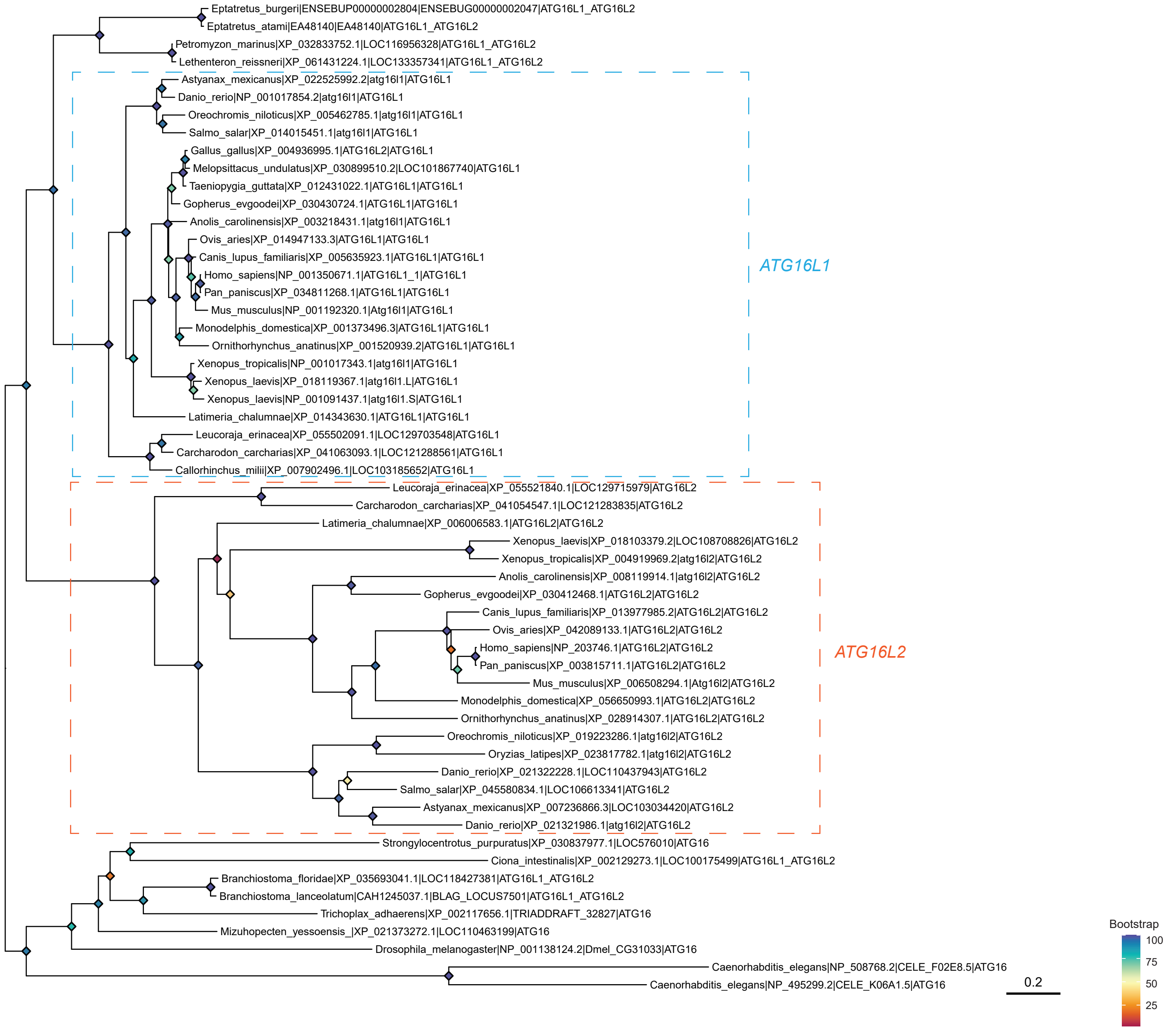

Eptatretus_burgeri|ENSEBUP00000002804|ENSEBUG00000002047|ATG16L1_ATG16L2
Eptatretus_atami|EA48140|EA48140|ATG16L1_ATG16L2
Petromyzon_marinus|XP_032833752.1|LOC116956328|ATG16L1_ATG16L2
Lethenteron_reissneri|XP_061431224.1|LOC133357341|ATG16L1_ATG16L2
Astyanax_mexicanus|XP_022525992.2|atg16l1|ATG16L1
Danio_rerio|NP_001017854.2|atg16l1|ATG16L1
Oreochromis_niloticus|XP_005462785.1|atg16l1|ATG16L1
Salmo_salar|XP_014015451.1|atg16l1|ATG16L1
Gallus_gallus|XP_004936995.1|ATG16L2|ATG16L1
Melopsittacus_undulatus|XP_030899510.2|LOC101867740|ATG16L1
Taeniopygia_guttata|XP_012431022.1|ATG16L1|ATG16L1
Gopherus_evgoodei|XP_030430724.1|ATG16L1|ATG16L1
Anolis_carolinensis|XP_003218431.1|atg16l1|ATG16L1
Ovis_aries|XP_014947133.3|ATG16L1|ATG16L1
Canis_lupus_familiaris|XP_005635923.1|ATG16L1|ATG16L1
Homo_sapiens|NP_001350671.1|ATG16L1_1|ATG16L1
Pan_paniscus|XP_034811268.1|ATG16L1|ATG16L1
Mus_musculus|NP_001192320.1|Atg16l1|ATG16L1
Monodelphis_domestica|XP_001373496.3|ATG16L1|ATG16L1
Ornithorhynchus_anatinus|XP_001520939.2|ATG16L1|ATG16L1
Xenopus_tropicalis|NP_001017343.1|atg16l1|ATG16L1
Xenopus_laevis|XP_018119367.1|atg16l1.L|ATG16L1
Xenopus_laevis|NP_001091437.1|atg16l1.S|ATG16L1
Latimeria_chalumnae|XP_014343630.1|ATG16L1|ATG16L1
Leucoraja_erinacea|XP_055502091.1|LOC129703548|ATG16L1
Carcharodon_carcharias|XP_041063093.1|LOC121288561|ATG16L1
Callorhinchus_milii|XP_007902496.1|LOC103185652|ATG16L1
ATG16L1
Leucoraja_erinacea|XP_055521840.1|LOC129715979|ATG16L2
Carcharodon_carcharias|XP_041054547.1|LOC121283835|ATG16L2
Latimeria_chalumnae|XP_006006583.1|ATG16L2|ATG16L2
Xenopus_laevis|XP_018103379.2|LOC108708826|ATG16L2
Xenopus_tropicalis|XP_004919969.2|atg16l2|ATG16L2
Anolis_carolinensis|XP_008119914.1|atg16l2|ATG16L2
Gopherus_evgoodei|XP_030412468.1|ATG16L2|ATG16L2
Canis_lupus_familiaris|XP_013977985.2|ATG16L2|ATG16L2
Ovis_aries|XP_042089133.1|ATG16L2|ATG16L2
Homo_sapiens|NP_203746.1|ATG16L2|ATG16L2
Pan_paniscus|XP_003815711.1|ATG16L2|ATG16L2
Mus_musculus|XP_006508294.1|Atg16l2|ATG16L2
Monodelphis_domestica|XP_056650993.1|ATG16L2|ATG16L2
Ornithorhynchus_anatinus|XP_028914307.1|ATG16L2|ATG16L2
Oreochromis_niloticus|XP_019223286.1|atg16l2|ATG16L2
Oryzias_latipes|XP_023817782.1|atg16l2|ATG16L2
Danio_rerio|XP_021322228.1|LOC110437943|ATG16L2
Salmo_salar|XP_045580834.1|LOC106613341|ATG16L2
Astyanax_mexicanus|XP_007236866.3|LOC103034420|ATG16L2
Danio_rerio|XP_021321986.1|atg16l2|ATG16L2
ATG16L2
Strongylocentrotus_purpuratus|XP_030837977.1|LOC576010|ATG16
Ciona_intestinalis|XP_002129273.1|LOC100175499|ATG16L1_ATG16L2
Branchiostoma_floridae|XP_035693041.1|LOC118427381|ATG16L1_ATG16L2
Branchiostoma_lanceolatum|CAH1245037.1|BLAG_LOCUS7501|ATG16L1_ATG16L2
Trichoplax_adhaerens|XP_002117656.1|TRIADDRAFT_32827|ATG16
Mizuhopecten_yessoensis_|XP_021373272.1|LOC110463199|ATG16
Drosophila_melanogaster|NP_001138124.2|Dmel_CG31033|ATG16
Caenorhabditis_elegans|NP_508768.2|CELE_F02E8.5|ATG16
Caenorhabditis_elegans|NP_495299.2|CELE_K06A1.5|ATG16
Bootstrap
100
75
50
25
0.2

Figure S2

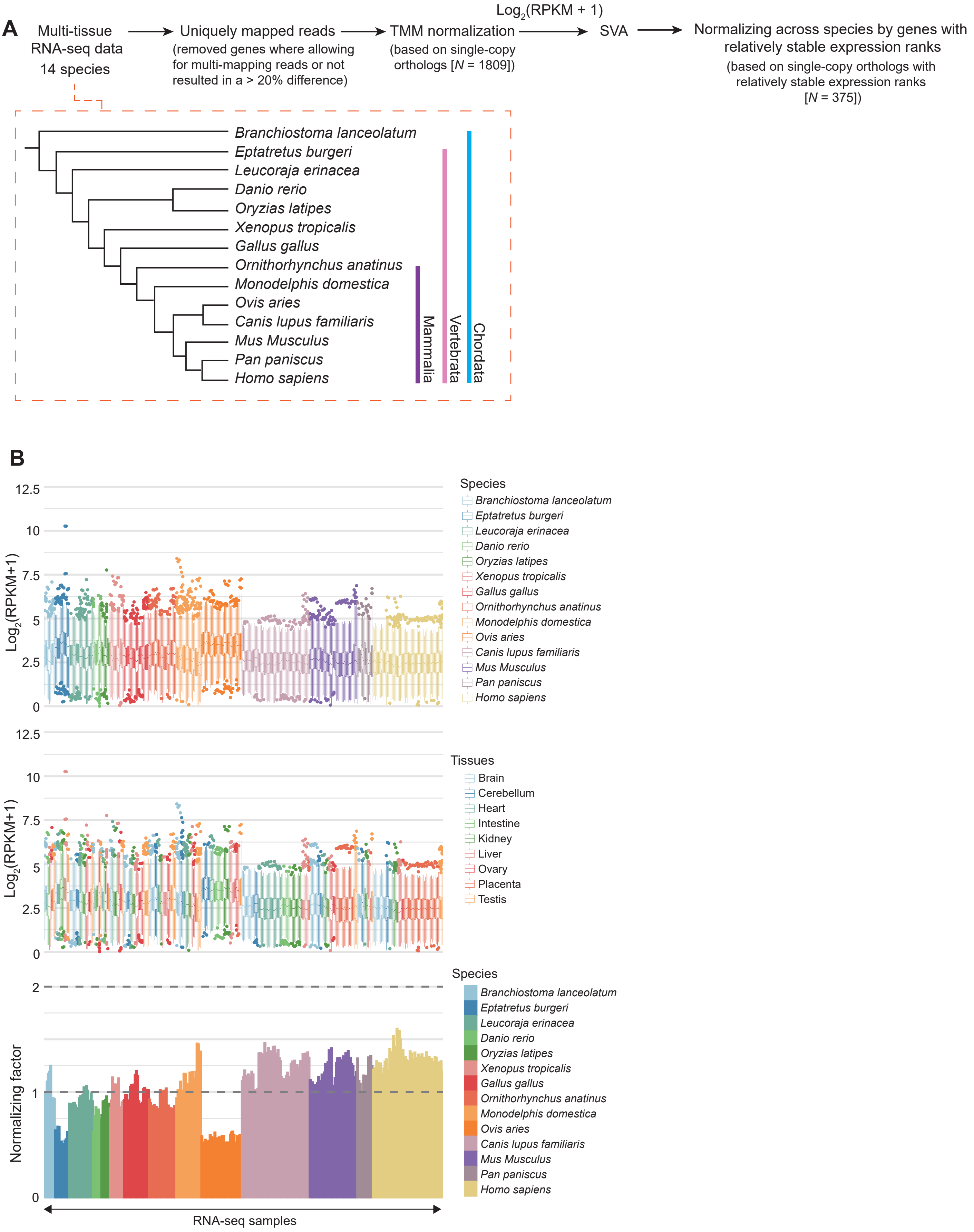

**Figure S2**. The quality control and normalization procedures for the RNA-seq data were applied to ensure that gene expression levels would be comparable among samples and species. (**A**) The overall quality control and normalization pipeline. Only the uniquely mapped reads were used in the analysis after removing genes in which counting all reads or counting only the uniquely mapped reads resulted in a >20% difference. In addition to the TMM normalization and the standard $\log_2$(RPKM+1) transformation, SVA was applied to remove hidden technical variables, and gene expression levels were further normalized based on the median of 375 single-copy orthologs with relatively stable expression ranks across samples and species. (**B**) The expression levels of 375 single-copy orthologs with relatively stable expression ranks across samples and species, colored by species (top) and tissue type (middle), and the normalizing factors calculated based on the median expression levels of these genes (bottom; all other samples were normalized to the leftmost sample). (**C**) SVA increased Pearson's r coefficient between samples of the same tissue types (calculated using all available genes). In each heatmap, each row and column represents one sample, and the samples are grouped by tissue type in a predefined order. For species in which SVA was not applied (see Materials and Methods for details), Pearson's r without SVA is shown.

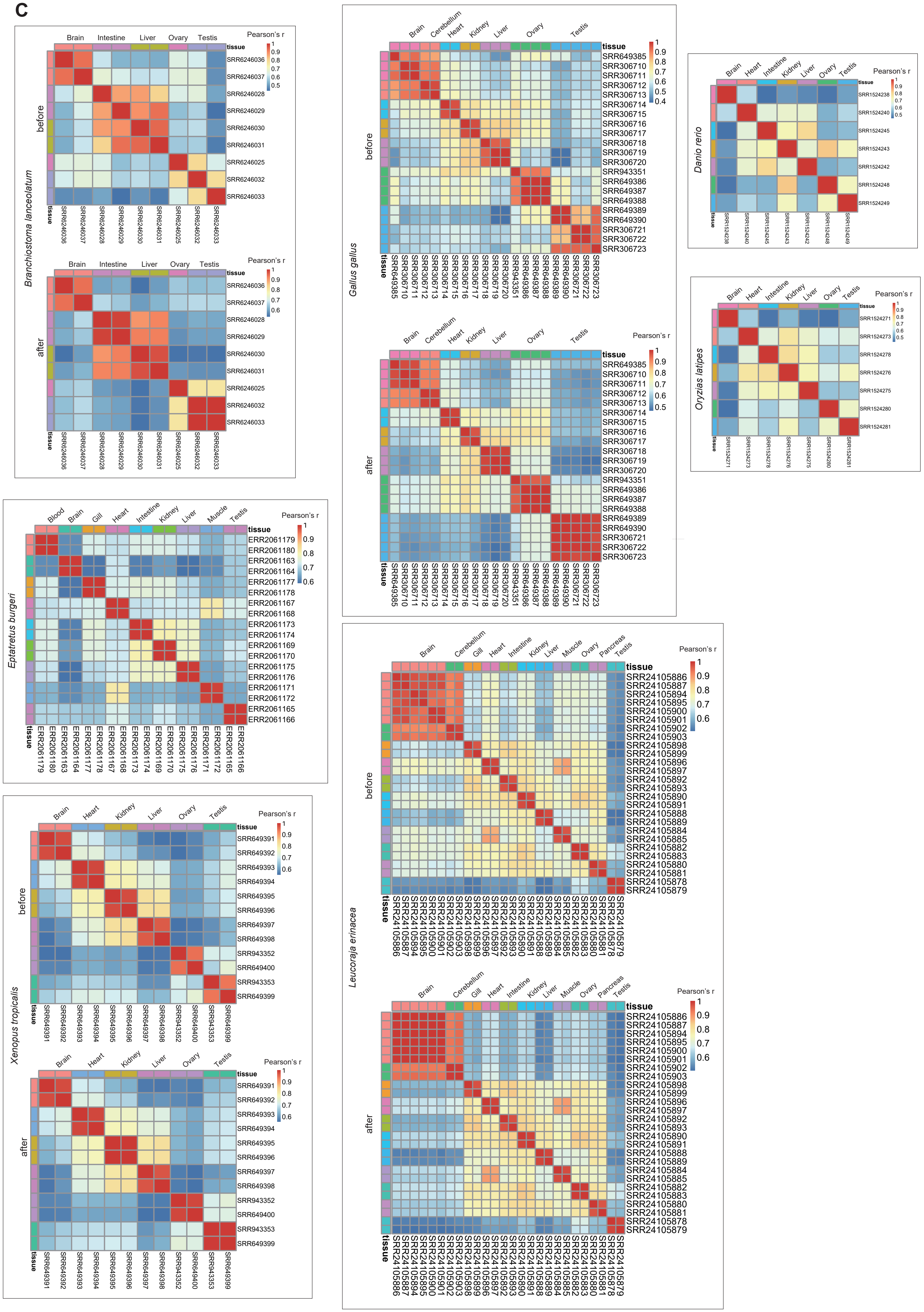
C
Branchiostoma lanceolatum
before
after
Brain Intestine Liver Ovary Testis
Pearson's r
tissue
SRR6246036 SRR6246037 SRR6246028 SRR6246029 SRR6246030 SRR6246031 SRR6246025 SRR6246032 SRR6246033
Gallus gallus
Brain Cerebellum Heart Kidney Liver Ovary Testis
SRR649385 SRR306710 SRR306711 SRR306712 SRR306713 SRR306714 SRR306715 SRR306716 SRR306717 SRR306718 SRR306719 SRR306720 SRR943351 SRR649386 SRR649387 SRR649388 SRR649389 SRR649390 SRR306721 SRR306722 SRR306723
Danio rerio
Brain Heart Intestine Kidney Liver Ovary Testis
SRR1524238 SRR1524240 SRR1524245 SRR1524243 SRR1524242 SRR1524248 SRR1524249
Oryzias latipes
SRR1524271 SRR1524273 SRR1524278 SRR1524276 SRR1524275 SRR1524280 SRR1524281
Eptatretus burgeri
Blood Brain Gill Heart Intestine Kidney Liver Muscle Testis
ERR2061179 ERR2061180 ERR2061163 ERR2061164 ERR2061177 ERR2061178 ERR2061167 ERR2061168 ERR2061173 ERR2061174 ERR2061169 ERR2061170 ERR2061175 ERR2061176 ERR2061171 ERR2061172 ERR2061165 ERR2061166
Xenopus tropicalis
Brain Heart Kidney Liver Ovary Testis
SRR649391 SRR649392 SRR649393 SRR649394 SRR649395 SRR649396 SRR649397 SRR649398 SRR943352 SRR649400 SRR943353 SRR649399
Leucoraja erinacea
Brain Cerebellum Gill Heart Intestine Kidney Liver Muscle Ovary Pancreas Testis
SRR24105886 SRR24105887 SRR24105894 SRR24105895 SRR24105900 SRR24105901 SRR24105902 SRR24105903 SRR24105898 SRR24105899 SRR24105896 SRR24105897 SRR24105892 SRR24105893 SRR24105890 SRR24105891 SRR24105888 SRR24105889 SRR24105884 SRR24105885 SRR24105882 SRR24105883 SRR24105880 SRR24105881 SRR24105878 SRR24105879

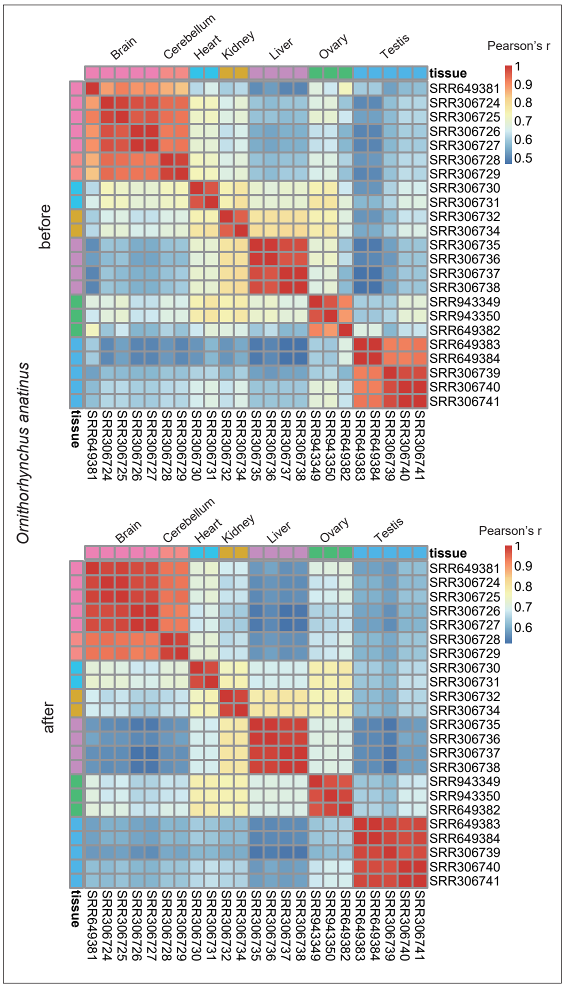
Ornithorhynchus anatinus
before
after
Brain
Cerebellum
Heart
Kidney
Liver
Ovary
Testis
tissue
Pearson's r
1
0.9
0.8
0.7
0.6
0.5
SRR649381
SRR306724
SRR306725
SRR306726
SRR306727
SRR306728
SRR306729
SRR306730
SRR306731
SRR306732
SRR306734
SRR306735
SRR306736
SRR306737
SRR306738
SRR943349
SRR943350
SRR649382
SRR649383
SRR649384
SRR306739
SRR306740
SRR306741

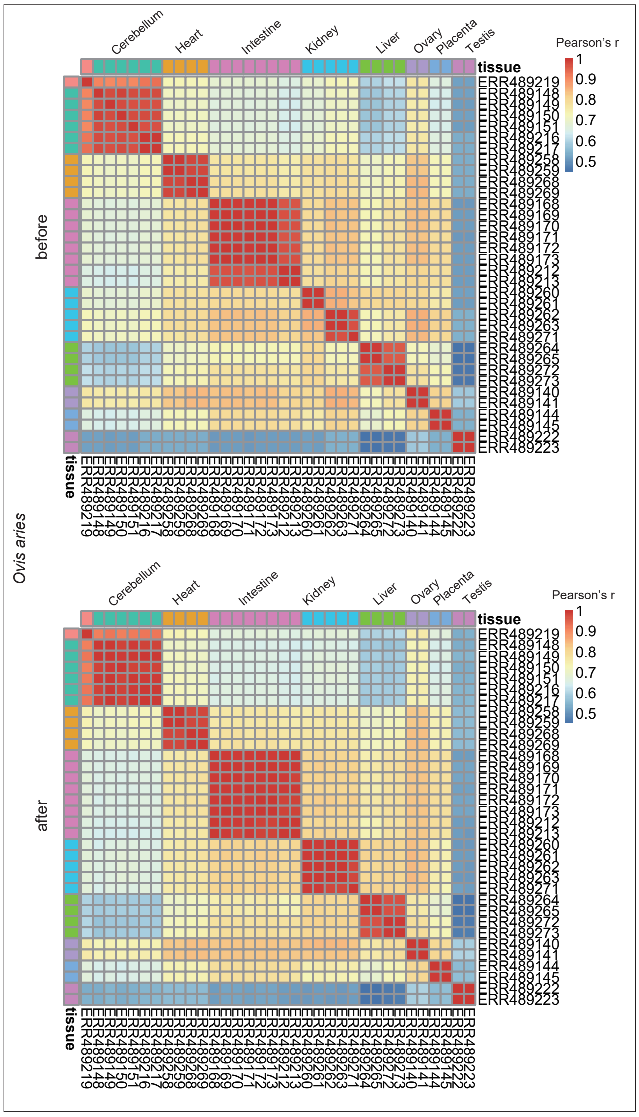
Ovis aries
before
after
Cerebellum
Heart
Intestine
Kidney
Liver
Ovary
Placenta
Testis
tissue
Pearson's r
1
0.9
0.8
0.7
0.6
0.5

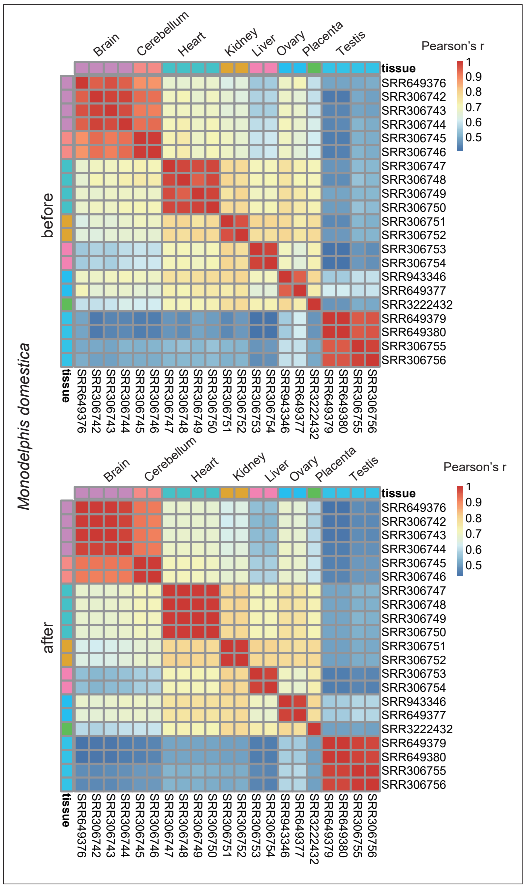
Monodelphis domestica
before
after
Brain
Cerebellum
Heart
Kidney
Liver
Ovary
Placenta
Testis
tissue
Pearson's r
1
0.9
0.8
0.7
0.6
0.5
SRR649376
SRR306742
SRR306743
SRR306744
SRR306745
SRR306746
SRR306747
SRR306748
SRR306749
SRR306750
SRR306751
SRR306752
SRR306753
SRR306754
SRR943346
SRR649377
SRR3222432
SRR649379
SRR649380
SRR306755
SRR306756

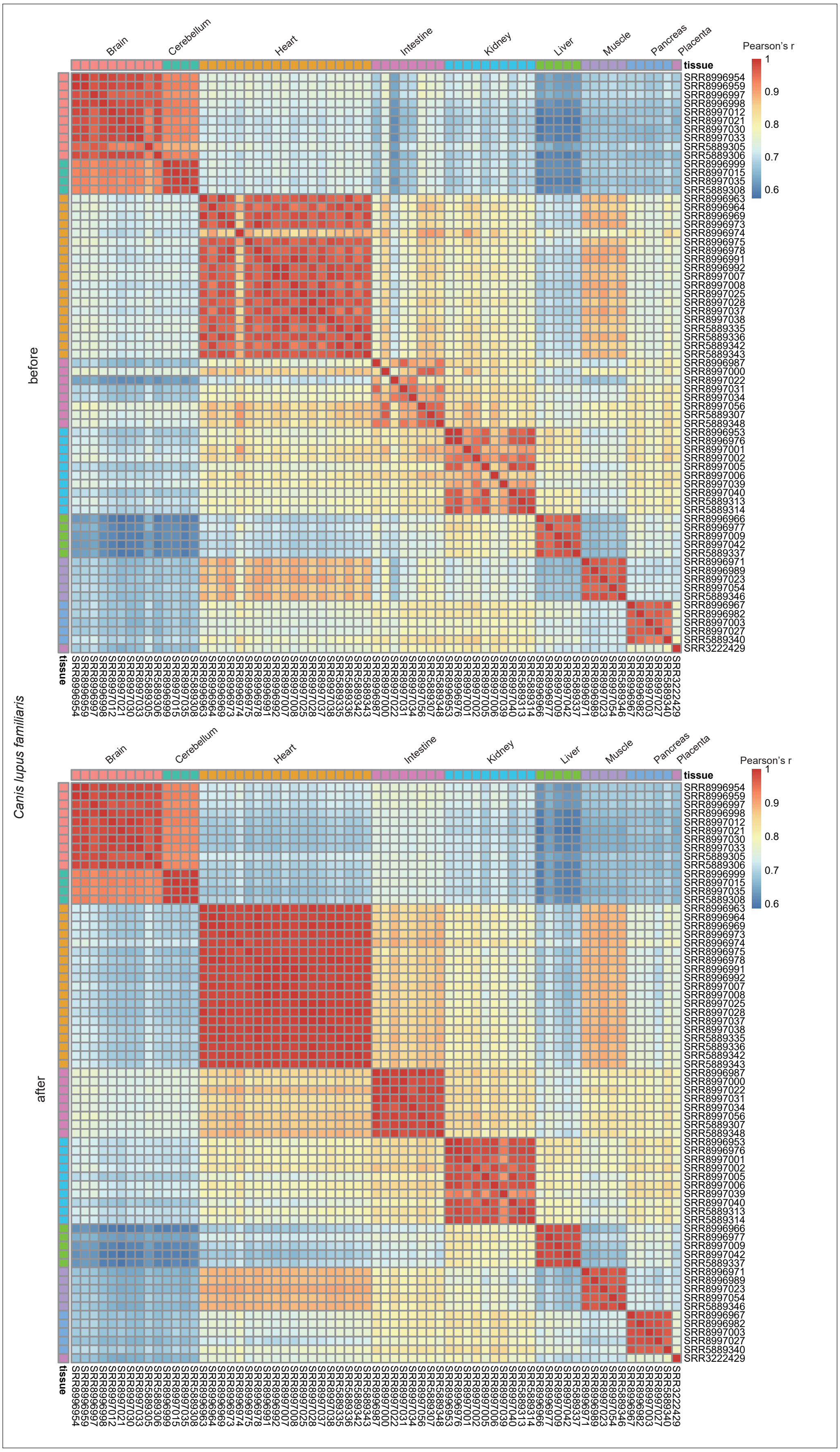

Canis lupus familiaris
before
after
Brain
Cerebellum
Heart
Intestine
Kidney
Liver
Muscle
Pancreas
Placenta
tissue
Pearson's r
1
0.9
0.8
0.7
0.6
SRR8996954
SRR8996959
SRR8996997
SRR8996998
SRR8997012
SRR8997021
SRR8997030
SRR8997033
SRR5889305
SRR5889306
SRR8996999
SRR8997015
SRR8997035
SRR5889308
SRR8996963
SRR8996964
SRR8996969
SRR8996973
SRR8996974
SRR8996975
SRR8996978
SRR8996991
SRR8996992
SRR8997007
SRR8997008
SRR8997025
SRR8997028
SRR8997037
SRR8997038
SRR5889335
SRR5889336
SRR5889342
SRR5889343
SRR8996987
SRR8997000
SRR8997022
SRR8997031
SRR8997034
SRR8997056
SRR5889307
SRR5889348
SRR8996953
SRR8996976
SRR8997001
SRR8997002
SRR8997005
SRR8997006
SRR8997039
SRR8997040
SRR5889313
SRR5889314
SRR8996966
SRR8996977
SRR8997009
SRR8997042
SRR5889337
SRR8996971
SRR8996989
SRR8997023
SRR8997054
SRR5889346
SRR8996967
SRR8996982
SRR8997003
SRR8997027
SRR5889340
SRR3222429

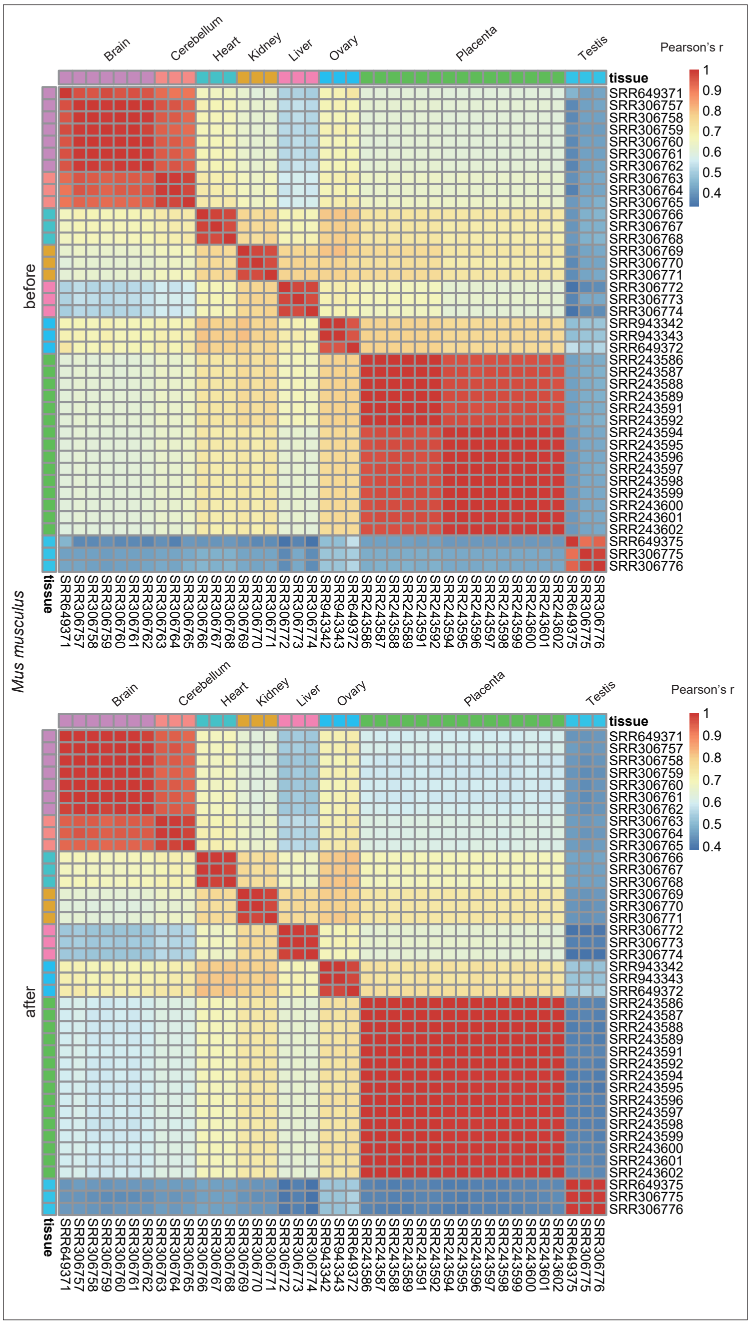
Mus musculus
before
after
Brain
Cerebellum
Heart
Kidney
Liver
Ovary
Placenta
Testis
tissue
Pearson's r
1
0.9
0.8
0.7
0.6
0.5
0.4

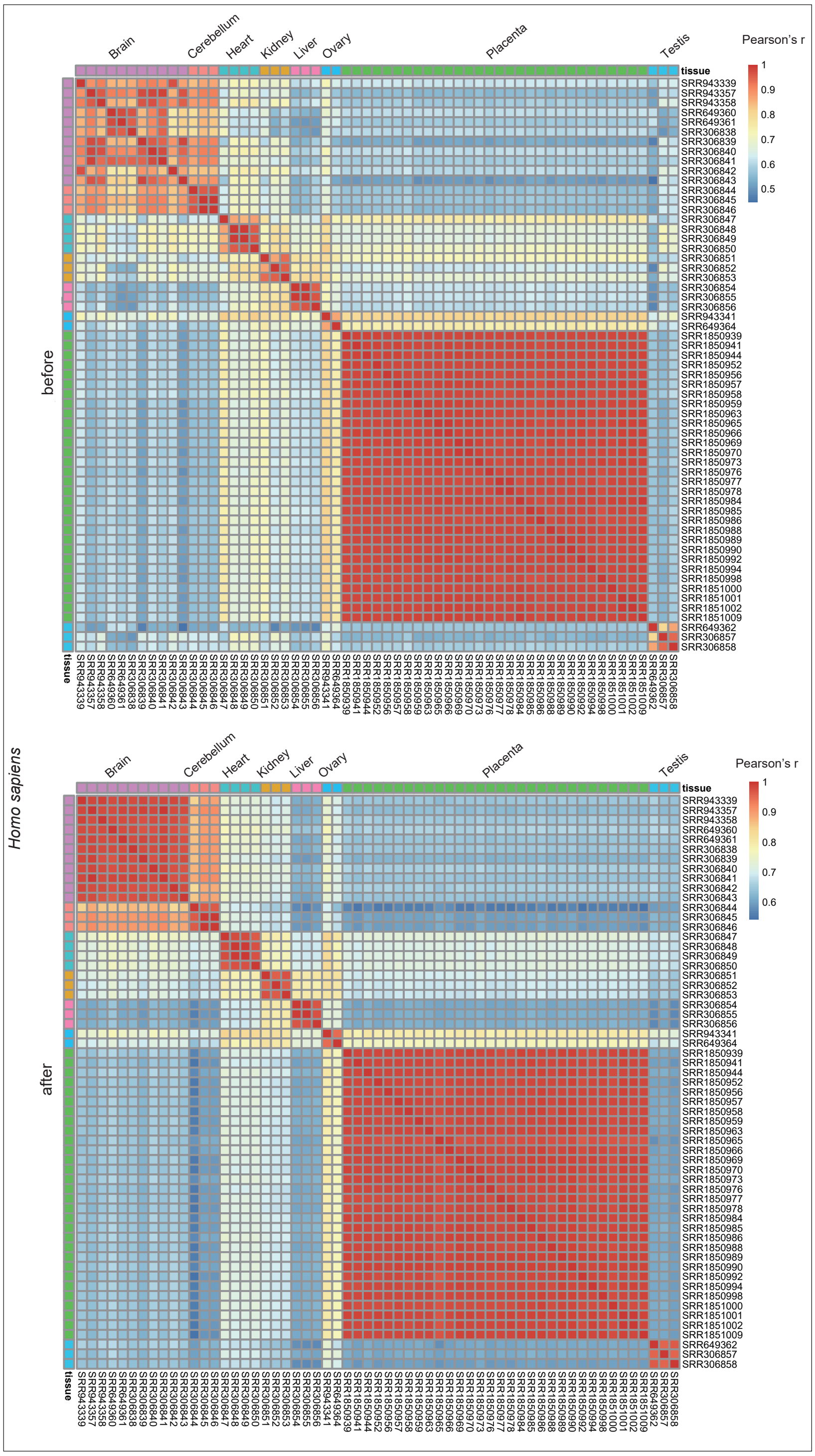
Homo sapiens
before
after
Brain
Cerebellum
Heart
Kidney
Liver
Ovary
Placenta
Testis
tissue
Pearson's r

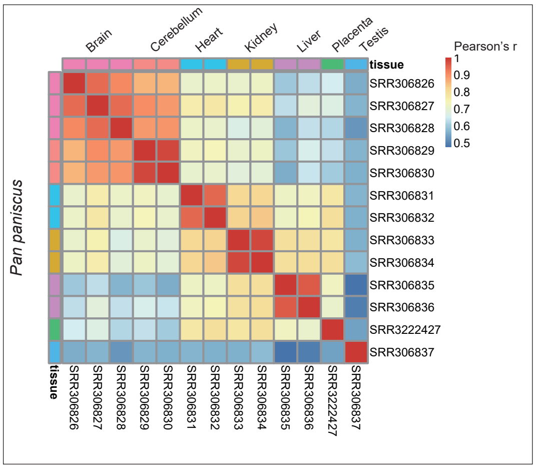
Pan paniscus
Brain
Cerebellum
Heart
Kidney
Liver
Placenta
Testis
tissue
Pearson's r
SRR306826
SRR306827
SRR306828
SRR306829
SRR306830
SRR306831
SRR306832
SRR306833
SRR306834
SRR306835
SRR306836
SRR3222427
SRR306837

### Figure S3

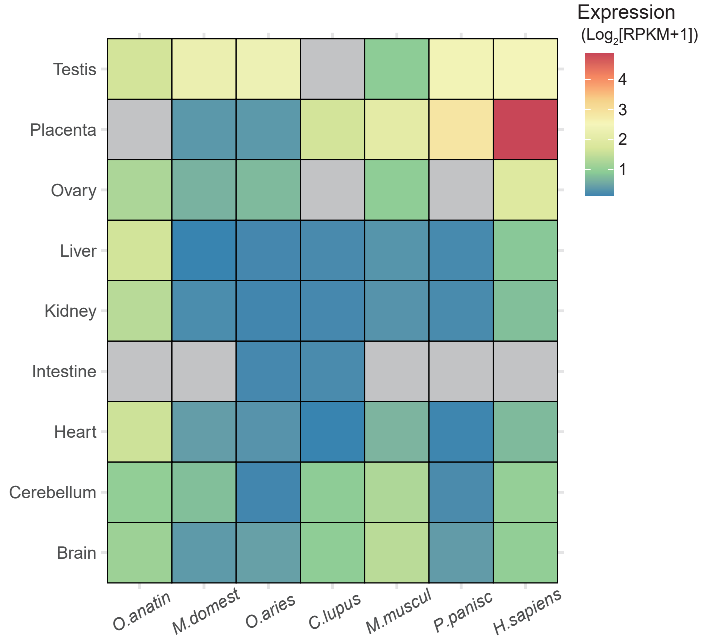

**Figure S3**. *ATG9B* has placenta- or testis-enriched expression in mammals. Color indicates the expression levels; gray specifically means no data were available. *O. anatin*, *Ornithorhynchus anatinus*; *M. domest*, *Monodelphis domestica*; *O. aries*, *Ovis aries*; *C. lupus*, *Canis lupus familiaris*; *M. muscul*, *Mus musculus*; *P. panisc*, *Pan paniscus*; *H. sapiens*, *Homo sapiens*.

Figure S4

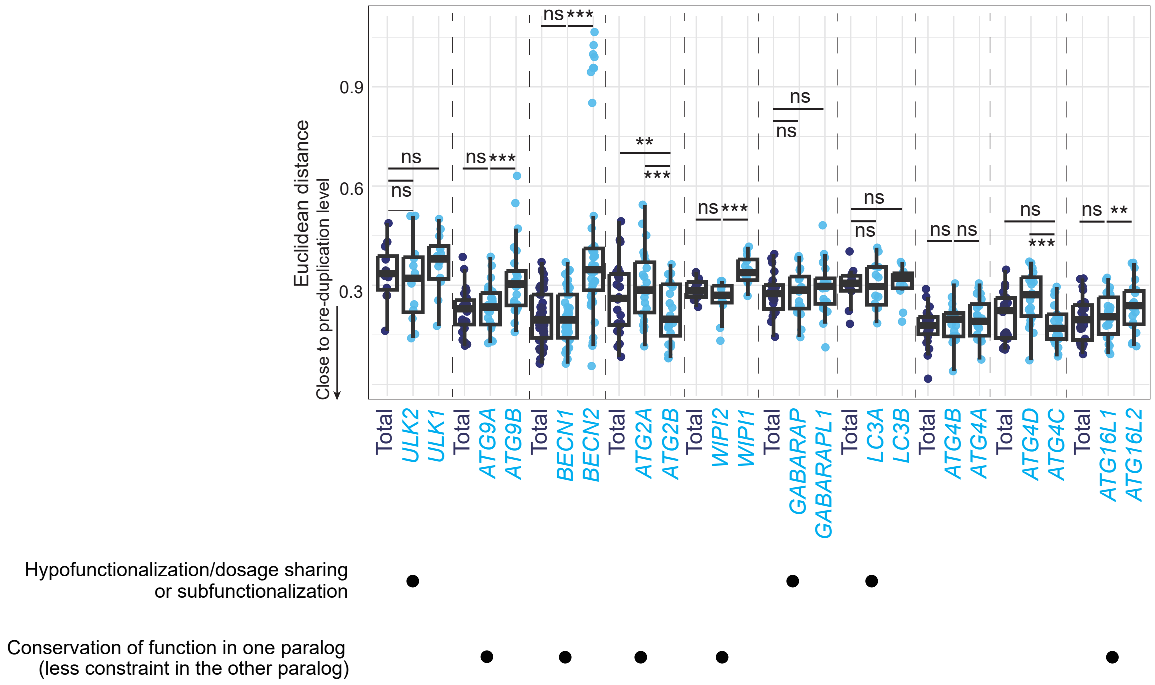

**Figure S4**. Evolutionary fate classification of the *ATG* paralogs based on the relative gene expression levels (gene expression in each tissue normalized to the total expression across all tissues), with associated evolutionary fate categories shown below. A generalized linear mixed model was used to compare the differences in Euclidean distances. *, $p < 0.05$. **, $p < 0.01$. ***, $p < 0.001$. ns, non-significant ($p > 0.05$).

**Table S1**. List of 30 Chordata species and five outgroup species used in the homology search and protein sequence evolution analysis.

| Taxonomy group | Species | Common name | Genome id |
|---|---|---|---|
| Cephalochordata | *Branchiostoma lanceolatum* | European lancelet | GCA_927797965.1 |
| Cephalochordata | *Branchiostoma floridae* | Florida lancelet | GCF_000003815.2 |
| Urochordata | *Ciona intestinalis* | Sea squirt | GCF_000224145.3 |
| Cyclostomata | *Eptatretus burgeri* | Inshore hagfish | GCA_900186335.2 |
| Cyclostomata | *Eptatretus atami* | Brown hagfish | Marlétaz et al. 2024 |
| Cyclostomata | *Petromyzon marinus* | Sea lamprey | GCF_010993605.1 |
| Cyclostomata | *Lethenteron reissneri* | Far Eastern brook lamprey | GCF_015708825.1 |
| Cartilaginous fish | *Callorhinchus milii* | Australian ghostshark | GCF_018977255.1 |
| Cartilaginous fish | *Carcharodon carcharias* | Great white shark | GCF_017639515.1 |
| Cartilaginous fish | *Leucoraja erinacea* | Little skate | GCF_028641065.1 |
| Ray-finned fish | *Danio rerio* | Zebrafish | GCF_000002035.6 |
| Ray-finned fish | *Salmo salar* | Atlantic salmon | GCF_905237065.1 |
| Ray-finned fish | *Oryzias latipes* | Medaka | GCF_002234675.1 |
| Ray-finned fish | *Astyanax mexicanus* | Mexican tetra | GCF_023375975.1 |
| Ray-finned fish | *Oreochromis niloticus* | Nile tilapia | GCF_001858045.2 |
| Lobe-finned fish | *Latimeria chalumnae* | West Indian Ocean coelacanth | GCF_000225785.1 |
| Amphibia | *Xenopus tropicalis* | Western clawed frog | GCF_000004195.4 |
| Amphibia | *Xenopus laevis* | African clawed frog | GCF_017654675.1 |
| Sauropsida | *Gallus gallus* | Chicken | GCF_016699485.2 |
| Sauropsida | *Anolis carolinensis* | Green anole | GCF_000090745.1 |
| Sauropsida | *Taeniopygia guttata* | Sunda zebra finch | GCF_003957565.2 |
| Sauropsida | *Gopherus evgoodei* | Goode's thornscrub tortoise | GCF_007399415.2 |
| Sauropsida | *Melopsittacus undulatus* | Common parakeet | GCF_012275295.1 |
| Mammalia | *Ornithorhynchus anatinus* | Platypus | GCF_004115215.2 |
| Mammalia | *Monodelphis domestica* | Opossum | GCF_027887165.1 |
| Mammalia | *Canis lupus familiaris* | Dog | GCF_000002285.5 |
| Mammalia | *Ovis aries* | Sheep | GCF_016772045.2 |
| Mammalia | *Mus musculus* | Mouse | GCF_000001635.27 |
| Mammalia | *Pan paniscus* | Bonobo | GCF_029289425.1 |
| Mammalia | *Homo sapiens* | Human | GCF_000001405.40 |
| Outgroup | *Trichoplax adhaerens* | | GCF_000150275.1 |
| Outgroup | *Mizuhopecten yessoensis* | Yesso scallop | GCF_002113885.1 |

| | | | |
|---|---|---|---|
| Outgroup | *Strongylocentrotus purpuratus* | Purple sea urchin | GCF_000002235.5 |
| Outgroup | *Drosophila melanogaster* | | GCF_000001215.4 |
| Outgroup | *Caenorhabditis elegans* | | GCF_000002985.6 |

**Table S2**. Ohnolog status based on three publications.

| | If ohnolog from Vance and McLysaght 2023 | | |
|---|---|---|---|
| ATG_gene | Makino and McLysaght 2010 | Singh and Isambert 2020 | Nakatani 2021 (analyzed by Vance and McLysaght 2023) |
| *ULK1* | T | T | F |
| *ULK2* | | | |
| *ATG9A* | T | T | F |
| *ATG9B* | | | |
| *BECN1* | F | F | F |
| *BECN2* | | | |
| *ATG2A* | T | T | F |
| *ATG2B* | | | |
| *WIPI1* | T | T | F |
| *WIPI2* | | | |
| *GABARAP* | T | F | F |
| *GABARAPL1* | | | |
| *LC3A* | T | T | F |
| *LC3B* | | | |
| *ATG4A* | F | T | F |
| *ATG4B* | | | |
| *ATG4C* | T | T | F |
| *ATG4D* | | | |
| *ATG16L1* | T | T | F |
| *ATG16L2* | | | |

**Table S3**. List of RNA-seq samples in the 14 species used in the gene expression evolution analysis.

| Species | Genome id | SRA id | Tissue name | Tissue category | Citation of data source |
|---|---|---|---|---|---|
| *Branchiostoma lanceolatum* | GCA_927797965.1 | SRR6246036 | Neural tube | Brain | Marletaz et al. 2018 |
| *Branchiostoma lanceolatum* | GCA_927797965.1 | SRR6246037 | Neural tube | Brain | |
| *Branchiostoma lanceolatum* | GCA_927797965.1 | SRR6246028 | Gut | Intestine | |
| *Branchiostoma lanceolatum* | GCA_927797965.1 | SRR6246029 | Gut | Intestine | |
| *Branchiostoma lanceolatum* | GCA_927797965.1 | SRR6246030 | Hepatic diverticulum | Liver | |
| *Branchiostoma lanceolatum* | GCA_927797965.1 | SRR6246031 | Hepatic diverticulum | Liver | |
| *Branchiostoma lanceolatum* | GCA_927797965.1 | SRR6246025 | Female gonads | Ovary | |
| *Branchiostoma lanceolatum* | GCA_927797965.1 | SRR6246032 | Male gonads | Testis | |
| *Branchiostoma lanceolatum* | GCA_927797965.1 | SRR6246033 | Male gonads | Testis | |
| *Eptatretus burgeri* | GCA_900186335.2 | ERR2061179 | Blood | Blood | Yu et al. 2023 |
| *Eptatretus burgeri* | GCA_900186335.2 | ERR2061180 | Blood | Blood | |
| *Eptatretus burgeri* | GCA_900186335.2 | ERR2061163 | Brain | Brain | |
| *Eptatretus burgeri* | GCA_900186335.2 | ERR2061164 | Brain | Brain | |
| *Eptatretus burgeri* | GCA_900186335.2 | ERR2061177 | Gill | Gill | |
| *Eptatretus burgeri* | GCA_900186335.2 | ERR2061178 | Gill | Gill | |
| *Eptatretus burgeri* | GCA_900186335.2 | ERR2061167 | Heart | Heart | |
| *Eptatretus burgeri* | GCA_900186335.2 | ERR2061168 | Heart | Heart | |
| *Eptatretus burgeri* | GCA_900186335.2 | ERR2061173 | Intestine | Intestine | |
| *Eptatretus burgeri* | GCA_900186335.2 | ERR2061174 | Intestine | Intestine | |
| *Eptatretus burgeri* | GCA_900186335.2 | ERR2061169 | Kidney | Kidney | |
| *Eptatretus burgeri* | GCA_900186335.2 | ERR2061170 | Kidney | Kidney | |
| *Eptatretus burgeri* | GCA_900186335.2 | ERR2061175 | Liver | Liver | |
| *Eptatretus burgeri* | GCA_900186335.2 | ERR2061176 | Liver | Liver | |
| *Eptatretus burgeri* | GCA_900186335.2 | ERR2061171 | Skeletal muscle | Muscle | |
| *Eptatretus burgeri* | GCA_900186335.2 | ERR2061172 | Skeletal muscle | Muscle | |
| *Eptatretus burgeri* | GCA_900186335.2 | ERR2061165 | Testis | Testis | |
| *Eptatretus burgeri* | GCA_900186335.2 | ERR2061166 | Testis | Testis | |

| | | | | | |
|---|---|---|---|---|---|
| *Leucoraja erinacea* | GCF_028641065.1 | SRR24105886 | Midbrain | Brain | Marletaz et al. 2023 |
| *Leucoraja erinacea* | GCF_028641065.1 | SRR24105887 | Midbrain | Brain | |
| *Leucoraja erinacea* | GCF_028641065.1 | SRR24105894 | Hindbrain | Brain | |
| *Leucoraja erinacea* | GCF_028641065.1 | SRR24105895 | Hindbrain | Brain | |
| *Leucoraja erinacea* | GCF_028641065.1 | SRR24105900 | Forebrain | Brain | |
| *Leucoraja erinacea* | GCF_028641065.1 | SRR24105901 | Forebrain | Brain | |
| *Leucoraja erinacea* | GCF_028641065.1 | SRR24105902 | Cerebellum | Cerebellum | |
| *Leucoraja erinacea* | GCF_028641065.1 | SRR24105903 | Cerebellum | Cerebellum | |
| *Leucoraja erinacea* | GCF_028641065.1 | SRR24105898 | Gill | Gill | |
| *Leucoraja erinacea* | GCF_028641065.1 | SRR24105899 | Gill | Gill | |
| *Leucoraja erinacea* | GCF_028641065.1 | SRR24105896 | Heart | Heart | |
| *Leucoraja erinacea* | GCF_028641065.1 | SRR24105897 | Heart | Heart | |
| *Leucoraja erinacea* | GCF_028641065.1 | SRR24105892 | Intestine | Intestine | |
| *Leucoraja erinacea* | GCF_028641065.1 | SRR24105893 | Intestine | Intestine | |
| *Leucoraja erinacea* | GCF_028641065.1 | SRR24105890 | Kidney | Kidney | |
| *Leucoraja erinacea* | GCF_028641065.1 | SRR24105891 | Kidney | Kidney | |
| *Leucoraja erinacea* | GCF_028641065.1 | SRR24105888 | Liver | Liver | |
| *Leucoraja erinacea* | GCF_028641065.1 | SRR24105889 | Liver | Liver | |
| *Leucoraja erinacea* | GCF_028641065.1 | SRR24105884 | Muscle | Muscle | |
| *Leucoraja erinacea* | GCF_028641065.1 | SRR24105885 | Muscle | Muscle | |
| *Leucoraja erinacea* | GCF_028641065.1 | SRR24105882 | Ovary | Ovary | |
| *Leucoraja erinacea* | GCF_028641065.1 | SRR24105883 | Ovary | Ovary | |
| *Leucoraja erinacea* | GCF_028641065.1 | SRR24105880 | Pancreas | Pancreas | |
| *Leucoraja erinacea* | GCF_028641065.1 | SRR24105881 | Pancreas | Pancreas | |
| *Leucoraja erinacea* | GCF_028641065.1 | SRR24105878 | Testis | Testis | |
| *Leucoraja erinacea* | GCF_028641065.1 | SRR24105879 | Testis | Testis | |
| *Danio rerio* | GCF_000002035.6 | SRR1524238 | Brain | Brain | Pasquier et al. 2016 |
| *Danio rerio* | GCF_000002035.6 | SRR1524240 | Heart | Heart | |

| | | | | | |
|---|---|---|---|---|---|
| *Danio rerio* | GCF_000002035.6 | SRR1524245 | Intestine | Intestine | |
| *Danio rerio* | GCF_000002035.6 | SRR1524243 | Kidney | Kidney | |
| *Danio rerio* | GCF_000002035.6 | SRR1524242 | Liver | Liver | |
| *Danio rerio* | GCF_000002035.6 | SRR1524248 | Ovary | Ovary | |
| *Danio rerio* | GCF_000002035.6 | SRR1524249 | Testis | Testis | |
| *Oryzias latipes* | GCF_002234675.1 | SRR1524271 | Brain | Brain | |
| *Oryzias latipes* | GCF_002234675.1 | SRR1524273 | Heart | Heart | |
| *Oryzias latipes* | GCF_002234675.1 | SRR1524278 | Intestine | Intestine | |
| *Oryzias latipes* | GCF_002234675.1 | SRR1524276 | Kidney | Kidney | |
| *Oryzias latipes* | GCF_002234675.1 | SRR1524275 | Liver | Liver | |
| *Oryzias latipes* | GCF_002234675.1 | SRR1524280 | Ovary | Ovary | |
| *Oryzias latipes* | GCF_002234675.1 | SRR1524281 | Testis | Testis | |
| *Xenopus tropicalis* | GCF_000004195.4 | SRR649391 | Brain | Brain | Necsulea et al. 2014 |
| *Xenopus tropicalis* | GCF_000004195.4 | SRR649392 | Brain | Brain | |
| *Xenopus tropicalis* | GCF_000004195.4 | SRR649393 | Heart | Heart | |
| *Xenopus tropicalis* | GCF_000004195.4 | SRR649394 | Heart | Heart | |
| *Xenopus tropicalis* | GCF_000004195.4 | SRR649395 | Kidney | Kidney | |
| *Xenopus tropicalis* | GCF_000004195.4 | SRR649396 | Kidney | Kidney | |
| *Xenopus tropicalis* | GCF_000004195.4 | SRR649397 | Liver | Liver | |
| *Xenopus tropicalis* | GCF_000004195.4 | SRR649398 | Liver | Liver | |
| *Xenopus tropicalis* | GCF_000004195.4 | SRR943352 | Ovary | Ovary | |
| *Xenopus tropicalis* | GCF_000004195.4 | SRR649400 | Ovary | Ovary | |
| *Xenopus tropicalis* | GCF_000004195.4 | SRR943353 | Testis | Testis | |
| *Xenopus tropicalis* | GCF_000004195.4 | SRR649399 | Testis | Testis | |
| *Gallus gallus* | GCF_016699485.2 | SRR649385 | Brain | Brain | Brawand et al. 2011, Necsulea et al. 2014 |
| *Gallus gallus* | GCF_016699485.2 | SRR306710 | Brain | Brain | |
| *Gallus gallus* | GCF_016699485.2 | SRR306711 | Brain | Brain | |
| *Gallus gallus* | GCF_016699485.2 | SRR306712 | Cerebellum | Cerebellum | |
| *Gallus gallus* | GCF_016699485.2 | SRR306713 | Cerebellum | Cerebellum | |
| *Gallus gallus* | GCF_016699485.2 | SRR306714 | Heart | Heart | |
| *Gallus gallus* | GCF_016699485.2 | SRR306715 | Heart | Heart | |

| | | | | |
|---|---|---|---|---|
| *Gallus gallus* | GCF_016699485.2 | SRR306716 | Kidney | Kidney |
| *Gallus gallus* | GCF_016699485.2 | SRR306717 | Kidney | Kidney |
| *Gallus gallus* | GCF_016699485.2 | SRR306718 | Liver | Liver |
| *Gallus gallus* | GCF_016699485.2 | SRR306719 | Liver | Liver |
| *Gallus gallus* | GCF_016699485.2 | SRR306720 | Liver | Liver |
| *Gallus gallus* | GCF_016699485.2 | SRR943351 | Ovary | Ovary |
| *Gallus gallus* | GCF_016699485.2 | SRR649386 | Ovary | Ovary |
| *Gallus gallus* | GCF_016699485.2 | SRR649387 | Ovary | Ovary |
| *Gallus gallus* | GCF_016699485.2 | SRR649388 | Ovary | Ovary |
| *Gallus gallus* | GCF_016699485.2 | SRR649389 | Testis | Testis |
| *Gallus gallus* | GCF_016699485.2 | SRR649390 | Testis | Testis |
| *Gallus gallus* | GCF_016699485.2 | SRR306721 | Testis | Testis |
| *Gallus gallus* | GCF_016699485.2 | SRR306722 | Testis | Testis |
| *Gallus gallus* | GCF_016699485.2 | SRR306723 | Testis | Testis |
| *Ornithorhynchus anatinus* | GCF_004115215.2 | SRR649381 | Brain | Brain |
| *Ornithorhynchus anatinus* | GCF_004115215.2 | SRR306724 | Brain | Brain |
| *Ornithorhynchus anatinus* | GCF_004115215.2 | SRR306725 | Brain | Brain |
| *Ornithorhynchus anatinus* | GCF_004115215.2 | SRR306726 | Brain | Brain |
| *Ornithorhynchus anatinus* | GCF_004115215.2 | SRR306727 | Brain | Brain |
| *Ornithorhynchus anatinus* | GCF_004115215.2 | SRR306728 | Cerebellum | Cerebellum |
| *Ornithorhynchus anatinus* | GCF_004115215.2 | SRR306729 | Cerebellum | Cerebellum |
| *Ornithorhynchus anatinus* | GCF_004115215.2 | SRR306730 | Heart | Heart |
| *Ornithorhynchus anatinus* | GCF_004115215.2 | SRR306731 | Heart | Heart |
| *Ornithorhynchus anatinus* | GCF_004115215.2 | SRR306732 | Kidney | Kidney |
| *Ornithorhynchus anatinus* | GCF_004115215.2 | SRR306734 | Kidney | Kidney |
| *Ornithorhynchus anatinus* | GCF_004115215.2 | SRR306735 | Liver | Liver |
| *Ornithorhynchus anatinus* | GCF_004115215.2 | SRR306736 | Liver | Liver |
| *Ornithorhynchus anatinus* | GCF_004115215.2 | SRR306737 | Liver | Liver |
| *Ornithorhynchus anatinus* | GCF_004115215.2 | SRR306738 | Liver | Liver |
| *Ornithorhynchus anatinus* | GCF_004115215.2 | SRR943349 | Ovary | Ovary |

| | | | | | |
|---|---|---|---|---|---|
| *Ornithorhynchus anatinus* | GCF_004115215.2 | SRR943350 | Ovary | Ovary | |
| *Ornithorhynchus anatinus* | GCF_004115215.2 | SRR649382 | Ovary | Ovary | |
| *Ornithorhynchus anatinus* | GCF_004115215.2 | SRR649383 | Testis | Testis | |
| *Ornithorhynchus anatinus* | GCF_004115215.2 | SRR649384 | Testis | Testis | |
| *Ornithorhynchus anatinus* | GCF_004115215.2 | SRR306739 | Testis | Testis | |
| *Ornithorhynchus anatinus* | GCF_004115215.2 | SRR306740 | Testis | Testis | |
| *Ornithorhynchus anatinus* | GCF_004115215.2 | SRR306741 | Testis | Testis | |
| *Monodelphis domestica* | GCF_027887165.1 | SRR649376 | Brain | Brain | Brawand et al. 2011, Necsulea et al. 2014, Armstrong et al. 2017 |
| *Monodelphis domestica* | GCF_027887165.1 | SRR306742 | Brain | Brain | |
| *Monodelphis domestica* | GCF_027887165.1 | SRR306743 | Brain | Brain | |
| *Monodelphis domestica* | GCF_027887165.1 | SRR306744 | Brain | Brain | |
| *Monodelphis domestica* | GCF_027887165.1 | SRR306745 | Cerebellum | Cerebellu m | |
| *Monodelphis domestica* | GCF_027887165.1 | SRR306746 | Cerebellum | Cerebellu m | |
| *Monodelphis domestica* | GCF_027887165.1 | SRR306747 | Heart | Heart | |
| *Monodelphis domestica* | GCF_027887165.1 | SRR306748 | Heart | Heart | |
| *Monodelphis domestica* | GCF_027887165.1 | SRR306749 | Heart | Heart | |
| *Monodelphis domestica* | GCF_027887165.1 | SRR306750 | Heart | Heart | |
| *Monodelphis domestica* | GCF_027887165.1 | SRR306751 | Kidney | Kidney | |
| *Monodelphis domestica* | GCF_027887165.1 | SRR306752 | Kidney | Kidney | |
| *Monodelphis domestica* | GCF_027887165.1 | SRR306753 | Liver | Liver | |
| *Monodelphis domestica* | GCF_027887165.1 | SRR306754 | Liver | Liver | |
| *Monodelphis domestica* | GCF_027887165.1 | SRR943346 | Ovary | Ovary | |
| *Monodelphis domestica* | GCF_027887165.1 | SRR649377 | Ovary | Ovary | |
| *Monodelphis domestica* | GCF_027887165.1 | SRR3222432 | Placenta | Placenta | |
| *Monodelphis domestica* | GCF_027887165.1 | SRR649379 | Testis | Testis | |
| *Monodelphis domestica* | GCF_027887165.1 | SRR649380 | Testis | Testis | |
| *Monodelphis domestica* | GCF_027887165.1 | SRR306755 | Testis | Testis | |

| | | | | | |
|---|---|---|---|---|---|
| *Monodelphis domestica* | GCF_027887165.1 | SRR306756 | Testis | Testis | |
| *Ovis aries* | GCF_016772045.2 | ERR489148 | Cerebellum | Cerebellum | University of Edinburgh sheep gene expression atlas, Archibald et al. 2010 |
| *Ovis aries* | GCF_016772045.2 | ERR489149 | Cerebellum | Cerebellum | |
| *Ovis aries* | GCF_016772045.2 | ERR489150 | Cerebellum | Cerebellum | |
| *Ovis aries* | GCF_016772045.2 | ERR489151 | Cerebellum | Cerebellum | |
| *Ovis aries* | GCF_016772045.2 | ERR489216 | Brain cerebellum | Cerebellum | |
| *Ovis aries* | GCF_016772045.2 | ERR489217 | Brain cerebellum | Cerebellum | |
| *Ovis aries* | GCF_016772045.2 | ERR489219 | Brain cerebrum | Brain | |
| *Ovis aries* | GCF_016772045.2 | ERR489258 | Ventricle | Heart | |
| *Ovis aries* | GCF_016772045.2 | ERR489259 | Ventricle | Heart | |
| *Ovis aries* | GCF_016772045.2 | ERR489268 | Heart vertricle | Heart | |
| *Ovis aries* | GCF_016772045.2 | ERR489269 | Heart vertricle | Heart | |
| *Ovis aries* | GCF_016772045.2 | ERR489168 | Colon | Intestine | |
| *Ovis aries* | GCF_016772045.2 | ERR489169 | Colon | Intestine | |
| *Ovis aries* | GCF_016772045.2 | ERR489170 | Colon | Intestine | |
| *Ovis aries* | GCF_016772045.2 | ERR489171 | Colon | Intestine | |
| *Ovis aries* | GCF_016772045.2 | ERR489172 | Colon | Intestine | |
| *Ovis aries* | GCF_016772045.2 | ERR489173 | Colon | Intestine | |
| *Ovis aries* | GCF_016772045.2 | ERR489212 | Colon | Intestine | |
| *Ovis aries* | GCF_016772045.2 | ERR489213 | Colon | Intestine | |
| *Ovis aries* | GCF_016772045.2 | ERR489260 | Kidney cortex | Kidney | |
| *Ovis aries* | GCF_016772045.2 | ERR489261 | Kidney cortex | Kidney | |
| *Ovis aries* | GCF_016772045.2 | ERR489262 | Kidney medulla | Kidney | |
| *Ovis aries* | GCF_016772045.2 | ERR489263 | Kidney medulla | Kidney | |
| *Ovis aries* | GCF_016772045.2 | ERR489271 | Kidney medulla | Kidney | |
| *Ovis aries* | GCF_016772045.2 | ERR489264 | Liver | Liver | |
| *Ovis aries* | GCF_016772045.2 | ERR489265 | Liver | Liver | |
| *Ovis aries* | GCF_016772045.2 | ERR489272 | Liver | Liver | |
| *Ovis aries* | GCF_016772045.2 | ERR489273 | Liver | Liver | |
| *Ovis aries* | GCF_016772045.2 | ERR489140 | Ovary | Ovary | |
| *Ovis aries* | GCF_016772045.2 | ERR489141 | Ovary | Ovary | |
| *Ovis aries* | GCF_016772045.2 | ERR489144 | Placenta | Placenta | |

| | | | | | |
|---|---|---|---|---|---|
| *Ovis aries* | GCF_016772045.2 | ERR489145 | Placenta | Placenta | |
| *Ovis aries* | GCF_016772045.2 | ERR489222 | Testes | Testis | |
| *Ovis aries* | GCF_016772045.2 | ERR489223 | Testes | Testis | |
| *Canis lupus familiaris* | GCF_000002285.5 | SRR8996954 | Occipital cortex | Brain | Broad Institute, Armstrong et al. 2017 |
| *Canis lupus familiaris* | GCF_000002285.5 | SRR8996959 | Frontal cortex | Brain | |
| *Canis lupus familiaris* | GCF_000002285.5 | SRR8996997 | Frontal cortex | Brain | |
| *Canis lupus familiaris* | GCF_000002285.5 | SRR8996998 | Occipital cortex | Brain | |
| *Canis lupus familiaris* | GCF_000002285.5 | SRR8997012 | Occipital cortex | Brain | |
| *Canis lupus familiaris* | GCF_000002285.5 | SRR8997021 | Frontal cortex | Brain | |
| *Canis lupus familiaris* | GCF_000002285.5 | SRR8997030 | Occipital cortex | Brain | |
| *Canis lupus familiaris* | GCF_000002285.5 | SRR8997033 | Frontal cortex | Brain | |
| *Canis lupus familiaris* | GCF_000002285.5 | SRR5889305 | Frontal cortex | Brain | |
| *Canis lupus familiaris* | GCF_000002285.5 | SRR5889306 | Occipital cortex | Brain | |
| *Canis lupus familiaris* | GCF_000002285.5 | SRR8996999 | Cerebellum | Cerebellum | |
| *Canis lupus familiaris* | GCF_000002285.5 | SRR8997015 | Cerebellum | Cerebellum | |
| *Canis lupus familiaris* | GCF_000002285.5 | SRR8997035 | Cerebellum | Cerebellum | |
| *Canis lupus familiaris* | GCF_000002285.5 | SRR5889308 | Cerebellum | Cerebellum | |
| *Canis lupus familiaris* | GCF_000002285.5 | SRR8996963 | Left ventricle | Heart | |
| *Canis lupus familiaris* | GCF_000002285.5 | SRR8996964 | Left atrium | Heart | |
| *Canis lupus familiaris* | GCF_000002285.5 | SRR8996969 | Right ventricle | Heart | |
| *Canis lupus familiaris* | GCF_000002285.5 | SRR8996973 | Right ventricle | Heart | |
| *Canis lupus familiaris* | GCF_000002285.5 | SRR8996974 | Right atrium | Heart | |
| *Canis lupus familiaris* | GCF_000002285.5 | SRR8996975 | Left atrium | Heart | |
| *Canis lupus familiaris* | GCF_000002285.5 | SRR8996978 | Left ventricle | Heart | |
| *Canis lupus familiaris* | GCF_000002285.5 | SRR8996991 | Right ventricle | Heart | |
| *Canis lupus familiaris* | GCF_000002285.5 | SRR8996992 | Right atrium | Heart | |
| *Canis lupus familiaris* | GCF_000002285.5 | SRR8997007 | Left atrium | Heart | |
| *Canis lupus familiaris* | GCF_000002285.5 | SRR8997008 | Left ventricle | Heart | |

| | | | | |
|---|---|---|---|---|
| *Canis lupus familiaris* | GCF_000002285.5 | SRR8997025 | Right ventricle | Heart |
| *Canis lupus familiaris* | GCF_000002285.5 | SRR8997028 | Right atrium | Heart |
| *Canis lupus familiaris* | GCF_000002285.5 | SRR8997037 | Left ventricle | Heart |
| *Canis lupus familiaris* | GCF_000002285.5 | SRR8997038 | Left atrium | Heart |
| *Canis lupus familiaris* | GCF_000002285.5 | SRR5889335 | Left atrium | Heart |
| *Canis lupus familiaris* | GCF_000002285.5 | SRR5889336 | Left ventricle | Heart |
| *Canis lupus familiaris* | GCF_000002285.5 | SRR5889342 | Right atrium | Heart |
| *Canis lupus familiaris* | GCF_000002285.5 | SRR5889343 | Right ventricle | Heart |
| *Canis lupus familiaris* | GCF_000002285.5 | SRR8996987 | Small intestine | Intestine |
| *Canis lupus familiaris* | GCF_000002285.5 | SRR8997000 | Colon | Intestine |
| *Canis lupus familiaris* | GCF_000002285.5 | SRR8997022 | Colon | Intestine |
| *Canis lupus familiaris* | GCF_000002285.5 | SRR8997031 | Small intestine | Intestine |
| *Canis lupus familiaris* | GCF_000002285.5 | SRR8997034 | Colon | Intestine |
| *Canis lupus familiaris* | GCF_000002285.5 | SRR8997056 | Small intestine | Intestine |
| *Canis lupus familiaris* | GCF_000002285.5 | SRR5889307 | Colon | Intestine |
| *Canis lupus familiaris* | GCF_000002285.5 | SRR5889348 | Small intestine | Intestine |
| *Canis lupus familiaris* | GCF_000002285.5 | SRR8996953 | Kidney cortex | Kidney |
| *Canis lupus familiaris* | GCF_000002285.5 | SRR8996976 | Kidney medulla | Kidney |
| *Canis lupus familiaris* | GCF_000002285.5 | SRR8997001 | Kidney cortex | Kidney |
| *Canis lupus familiaris* | GCF_000002285.5 | SRR8997002 | Kidney medulla | Kidney |
| *Canis lupus familiaris* | GCF_000002285.5 | SRR8997005 | Kidney cortex | Kidney |
| *Canis lupus familiaris* | GCF_000002285.5 | SRR8997006 | Kidney medulla | Kidney |
| *Canis lupus familiaris* | GCF_000002285.5 | SRR8997039 | Kidney medulla | Kidney |
| *Canis lupus familiaris* | GCF_000002285.5 | SRR8997040 | Kidney cortex | Kidney |
| *Canis lupus familiaris* | GCF_000002285.5 | SRR5889313 | Kidney medulla | Kidney |
| *Canis lupus familiaris* | GCF_000002285.5 | SRR5889314 | Kidney cortex | Kidney |
| *Canis lupus familiaris* | GCF_000002285.5 | SRR8996966 | Liver | Liver |

| | | | | | |
|---|---|---|---|---|---|
| *Canis lupus familiaris* | GCF_000002285.5 | SRR8996977 | Liver | Liver | |
| *Canis lupus familiaris* | GCF_000002285.5 | SRR8997009 | Liver | Liver | |
| *Canis lupus familiaris* | GCF_000002285.5 | SRR8997042 | Liver | Liver | |
| *Canis lupus familiaris* | GCF_000002285.5 | SRR5889337 | Liver | Liver | |
| *Canis lupus familiaris* | GCF_000002285.5 | SRR8996971 | Skeletal muscle | Muscle | |
| *Canis lupus familiaris* | GCF_000002285.5 | SRR8996989 | Skeletal muscle | Muscle | |
| *Canis lupus familiaris* | GCF_000002285.5 | SRR8997023 | Skeletal muscle | Muscle | |
| *Canis lupus familiaris* | GCF_000002285.5 | SRR8997054 | Skeletal muscle | Muscle | |
| *Canis lupus familiaris* | GCF_000002285.5 | SRR5889346 | Skeletal muscle | Muscle | |
| *Canis lupus familiaris* | GCF_000002285.5 | SRR8996967 | Pancreas | Pancreas | |
| *Canis lupus familiaris* | GCF_000002285.5 | SRR8996982 | Pancreas | Pancreas | |
| *Canis lupus familiaris* | GCF_000002285.5 | SRR8997003 | Pancreas | Pancreas | |
| *Canis lupus familiaris* | GCF_000002285.5 | SRR8997027 | Pancreas | Pancreas | |
| *Canis lupus familiaris* | GCF_000002285.5 | SRR5889340 | Pancreas | Pancreas | |
| *Canis lupus familiaris* | GCF_000002285.5 | SRR3222429 | Placenta | Placenta | |
| *Mus musculus* | GCF_000001635.27 | SRR649371 | Brain | Brain | Brawand et al. 2011, Necsulea et al. 2014, Wang et al. 2011 |
| *Mus musculus* | GCF_000001635.27 | SRR306757 | Brain | Brain | |
| *Mus musculus* | GCF_000001635.27 | SRR306758 | Brain | Brain | |
| *Mus musculus* | GCF_000001635.27 | SRR306759 | Brain | Brain | |
| *Mus musculus* | GCF_000001635.27 | SRR306760 | Brain | Brain | |
| *Mus musculus* | GCF_000001635.27 | SRR306761 | Brain | Brain | |
| *Mus musculus* | GCF_000001635.27 | SRR306762 | Brain | Brain | |
| *Mus musculus* | GCF_000001635.27 | SRR306763 | Cerebellum | Cerebellum | |
| *Mus musculus* | GCF_000001635.27 | SRR306764 | Cerebellum | Cerebellum | |
| *Mus musculus* | GCF_000001635.27 | SRR306765 | Cerebellum | Cerebellum | |
| *Mus musculus* | GCF_000001635.27 | SRR306766 | Heart | Heart | |
| *Mus musculus* | GCF_000001635.27 | SRR306767 | Heart | Heart | |

| | | | | |
|---|---|---|---|---|
| *Mus musculus* | GCF_000001635.27 | SRR306768 | Heart | Heart |
| *Mus musculus* | GCF_000001635.27 | SRR306769 | Kidney | Kidney |
| *Mus musculus* | GCF_000001635.27 | SRR306770 | Kidney | Kidney |
| *Mus musculus* | GCF_000001635.27 | SRR306771 | Kidney | Kidney |
| *Mus musculus* | GCF_000001635.27 | SRR306772 | Liver | Liver |
| *Mus musculus* | GCF_000001635.27 | SRR306773 | Liver | Liver |
| *Mus musculus* | GCF_000001635.27 | SRR306774 | Liver | Liver |
| *Mus musculus* | GCF_000001635.27 | SRR943342 | Ovary | Ovary |
| *Mus musculus* | GCF_000001635.27 | SRR943343 | Ovary | Ovary |
| *Mus musculus* | GCF_000001635.27 | SRR649372 | Ovary | Ovary |
| *Mus musculus* | GCF_000001635.27 | SRR243586 | Placenta | Placenta |
| *Mus musculus* | GCF_000001635.27 | SRR243587 | Placenta | Placenta |
| *Mus musculus* | GCF_000001635.27 | SRR243588 | Placenta | Placenta |
| *Mus musculus* | GCF_000001635.27 | SRR243589 | Placenta | Placenta |
| *Mus musculus* | GCF_000001635.27 | SRR243591 | Placenta | Placenta |
| *Mus musculus* | GCF_000001635.27 | SRR243592 | Placenta | Placenta |
| *Mus musculus* | GCF_000001635.27 | SRR243594 | Placenta | Placenta |
| *Mus musculus* | GCF_000001635.27 | SRR243595 | Placenta | Placenta |
| *Mus musculus* | GCF_000001635.27 | SRR243596 | Placenta | Placenta |
| *Mus musculus* | GCF_000001635.27 | SRR243597 | Placenta | Placenta |
| *Mus musculus* | GCF_000001635.27 | SRR243598 | Placenta | Placenta |
| *Mus musculus* | GCF_000001635.27 | SRR243599 | Placenta | Placenta |
| *Mus musculus* | GCF_000001635.27 | SRR243600 | Placenta | Placenta |
| *Mus musculus* | GCF_000001635.27 | SRR243601 | Placenta | Placenta |
| *Mus musculus* | GCF_000001635.27 | SRR243602 | Placenta | Placenta |
| *Mus musculus* | GCF_000001635.27 | SRR649375 | Testis | Testis |
| *Mus musculus* | GCF_000001635.27 | SRR306775 | Testis | Testis |

| | | | | | |
|---|---|---|---|---|---|
| *Mus musculus* | GCF_000001635.27 | SRR306776 | Testis | Testis | |
| *Pan paniscus* | GCF_029289425.1 | SRR306826 | Brain prefrontal cortex | Brain | Brawand et al. 2011, Necsulea et al. 2014, Armstrong et al. 2017 |
| *Pan paniscus* | GCF_029289425.1 | SRR306827 | Brain prefrontal cortex | Brain | |
| *Pan paniscus* | GCF_029289425.1 | SRR306828 | Brain prefrontal cortex | Brain | |
| *Pan paniscus* | GCF_029289425.1 | SRR306829 | Cerebellum | Cerebellum | |
| *Pan paniscus* | GCF_029289425.1 | SRR306830 | Cerebellum | Cerebellum | |
| *Pan paniscus* | GCF_029289425.1 | SRR306831 | Heart | Heart | |
| *Pan paniscus* | GCF_029289425.1 | SRR306832 | Heart | Heart | |
| *Pan paniscus* | GCF_029289425.1 | SRR306833 | Kidney | Kidney | |
| *Pan paniscus* | GCF_029289425.1 | SRR306834 | Kidney | Kidney | |
| *Pan paniscus* | GCF_029289425.1 | SRR306835 | Liver | Liver | |
| *Pan paniscus* | GCF_029289425.1 | SRR306836 | Liver | Liver | |
| *Pan paniscus* | GCF_029289425.1 | SRR3222427 | Placenta | Placenta | |
| *Pan paniscus* | GCF_029289425.1 | SRR306837 | Testis | Testis | |
| *Homo sapiens* | GCF_000001405.40 | SRR943339 | Brain frontal cortex | Brain | Brawand et al. 2011, Necsulea et al. 2014, Hughes et al. 2015 |
| *Homo sapiens* | GCF_000001405.40 | SRR943357 | Brain frontal cortex | Brain | |
| *Homo sapiens* | GCF_000001405.40 | SRR943358 | Brain frontal cortex | Brain | |
| *Homo sapiens* | GCF_000001405.40 | SRR649360 | Brain frontal cortex | Brain | |
| *Homo sapiens* | GCF_000001405.40 | SRR649361 | Brain frontal cortex | Brain | |
| *Homo sapiens* | GCF_000001405.40 | SRR306838 | Brain frontal cortex | Brain | |
| *Homo sapiens* | GCF_000001405.40 | SRR306839 | Brain frontal cortex | Brain | |
| *Homo sapiens* | GCF_000001405.40 | SRR306840 | Brain prefrontal cortex | Brain | |
| *Homo sapiens* | GCF_000001405.40 | SRR306841 | Brain prefrontal cortex | Brain | |
| *Homo sapiens* | GCF_000001405.40 | SRR306842 | Brain prefrontal cortex | Brain | |
| *Homo sapiens* | GCF_000001405.40 | SRR306843 | Brain temporal lobe | Brain | |
| *Homo sapiens* | GCF_000001405.40 | SRR306844 | Cerebellum | Cerebellum | |

| | | | | |
|---|---|---|---|---|
| *Homo sapiens* | GCF_000001405.40 | SRR306845 | Cerebellum | Cerebellum |
| *Homo sapiens* | GCF_000001405.40 | SRR306846 | Cerebellum | Cerebellum |
| *Homo sapiens* | GCF_000001405.40 | SRR306847 | Heart | Heart |
| *Homo sapiens* | GCF_000001405.40 | SRR306848 | Heart | Heart |
| *Homo sapiens* | GCF_000001405.40 | SRR306849 | Heart | Heart |
| *Homo sapiens* | GCF_000001405.40 | SRR306850 | Heart | Heart |
| *Homo sapiens* | GCF_000001405.40 | SRR306851 | Kidney | Kidney |
| *Homo sapiens* | GCF_000001405.40 | SRR306852 | Kidney | Kidney |
| *Homo sapiens* | GCF_000001405.40 | SRR306853 | Kidney | Kidney |
| *Homo sapiens* | GCF_000001405.40 | SRR306854 | Liver | Liver |
| *Homo sapiens* | GCF_000001405.40 | SRR306855 | Liver | Liver |
| *Homo sapiens* | GCF_000001405.40 | SRR306856 | Liver | Liver |
| *Homo sapiens* | GCF_000001405.40 | SRR943341 | Ovary | Ovary |
| *Homo sapiens* | GCF_000001405.40 | SRR649364 | Ovary | Ovary |
| *Homo sapiens* | GCF_000001405.40 | SRR1850939 | placenta | Placenta |
| *Homo sapiens* | GCF_000001405.40 | SRR1850941 | placenta | Placenta |
| *Homo sapiens* | GCF_000001405.40 | SRR1850944 | placenta | Placenta |
| *Homo sapiens* | GCF_000001405.40 | SRR1850952 | placenta | Placenta |
| *Homo sapiens* | GCF_000001405.40 | SRR1850956 | placenta | Placenta |
| *Homo sapiens* | GCF_000001405.40 | SRR1850957 | placenta | Placenta |
| *Homo sapiens* | GCF_000001405.40 | SRR1850958 | placenta | Placenta |
| *Homo sapiens* | GCF_000001405.40 | SRR1850959 | placenta | Placenta |
| *Homo sapiens* | GCF_000001405.40 | SRR1850963 | placenta | Placenta |
| *Homo sapiens* | GCF_000001405.40 | SRR1850965 | placenta | Placenta |
| *Homo sapiens* | GCF_000001405.40 | SRR1850966 | placenta | Placenta |
| *Homo sapiens* | GCF_000001405.40 | SRR1850969 | placenta | Placenta |
| *Homo sapiens* | GCF_000001405.40 | SRR1850970 | placenta | Placenta |

| | | | | |
|---|---|---|---|---|
| *Homo sapiens* | GCF_000001405.40 | SRR1850973 | placenta | Placenta |
| *Homo sapiens* | GCF_000001405.40 | SRR1850976 | placenta | Placenta |
| *Homo sapiens* | GCF_000001405.40 | SRR1850977 | placenta | Placenta |
| *Homo sapiens* | GCF_000001405.40 | SRR1850978 | placenta | Placenta |
| *Homo sapiens* | GCF_000001405.40 | SRR1850984 | placenta | Placenta |
| *Homo sapiens* | GCF_000001405.40 | SRR1850985 | placenta | Placenta |
| *Homo sapiens* | GCF_000001405.40 | SRR1850986 | placenta | Placenta |
| *Homo sapiens* | GCF_000001405.40 | SRR1850988 | placenta | Placenta |
| *Homo sapiens* | GCF_000001405.40 | SRR1850989 | placenta | Placenta |
| *Homo sapiens* | GCF_000001405.40 | SRR1850990 | placenta | Placenta |
| *Homo sapiens* | GCF_000001405.40 | SRR1850992 | placenta | Placenta |
| *Homo sapiens* | GCF_000001405.40 | SRR1850994 | placenta | Placenta |
| *Homo sapiens* | GCF_000001405.40 | SRR1850998 | placenta | Placenta |
| *Homo sapiens* | GCF_000001405.40 | SRR1851000 | placenta | Placenta |
| *Homo sapiens* | GCF_000001405.40 | SRR1851001 | placenta | Placenta |
| *Homo sapiens* | GCF_000001405.40 | SRR1851002 | placenta | Placenta |
| *Homo sapiens* | GCF_000001405.40 | SRR1851009 | placenta | Placenta |
| *Homo sapiens* | GCF_000001405.40 | SRR649362 | Testis | Testis |
| *Homo sapiens* | GCF_000001405.40 | SRR306857 | Testis | Testis |
| *Homo sapiens* | GCF_000001405.40 | SRR306858 | Testis | Testis |
